# A tough nutlet to crack: resolving the phylogeny of *Thesium* (Thesiaceae), the largest genus in Santalales

**DOI:** 10.1101/2023.04.12.536538

**Authors:** Miguel A. García, Ladislav Mucina, Daniel L. Nickrent

## Abstract

With over 350 species, *Thesium* is the largest genus in Santalales. It is found on all continents except Antarctica; however, its highest diversity is in the Cape Floristic Region of South Africa where approximately half the species occur. *Thesium* samples of ca. 590 collections from throughout its entire geographic range were obtained and nuclear ribosomal ITS sequenced from 396 accessions representing 196 named taxa and 30 currently unnamed taxa for a total of ca. 230 species. In addition, two chloroplast genome spacers (*trnDT* and *trnLF*) were sequenced from 269 and 315 accessions, respectively. Maximum parsimony, maximum likelihood and Bayesian methods were employed to generate gene trees and infer phylogenies. The value of the morphological characters traditionally used in the taxonomy of the genus and previous infrageneric classifications are discussed. Broad scale relationships were generally congruent among the ITS and the chloroplast trees. For example, both the nuclear and chloroplast trees support the presence of Eurasian and African clades. In contrast, major incongruence was detected between nuclear and chloroplast trees for a number of taxa including the recently described *T. nautimontanum* that is sister to the entire African clade on the ITS tree. Although the causes of this incongruence are currently unknown, a novel form of chloroplast capture is hypothesized. A hypothesis of the biogeographical history of the genus based on our molecular phylogeny is presented.

## INTRODUCTION

Within the order Santalales, evolutionary radiations generating 100 species, or more, have occurred among several mistletoe genera such as *Amyema* Tiegh. and *Psittacanthus* Mart. (both Loranthaceae) as well as *Dendrophthora* Eichl. and *Phoradendron* Nutt. (both Viscaceae); however, the greatest proliferation of species has been in the root parasite *Thesium* L. Placed by some in Santalaceae, the classification in Nickrent & al. (2010) utilized molecular phylogenetic as well as morphological evidence to support its classification in a distinct family, Thesiaceae. Generic limits within that family were further defined by Nickrent & García (2015) such that now the family contains *Buckleya* Torr. (Asia and eastern North America), *Lacomucinaea* Nickrent & M.A.García (southern Africa), *Osyridicarpos* A.DC. (Africa), and *Thesium* (Africa, Europe, Asia, Australia and South America, Fig. 1). As shown in Nickrent & García (2015), four previously named genera are now contained within a broadly defined *Thesium*: *Austroamericium* Hendrych, *Chrysothesium* (Jaub. & Spach) Hendrych, *Kunkeliella* Stearn, and *Thesidium* Sond. Although present on all continents except Antarctica, *Thesium* is most diverse in South Africa with successively fewer species in other areas. One species (*T. ramosum* Hayne) has been introduced into North America and is considered a noxious weed (Nickrent, 2016).

**Fig. 1.**
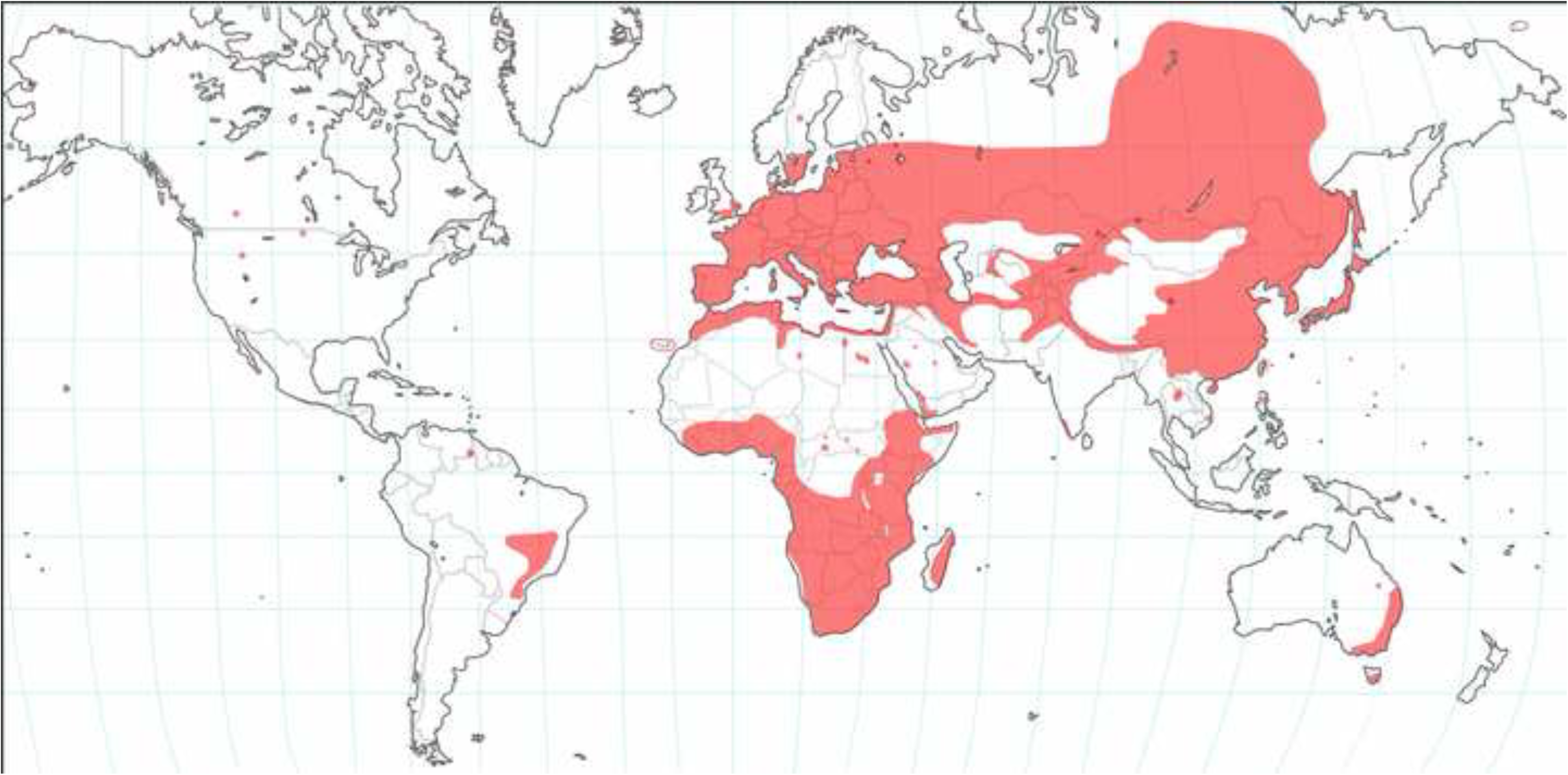
Map showing the distribution of *Thesium* worldwide. Data from GBIF were vetted and used for generating the first iteration. Additional information was obtained from specimen records given in various floras.

*Thesium* species include hemiparasitic annuals and perennials whose growth forms include herbs, subshrubs, shrubs, and scandent shrubs. Most species are stoloniferous or rhizomatous, but in those adapted to fire prone regions of Southern and Tropical Africa, the shoots arise from a caudex, which indicates their post-disturbance resprouting strategy. The alternate leaves may have broad lamina or, in xerophytically adapted species, may be reduced to scales (thus the plants are squamate). In general, vegetative features appear quite plastic, contrasting with the somewhat conservative floral morphology. Plasticity is most apparent when comparing juvenile to adult forms. Inflorescences are extremely variable and include simple and compound cymes, panicles, corymbs, racemes, spikes and condensed systems resembling capitula. Vegetative and inflorescence “zones” are often not well differentiated where stem leaves grade into floral bracts, thus producing what Troll (1964) termed frondose inflorescences. In many cases the inflorescence is a compound heterothetic system, e.g. racemes of dichasia, panicles of cymes, etc. Some species with seemingly simple inflorescences show, through the sequence of flower development, that the system is evolutionarily reduced from a more complex type. An example of this can be seen in brachyblast (short shoot) members present in Tropical Africa where clusters of bracts at the inflorescence base (and sometimes additional lateral flowers) suggest reduction from a more complex branched system. Flowers are usually subtended by two bracteoles and this unit by a bract. In squamate *Thesium* species, floral bracts can grade gradually into vegetative scale leaves. *Thesium* inflorescences can also exhibit recaulescence (Weberling, 1989) where the position of axillary floral buds is shifted owing to the codevelopment (“stretching”) of the common basal portions of the pedicel and subtending bract. The resulting partial inflorescence shows the lower portion of the subtending bract fused to an elongated peduncle that is terminated by the bracteoles, pedicel (if present), and flower. Because this process generates inflorescences that are developmentally complex and thus difficult to describe morphologically, some authors do not distinguish pedicel from peduncle (e.g., Visser et al., 2018). Here we consider the pedicel to be the axis emerging above the bracteoles that joins the lower portion of the inferior ovary. In many *Thesium* species this pedicel is absent, hence the flower is sessile. In others the pedicel dilates into an elaiosome upon fruiting. The peduncular axis below the bracteoles may show no recaulescence or it may be more or less fused to the bract. Varying degrees of recaulescence on the same floral axis, i.e. proximal flowers showing less fusion between peduncle and bract distal flowers, is known as progressive recaulescence and can be seen in members of the *Australe* Clade 21.

*Thesium* flowers are generally 5-merous and monochlamydous, i.e. with one perianth whorl that is best interpreted as the corolla (Der & Nickrent, 2008). The calyx is either absent (completely fused with inferior ovary), highly reduced, or with clear lobes as is seen in *T. libericum* Hepper & Keay. In some species, an external gland occurs at the base of the petal lobe sinuses that occupies the same position as sepal lobe remnants in other species. When present, the floral hypanthium may be short as in campanulate or rotate flowers to quite long as seen in tubular flowers. The term perianth tube has been used by other authors, however, we prefer hypanthium because this structure is the result of adnation of calyx, corolla and androecium. We also refrain from using the term receptacle (or receptacular tube) because this has implications on the evolution of the inferior ovary that have not been confirmed. The valvate corolla lobes may be completely free to the junction with the hypanthium or may be connate for various lengths. The inside apex of the corolla lobes may bear a dense beard of trichomes, a feature traditionally used to define the African *Thesium* sect. *Barbata* A.W.Hill. In *Thesium* sect. *Frisea* (Rchb.) A.DC. (≡ *Annulata* A.W.Hill), a ring of short downwardly directed golden hairs is inserted at the base of the corolla tube at the level of departure of the filaments. Hairs opposite the stamens (post-staminal trichomes) that adhere to the dorsal part of the anthers may be present or absent (e.g. *Thesium* sects. *Frisea* and *Penicillata* A.W.Hill).

Stamens are antipetalous and the filaments may be free at the junction of the hypanthium and the corolla lobes or positioned more distally on individual corolla lobes. The filament lengths vary widely such that both included and exserted anthers can be seen in various species. The positional relationship between the anthers and stigma is also determined by style length, which varies from very short and conical to longer than the hypanthium and corolla lobes.

The fruit in *Thesium* is a nutlike achene whose surface has prominent venation or a fleshy drupe with a smooth surface (*T.* sect. *Kunkeliella*, *T. triflorum* L.f., *T. radicans* Hohen ex A.DC.). The withered corolla may remain attached to the apex of the mature fruit and is of various lengths depending upon the species. This remnant may be accrescent during fruit development. In many *Thesium* species the pedicel and base of the ovary enlarges upon fruiting forming an elaiosome. This structure is attractive to ants, which remove and disseminate the fruits. Such myrmecochory has been associated with short distance dispersal, but the wide occurrence of the genus (e.g. Madagascar and South America) suggests other long-distance mechanisms may also play an important role.

The history of the infrageneric classification of *Thesium* has been reviewed in several papers including Hill (1915a), Hendrych (1972), Moore & al. (2010), Nickrent & García (2015), and Zhigila & al. (2020). Most systematic studies in the genus have been focused on particular geographical regions. Taxonomic treatments considering the whole range of distribution date from the 1800s; modern treatments, taking into account the morphological diversity of the whole genus, are lacking. More recently the genus has attracted the attention of researchers resulting in the publication of several taxonomic revisions of species groups in South Africa (Mashego & Le Roux, 2018; Visser & al., 2018; Zhigila & al., 2019a, Lombard & al., 2021) and the description of new taxa (e.g., Romo & al., 2004; Nickrent & García, 2015; García & al., 2018; Zhigila & al., 2019b; Lombard & al., 2019; Zhigila & Muasya, 2022; Lombard & Le Roux, 2023). Moore & al. (2010), although with very limited sampling and a focus on the Cape species of South Africa, published the first dated molecular phylogeny of the genus in which *Thesium* was resolved as paraphyletic relative to *Thesidium* and *Austroamericium*. Nickrent & García (2015), using a sampling from the previous work, included critical taxa such as *T. lineatum* L.f., *T. mauritanicum* Batt., two species of *Kunkeliella*, three species of *Chrysothesium* and *Osyridicarpos*. This work resolved *T. lineatum* (now recognized as a new genus *Lacomucinaea*) as sister to *Osyridicarpos*, and placed the genera *Thesidium*, *Kunkeliella* and the polyphyletic *Chysothesium* within *Thesium*. Zhigila & al. (2020) increased the sampling of South African species and, based on molecular phylogenetic data, proposed dividing the genus into five subgenera, all of which are monophyletic except *T.* subgen. *Discothesium* (A.DC.) Zhigila & al. which is paraphyletic.

The phylogenies published to date are largely biased towards species in subgenera *Discothesium*, *Hagnothesium* (A.DC.) Zhigila & al. and *Frisea* Rchb. of the Cape Floristic Region (CFR), with a very poor sampling of species from Eurasia (*T.* subgen. *Thesium*) and the rest of Africa (*T.* subgen. *Psilothesium* (A.DC.) Zhigila & al.) that together represent more than half of the species diversity of the genus. The goals of our study are: to 1) present a well-sampled molecular phylogeny of the five subgenera of *Thesium* based on DNA sequences from one nuclear (nrDNA ITS) and two chloroplast regions (*trnLF* and *trnDT*); 2) identify well-supported clades and evaluate how the infrageneric classifications proposed by Hill (1915a, 1915b), Bobrov (1936) and Hendrych (1972) and the informal groups by Polhill (2005) and Hilliard (2006) relate to our phylogenetic results; 3) discuss in a phylogenetic framework the value of the morphological characters used in the most important taxonomic treatments of the genus across its entire geographical distribution; 4) suggest hypotheses of phylogenetic relationships of the species not sampled in our study, highlighting the taxonomic and phylogenetic work needed to achieve a natural infrageneric classification of *Thesium*; 5) elaborate a hypothesis of the biogeographical history of the genus based on our molecular phylogeny; and 6) discuss the topological incongurences found between nuclear and chloroplast phylogenies, especially in *T.* subgen. *Frisea*.

## MATERIALS AND METHODS

### Taxon sampling

Our sampling strategy within the speciose genus *Thesium* involved several operational facets. Our overall goal was to obtain as many species as possible across the worldwide distribution of the genus. We began with a list of approximately 600 *Thesium* names obtained from the International Plant Names Index (IPNI, 2023) and reduced this number by removing recognized synonyms. Some of these synonyms are indicated on the IPNI site whereas others were obtained from various regional floristic treatments of the genus. More recently, our list of accepted names and synonyms was compared to data present on World Plants (Hassler, 2004--2023). Direct examination of specimens showed that additional synonyms also likely exist; however, these were treated as separate operational taxa whose genetic relationships were to be determined in this and future studies. In the discussion of the clades resulting from the phylogenetic analyses, we added information on all unsampled species that, based on morphology, might be related to those included in particular clades in our analyses. The idea was to propose hypotheses of phylogenetic relationships of all the species of the genus that could be tested when additional material becomes available.

Tissue samples were obtained via two methods: silica gel dried shoots obtained from living plants and from herbarium specimens. Field work was conducted by the first author within Europe and Armenia (2005), second author in South Africa spanning a period 1996 and 2009, third author in South Africa (1996), and by all authors in South Africa (2007). Fresh leaf and stem samples were dried on silica gel and standard herbarium voucher specimens prepared. These specimens are deposited at MA, BH and STE (NBG) with some duplicates in BR. Several herbaria were visited (BR, LE, MO, NY, and UPS) and specimen loans obtained from others (BRLU, LG, NBG, NU, NH, and POZG) for morphological revision. When permitted, samples of *Thesium* tissue were removed from these herbarium specimens for DNA extraction. It should be noted that some herbaria that contain vast collections of South African *Thesium*, such as K, NBG and PRE, do not allow DNA extraction from their specimens. These restrictions were followed, thus many species from the Cape region of South Africa could not be sampled.

All specimens used in the molecular analyses were identified after reviewing type material directly or by viewing high-resolution images available from the JSTOR Global Plants web site (https://plants.jstor.org) and comparing with other herbarium specimens. Original protologues and descriptions from regional floras were also examined. These steps were crucial because a high percentage of herbarium specimens were misidentified, particularly from South Africa where a critical taxonomic revision of the genus is missing.

### Wet lab methods

Genomic DNA was obtained using two methods. The first was a standard 2X CTAB protocol (Nickrent, 1994; Nickrent & al., 2004) and the second used a cell disrupter with ceramic beads (BIO101/ThermoSavant FastPrep FP120). The protocol given in the DNeasy 96 Plant Kit (QIAGEN, Valencia, California, USA) was followed. The latter method proved to be particularly effective in yielding amplifiable DNA from herbarium specimens. PCR amplification of the ITS region was accomplished using the primers 18S 1830for (5’--AAC AAG GTT TCC GTA GGT GA--3’) and 26S 40rev (5’--TCC TCC GCT TAT TGA TAT GC--3’) with standard reaction mix and cycling conditions (Nickrent & al., 2004). The *trnDT* region of the chloroplast was amplified using the primers trnD^guc^ and trnT^ggu^ (Demesure & al., 1995) and the *trnLF* region using the primers published by Taberlet & al. (1991). The amplification products were cleaned using either the QIAquick PCR purification kit (QIAGEN, Valencia, California, USA) or the E.Z.N.A. Clean kit (Omega Biotech, Doraville, Georgia, USA). In-house cycle sequencing reactions were conducted in a GeneAmp 9700 thermocycler (Applied Biosystems, Foster City, California, USA) with the BigDye terminator Cycle Sequencing Ready Reaction Kit (Applied Biosystems), using the above ITS primers. Cycle sequencing reactions were purified using either an ethanol/sodium acetate precipitation method or with ExopSAP-IT^®^ (USB Corporation, Cleveland, Ohio, USA) generally following the manufacturer’s instructions (enzyme can be diluted 1:10).

Some sequencing was conducted using either an ABI Prism 377 DNA Sequencer or an AB 3730S capillary DNA analyzer (Applied Biosystems). Additional samples were sequenced by Macrogen Inc. (South Korea). If multiple bands were seen following the initial PCR reaction, the bands were excised from the gel and TA cloned using the pGEM- T easy-vector II cloning kit (Promega). The standard mini-prep step was avoided by using colony PCR. Here six white colonies from each plate were picked with sterile toothpicks and put directly into 0.2 ml PCR tubes each containing: 18.8 µl ddH2O, 2 µl MgCl2 (25µM), 2.5 µl Buffer 10X, 0.32 µl dNTPs (10µM), 0.2 µl Taq (5 units/µl), 0.625 µl T7 promoter primer (5µM), 0.625 µl SP6 primer (5µM). The PCR conditions were as follows: 94°C for 5 minutes followed by 35 cycles of 94°C for 1 minute, 50 °C for 1 minute and 72 °C for 1 minute, and a final extension at 72 °C for 20 minutes. The PCR products were then cleaned using ExopSAP-IT^®^. These purified products were sent to Macrogen Inc. for DNA sequencing. A complete list of all accessions sampled for ITS and the two chloroplast spacers is given in Appendix S1.

### Data Analyses

Five accessions of *Lacomucinaea* and one accession of *Osyridicarpos* were used as outgroups. *Buckleya* is also a member of Thesiaceae (Der & Nickrent, 2008; Nickrent & al., 2019) and has been shown to be sister to the other genera in Thesiaceae.

Although the *Buckleya* sequences were alignable with other taxa, their inclusion added a number of unique gaps to the alignment but did not improve ingroup resolution. For this reason, they were not included in our analyses. To test for species monophyly and examine intraspecific variation across different localities, two or more sequences were obtained from 85 species (Appendix S1).

Sequencher^®^ (Gene Codes Corp. version 4.2) was used to edit the electropherograms and to assemble contiguous sequences. The sequences were then imported into Se-Al v2.0a11 (Rambaut, 2004) and manually aligned. Although numerous gaps were introduced in the alignment, gap coding was not employed owing to their questionable phylogenetic value. For example, many gaps are autapomorphic and others occur at the boundaries of single nucleotide repeats (e.g. CCCCCC vs. CCCCC). Experiments with the automated multiple sequence alignment program MAFFT version 6 (Katoh & al., 2009) were conducted using a variety of PAM / K and gap penalty parameters. Visual inspection and ML tree scores indicated that manual alignment was superior. Alignments are provided in Appendix S2 (ITS) and Appendix S3 (cpDNA).

The data matrices were analyzed on the CIPRES Science Gateway (Miller & al., 2010) using maximum parsimony (MP), Bayesian inference (BI), and maximum likelihood (ML) methods. MP analyses were performed in PAUP* version 4.0a (Swofford, 2002). All characters were treated as unordered with equal weight. Searches for the most parsimonious trees were performed using the two-stage strategy described in Costea & al. (2015). Support for clades was inferred by nonparametric bootstrapping (Felsenstein, 1985), using 1000 heuristic bootstrap replicates, each with 20 random addition cycles with one tree held at each step, TBR branch swapping, and MULTREES option off (DeBry & Olmstead, 2000). Summary descriptions of the analyses, for individual as well as combined datasets, are provided in Table 1.

**Table 1.**
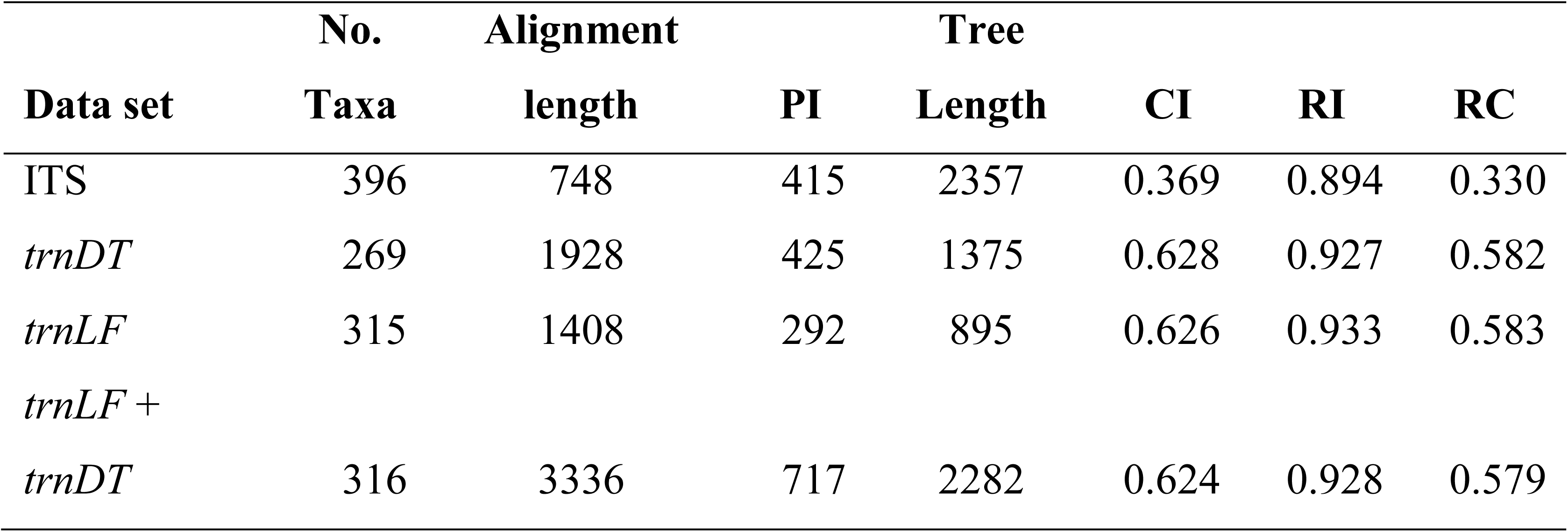
Characteracteristics of different data sets. PI = Number parsimony informative sites; CI = Consistency index minus uniformative sites; RI = Retention index; RC = Rescaled consistency index.

Bayesian phylogenetic inferences were performed with the program MrBayes v.3.2.6 (Ronquist & al., 2012). For all data matrices the symmetrical model with equal base frequencies and a gamma distributed rate variation among sites (SYM + G) was selected with the Akaike Information Criterion (AIC) implemented in MrModeltest v.2.3 (Nylander, 2004). Two independent analyses were executed, each with four chains, for 20 million generations each using default priors. Trees and parameters were saved every 1000 generations, each run producing 20,001 trees. For each of the runs the first 25% of all generations were discarded as burn-in. Both the 50% majority-rule consensus tree and the posterior probability of the nodes were calculated from the 30,002 remaining trees with MrBayes (15,001 from each run). As a measure of convergence, the average standard deviation of split frequencies at the end of the analyses were below 0.01 for all datasets. In the discussion, we have described the range of support of the clades based on the Bayesian Posterior Probabilities (BPP) as very low (0.50--0.70), low (0.71--0.80), moderate (0.81--0.90), good (0.91--0.95), high (0.95--0.99) and the highest (1.0).

Maximum likelihood trees were generated using GARLI version 2.10 (Zwickl, 2006) estimating the best-fit model parameters in the analysis. The starting tree topology was generated by the program using a fast ML stepwise-addition algorithm. The model of nucleotide substitution was General Time Reversible (GTR) with a discrete gamma model with four substitution rate categories with a gamma shape parameter and proportion of invariant sites, both estimated from data. 250 bootstrap repetitions were performed, the best trees from each bootstrap replicate were assembled into a single Nexus file and the majority rule consensus tree obtained in PAUP*. Phylograms (BI) and cladograms (MP and ML) resulting from analysis of the ITS, *trnDT*, *trnLF*, and concatenated chloroplast genes (cpDNA) datasets are provided in Supplmentary Figs. S1--S13.

## RESULTS

DNA was obtained from ca. 590 accessions. Gene sampling was most complete for nuclear ITS with 396 accessions sequenced. Nearly all of the accessions have ITS sequences because PCR amplification and sequencing was successful for over 70% of the samples attempted. Lower sampling was obtained for both chloroplast spacers, with 269 *trnDT* and 315 *trnLF* sequences generated (316 for the concatenated cpDNA partition, Appendix S1). Thirty of the accessions sequenced were not identified to species; however, 12 of these were members of the taxonomically complex *T.* sect. *Frisea* (Clade 39). In total, sequence data were obtained from 196 named species. Among the 396 accessions eight of these likely represent species new to science. In some cases, unidentified accessions represent samples where we disagreed with the original identification on the herbarium label, but could not be placed definitively in an existing species. Approximately 70% of the sequences were derived from herbarium samples, the oldest being over 100 years old. In total, ca. 230 *Thesium* species were sequenced in this study.

The ITS alignment was approximately half as long as either *trnDT* or *trnLF*, however, it yielded more parsimony informative sites proportionally than either of the two chloroplast spacer regions (Table 1), indicating the presence of more phylogenetic signal. The cpDNA dataset had 717 parsimony informative characters but generated over 5800 most parsimonious trees.

Support for various clades was reflected by Bayesian Inference Posterior Probabilities (BIPP), Maximum Parsimony Bootstrap (MPBS), Maximum Likelihood Bootstrap (MLBS) values for the four datasets (ITS, *trnDT*, *trnLF*, and cpDNA). The ITS Bayesian cladogram is shown in Figs. 2A-D; the same results displayed as a phylogram are presented in Supplementary Fig. S1 and the trees resulting from the MP and ML analyses with the corresponding BS values are presented in Figs. S2 and S3, respectively. The cpDNA Bayesian cladogram is shown in Fig. S4 and the phylogram in Fig. S5. The trees resulting from the MP and ML of the cpDNA dataset are available as Figs. S6 and S7, respectively.

**Fig. 2.**
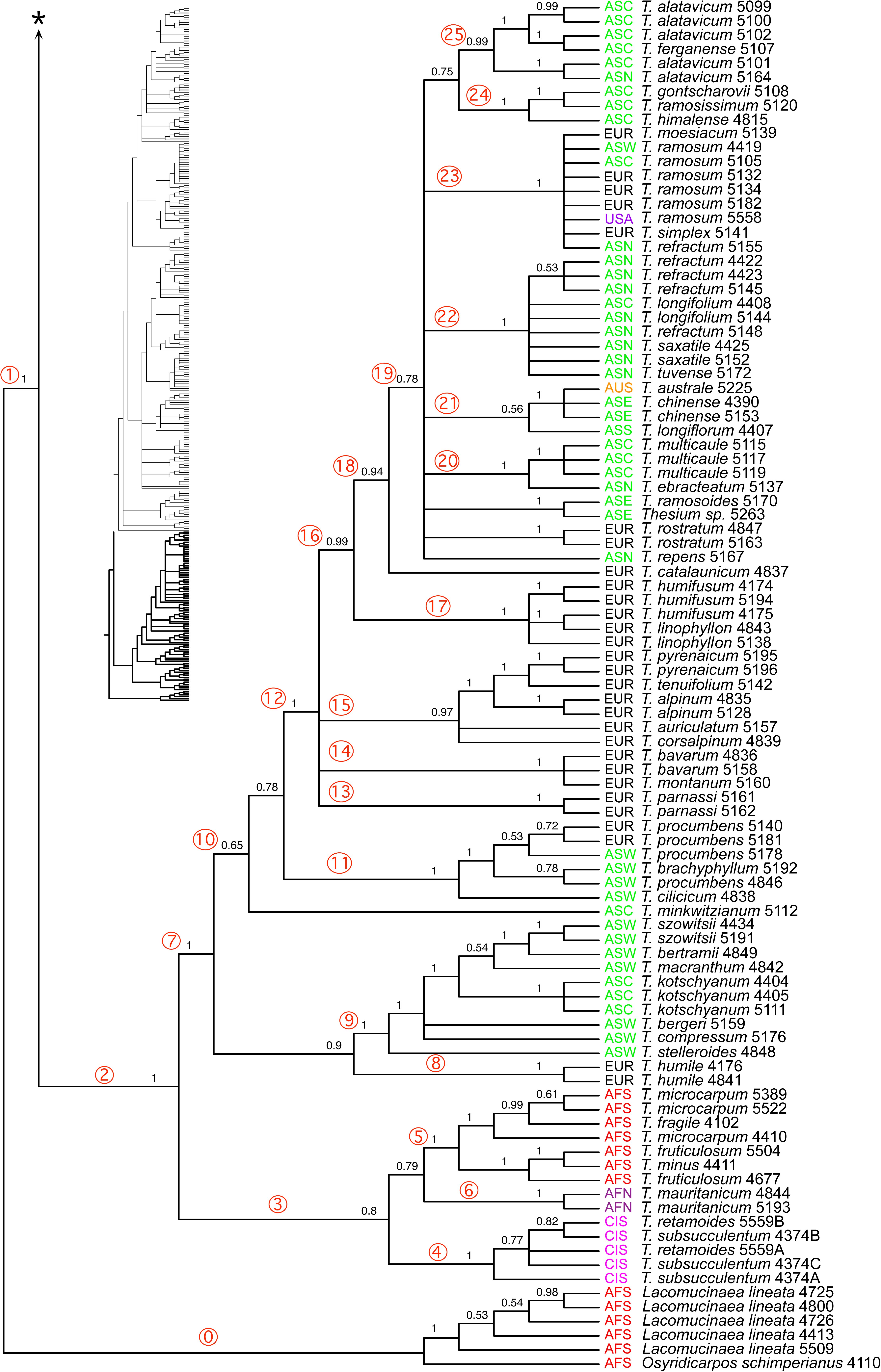
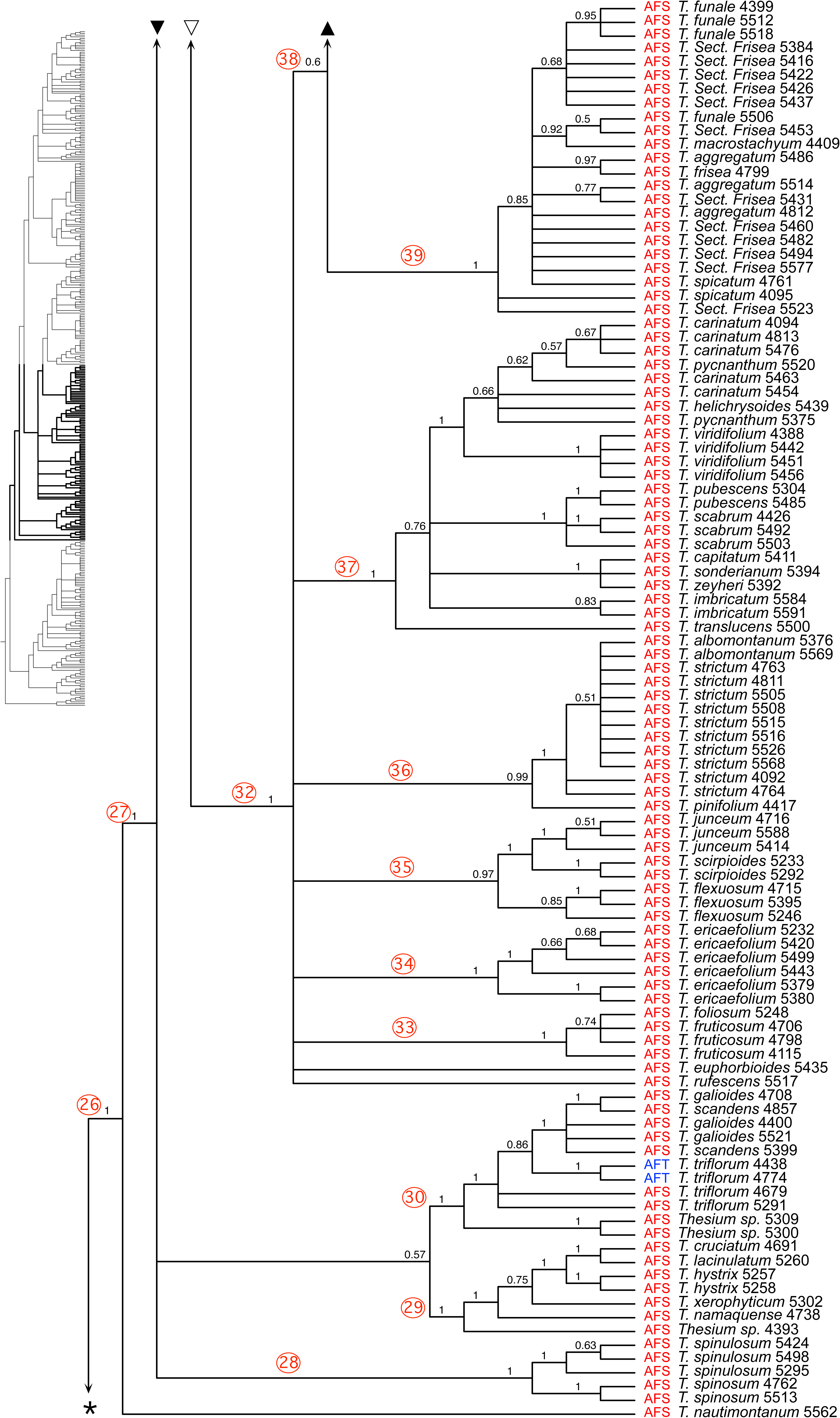
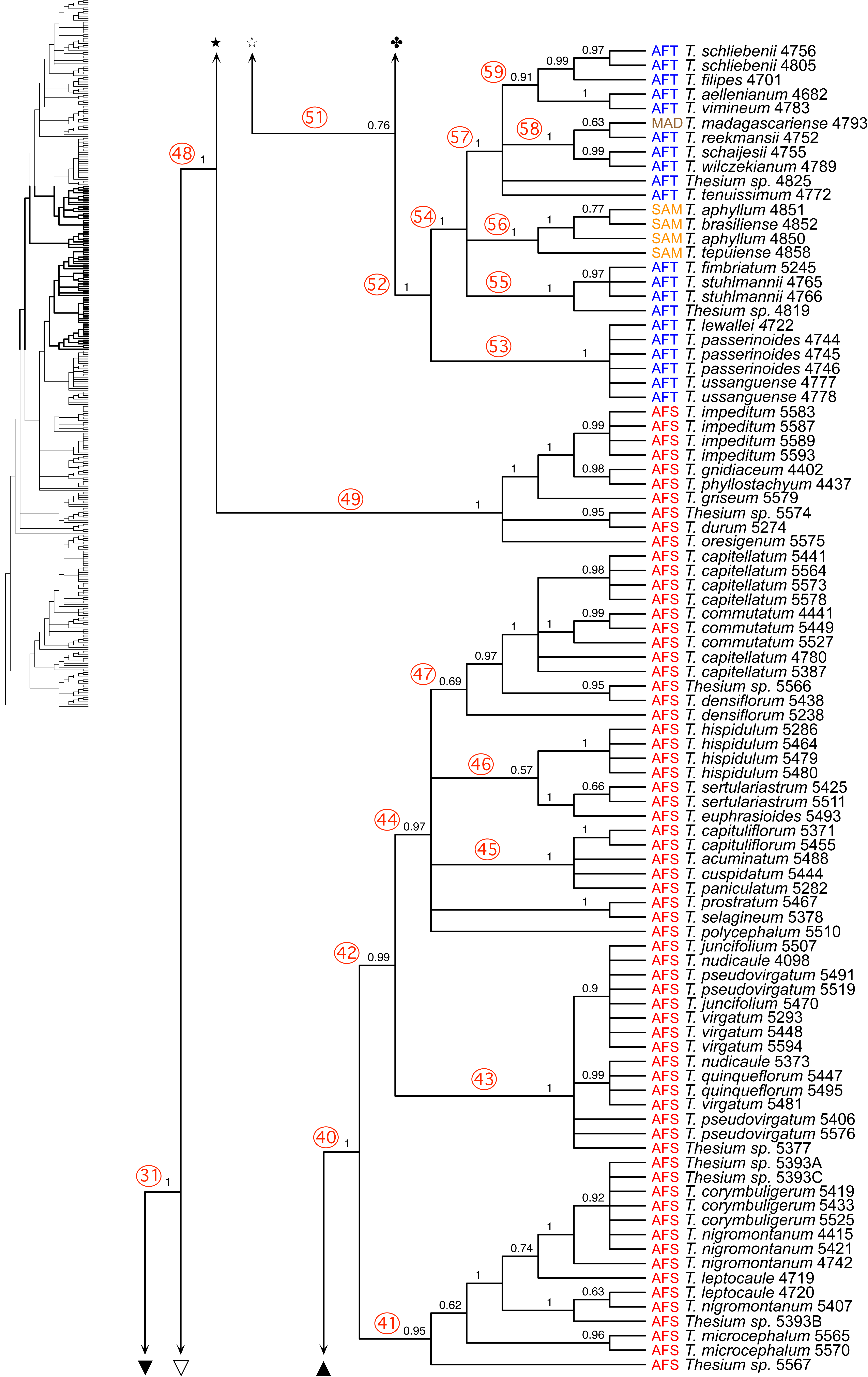
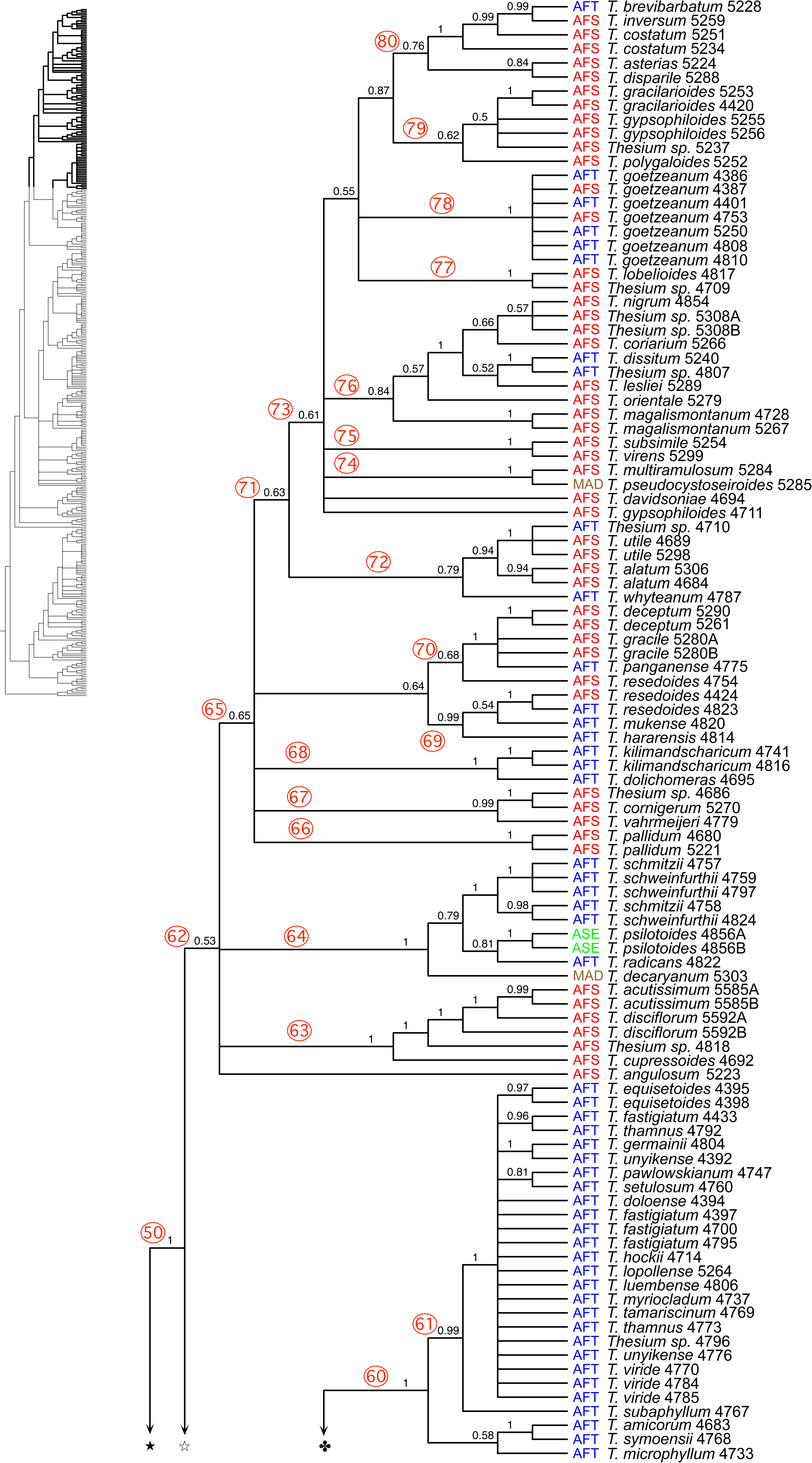
Phylogenetic analysis of *Thesium* using nuclear ribosomal internal transcribed spacer (ITS) sequences. This tree, rooted with *Osyridicarpos* and *Lacomucinea* (outgroups) gives Bayesian posterior probabilities at the nodes. Clades discussed in the text are numbered. Geographical sources for the sampled taxa are abbreviated as follows: AFN = Northern Africa, AFT = Tropical Africa, AFS = Southern Africa, ASC = Central Asia, ASE = Eastern Asia, ASN = Northern Asia, ASS = Southern Asia, ASW = Western Asia, AUS = Australia, CIS= Canary Islands, EUR = Europe, MAD = Madagascar, SAM = South America, and USA = United States of America.

The trees obtained with Bayesian, MP and ML with their PP and BS values from the individual *trnDT* and *trnLF* datasets are available as Figs. S8--S10 for *trnDT* and S11--S13 for *trnLF*. The ITS and chloroplast spacer datasets were not concatenated because significant numbers of conflicting relationships between them were detected, especially in the Core Cape Clade. This topic is further explored in the Discussion. Many of the clades present on the ITS Bayesian cladogram (Fig. 2) have been numbered, particularly those with high posterior probabilities (0.90 and greater). These numbers were included in all the chloroplast trees before the accession names to facilitate their comparison to the topologies present on the nuclear ITS tree.

The backbone relationships obtained with chloroplast and ITS trees are congruent with respect to the following. After divergence from the clade composed of the two outgroup genera (*Osyridicarpos* and *Lacomucinaea*), the ancestral *Thesium* taxon (Clade 1) underwent a major split that generated two clades (2 and 26), both of which contain present day species that occur in South Africa. Clade 2 contains a diverse assemblage of taxa that includes many members that have previously been considered sufficiently distinct to warrant separate generic status (i.e. *Chyrysothesium*, *Kunkeliella*, and *Thesidium*) and included in subgenera *Thesium* and *Hagnothesium*. In addition, almost all of the species currently present in Europe, N Africa and Asia were resolved in Clade 2. Clade 26 contains all remaining species of *Thesium* that are primarily from continental Africa. At the base of Clade 26 on the ITS trees is a recently named enigmatic species, *T. nautimontanum* M.A.García, Nickrent & Mucina endemic to the Matroosberg Mt. of South Africa.

Clade 27 is not fully resolved as shown by the presence of three polytomies (Clades 28, 29/30, and 31). Clades 28--30 contain species that often show thyrsoid inflorescences (racemes with partial inflorescences as dichasia or reduced dichasia) or single flowers, including *T. galioides* A.DC.*, T. scandens* E.Mey.*, T. spinosum* L.f.*, T. spinulosum* A.DC., and *T. triflorum.* These taxa are noted here because in trees from the chloroplast spacers they emerge as successive sister clades to the remaining *Thesium* species, not *T. nautimontanum* (Figs. S4--S13). Clade 31 contains all the Cape and tropical African species of *Thesium* (c. 170 species sampled here). Within it, Clade 32 contains 50 species found in the CFR (others present within Clade 48) and represents the monophyletic subgenus *Frisea*. Exclusively tropical African species only begin appearing in Clade 48, with the exception of *T. triflorum* that has a wide distribution spanning both regions. Clade 49 is significant as it is sister to the remaining *Thesium* clades on the ITS trees and is composed entirely of South African species of biogeographical significance (see Discussion). Clade 48, resolved with the highest support, is subgenus *Psilothesium* and within it, Clade 51 is composed mainly of tropical African species, with some interesting exceptions. Three species from South America emerge as Clade 56 and in Clade 58 one species is endemic to Madagascar. Clade 61 contains at least 16 tropical African species that all emerge from a polytomy. The last major clade (62) includes Clades 63--80 and can be informally referred to as the “Grassland Clade”. These species occur in southern as well as tropical Africa and in one clade (64), the geographic distributions implicate long-distance dispersal. One species (*T. decaryanum* Cavaco & Keraudren) is another tropical African taxon that dispersed to Madagascar (the others being *T. madagascariense* A.DC. of Clade 58 and *T. pseudocystoseiroides* Cavaco & Keraudren of Clade 74). Sister to the tropical African taxon *T. radicans* is *T. psilotoides* Hance from eastern Asia. The remaining 41 species that occur in Clades 66--80 are species of grassy biomes that occur in both southern and tropical Africa. These species present taxonomic difficulties and neither ITS nor the chloroplast genes provide sufficient signal to resolve all relationships.

## DISCUSSION

### Generic Concepts

The concatenated 3-gene analysis of Santalaceae *s. lat.* in Der & Nickrent (2008) included all five genera recognized at that time. The “*Thesium* Clade”, now recognized as Thesiaceae (Nickrent & al., 2010) showed *Buckleya* sister to the remaining species followed by *Osyridicarpos* that was sister to two clades, one containing two South African species of *Thesium* and one containing *Kunkeliella* and *Thesidium*. Later work using nuclear rDNA ITS and chloroplast spacers (Moore & al., 2010) included *Buckleya*, *Thesium*, and *Thesidium* but did not sample *Osyridicarpos* or the Canary Island endemic *Kunkeliella*. Their tree topology showed *Thesidium* sister to a clade of Eurasian taxa and that clade sister to *Thesium*. This topology presented a problem if one wished to use monophyly as primary criterion for applying the rank of genus (Backlund & Bremer, 1998). Several solutions were possible: 1) use a name other than *Thesium* for the Eurasian Clade and include the dioecious South African genus *Thesidium*, 2) use a name other than *Thesium* for the African Clade, or 3) combine *Thesidium* with *Thesium*. The first choice was undesirable for nomenclatural reasons because the type species of the genus is *T. alpinum* L. (Hitchcock & Green, 1929) which is Eurasian. The second choice was also undesirable because the vast majority of *Thesium* species occur in Africa. The third choice, which was actually followed by Forest & Manning (2013), combined *Kunkeliella* into *Thesium* based upon the topology of the molecular tree in Der & Nickrent (2008). The first molecular phylogenetic study that analyzed all recognized genera of Thesiaceae was by Nickrent & García (2015). The genus *Kunkeliella*, *T. mauritanicum* from North Africa, and three species that were considered by Hendrych (1994) to be in the genus *Chrysothesium* (*T. stelleroides* Jaub. & Spach*, T. minkwitzianum* B.Fedtsch., and *T. cilicicum* Hausskn. ex Bornm.) were all included. All of these taxa were resolved as part of the Eurasian Clade of *Thesium*. In addition, one South African taxon (*Thesium lineatum*) was found to be sister to *Osyridicarpos*. The option to include that species in *Osyridicarpos* was considered but given the two taxa exhibited many morphological differences, the authors named it as a distinct genus *Lacomucinaea*.

### Phylogenetic relationships, taxonomy and morphological characters

The discussion that follows is based mostly on the ITS phylogeny for several reasons: 1) better taxon sampling than for the two chloroplast markers, 2) better tree resolution and generally higher clade support, 3) incongruences between the nuclear and chloroplast phylogenies that apparently did not originate by reticulation events (see Discussion), 4) species relationships obtained from *trnDT* and *trnLF* analyzed alone and combined are in many cases incongruent with morphology, and 5) many species of *T.* subgen. *Frisea* that are morphologically well defined and strongly supported as monophyletic with ITS, are resolved as polyphyletic on the cpDNA trees (i.e. *T. strictum* P.J.Bergius). The clades and their associated Bayesian Posterior Probability (BPP) values are listed below with discussion of their composition. A list of what we consider to be valid *Thesium* species and their synonyms (360 species in total), taking into account the results of this study, is given in Appendix S4.

**0. *Lacomucinaea* Clade (BIPP 1). –** *Lacomucinaea lineata* (L.f.) Nickrent & M.A.García (Fig. 3A), previously known as *Thesium lineatum*, was classified as a distinct genus by Nickrent & García (2015) based on molecular and morphological evidence. This genus is sister to *Osyridicarpos* and that clade is sister *Thesium*, relationships with the highest support from both ITS and the chloroplast genes (Fig. 2A). One may ask why *Lacomucinaea* was not included within *Osyridicarpos*. These two genera share several morphological features, e.g. primary phloem fiber bundles near the stem surface, bisexual flowers, lack of a swollen pedicel, and fleshy fruits with smooth exocarps. They differ in other characters such as habit (shrub vs. scrambling subshrub), leaf morphology, leaf persistence, inflorescence structure, presence of external floral gland, style length and ribbing on ovary. Four species have been described in *Osyridicarpos* although it is frequently treated as monotypic (*O. schimperianus* A.DC.). Further studies might prove that several species should be recognized in the genus, all of them very different from *Lacomucinaea*. Thus, the preponderance of molecular and morphological evidence justifies recognition of two genera.

**Fig. 3.**
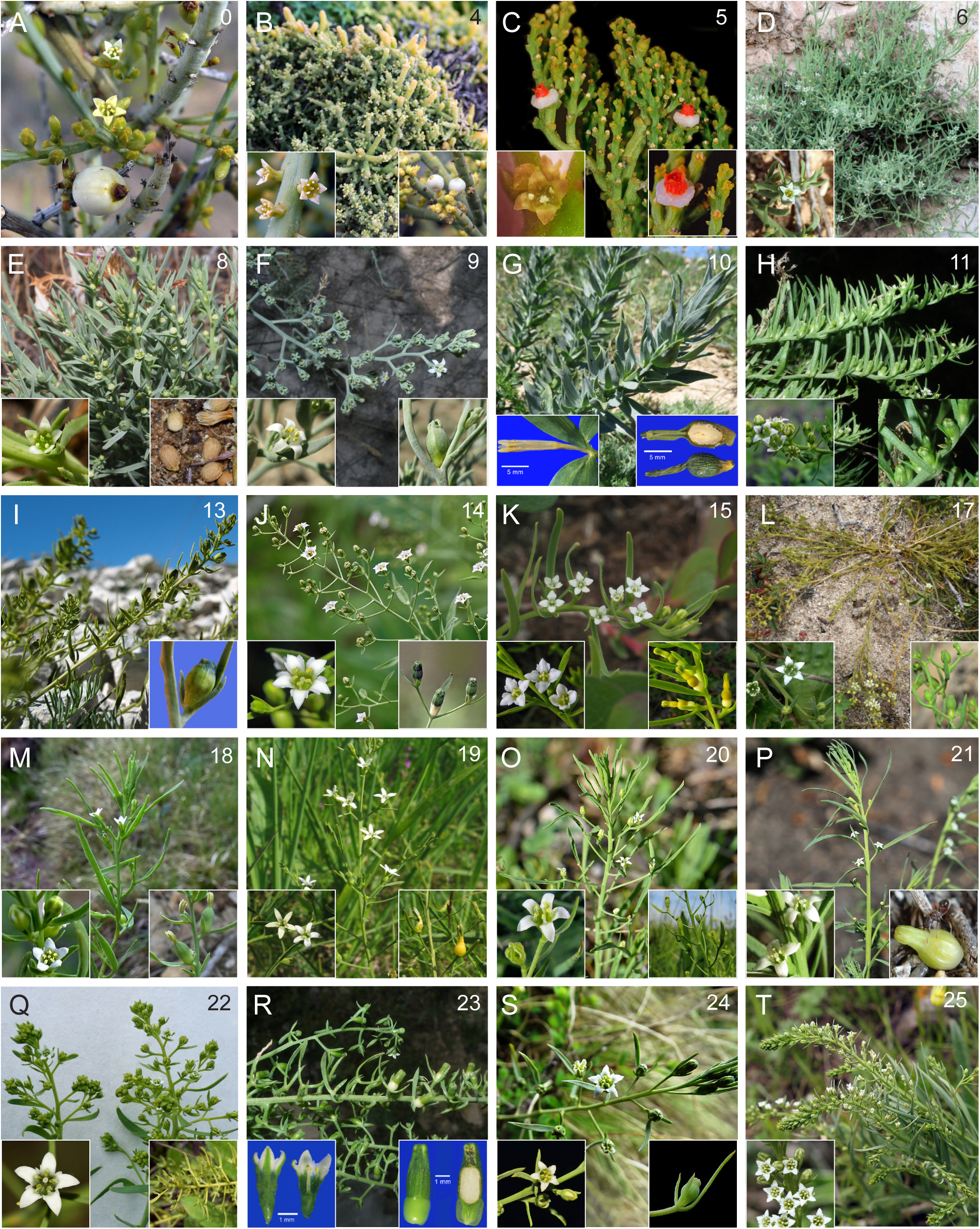
Photographs of the outgroup (*Lacomucinaea*) and representatives of the major clades of *Thesium*. The numbers in the upper right corners correspond to clade numbers on the Bayesian trees (Fig. 2A). Details of flower and/or fruits are shown in the insets. **A,** *Lacomucinaea* clade, *L. lineata*; **B,** *Kunkeliella* clade, *T. subsucculentum*; **C,** *Hagnothesium* clade, *T. fragile*; **D,** *Mauritanica* clade, *T. mauritanicum*; **E,** *Humilia* clade, *T. humile*; **F,** *Macranthia* clade, *T. szowitsii*; **G,** Eurasian clade, *T. minkwitzianum*; **H,** *Procumbens* clade, *T. procumbens*; **I,** *Parnassi* clade, *T. parnassi*; **J,** *Bavarum* clade, *T. bavarum*; **K,** *Alpina* clade, *T. alpinum*; **L,** *Linophylla* clade, *T. humifusum*; **M,** Second Asian Radiation, *T. catalaunicum*; **N,** *Rostratum* clade, *T. rostratum*; **O,** *Multicaule* clade, *T. ebracteatum*; **P,** *Australe* clade, *T. chinense*; **Q,** *Longifolia* clade, *T. refractum*; **R,** *Ramosum* clade, *T. ramosum*; **S,** *Himalensia* clade, *T. himalense*; **T,** *Alatavicum* clade, *T. alatavicum*.

**1. *Thesium* Clade (BIPP 1). –** This clade (Fig. 2A) contains all species now classified as *Thesium* and includes taxa formerly placed in *Austroamericium, Chrysothesium, Kunkeliella,* and *Thesidium.* The monophyly of *Thesium* including these genera and excluding *Lacomucinaea* is also resolved with chloroplast markers with the highest support.

**2. CMPB Clade (BIPP 1). –** This strongly supported clade (Fig. 2A) is named based on the distribution of species that occur in phytochoria that are peripheral to much of the African continent: Capensis (CFR of South Africa), Macaronesia (Canary Islands and Madeira), and Paleoboreal (North Africa, Europe, Asia).

There is no apparent morphological synapomorphy for this clade but most of the Eurasian and North African species of *Thesium* show corolla lobes with white bands along the edges of the abaxial surface and lateral auricles at the base that are variable in shape and size. In the cases in which they are not clearly developed, the margins of the lobes are swollen and more or less erose. These structures are also present in *T. mauritanicum* (Clade 6) and *Thesium* sect. *Kunkeliella* J.C.Manning & F.Forest (Clade 4) and could be homologous to the hairs on the margins of the petals of the African species in *Thesium* sect. *Barbata* and subsect.

*Fimbriata* A.W.Hill. This morphology is not seen in members of *Hagnothesium* (Clade 5) likely owing to the evolution of extremely reduced flowers in this clade. When the fruit is developing, the corolla lobes become incurved, and in those species with longer lobes a characteristic “window” appears at the base (see e.g. inset of Fig. 3F).

Zhigila & al. (2020) considered that long styles with anthers below the stigma is a character exclusive of *Thesium* subg. *Thesium*, but this character is also present in many tropical African species in Clade 51 of subgenus *Psilothesium*.

**3. CMNA Clade (BIPP 1). –** This clade (Fig. 2A) is a subset of the CMPB Clade that includes taxa present in the Cape, Canary Islands, and just the North African part of the Paleoboreal region. The clade contains all species formerly placed in *Kunkeliella*, *Thesidium* as well as some *Thesium*.

**4. *Kunkeliella* Clade (BIPP 1). –** This clade (Fig. 2A), composed of endemics to the Canary Islands archipelago, was formerly placed in the genus *Kunkeliella* (Stearn, 1972) and members were reclassified as *Thesium* sect. *Kunkeliella* by Forest & Manning (2013). Two species were sampled here: *T. subsucculentum* (Kämmer) J.C.Manning & F.Forest (Fig. 3B) from the north coast of Tenerife (Canary Islands, Spain) and *T. retamoides* (A.Santos) J.C.Manning & F.Forest from the southern part of the same island. The three ITS clones of *T. subsucculentum* (A, B, C) and two clones of *T. retamoides* (A, B) were not monophyletic based on species. However, Rodríguez-Rodrígez & al. (2022) found enough genetic variation based on microsatellite markers to consider them different species, a conclusion also supported by morphology. On Tenerife, all the species have stems covered with an indument of tiny hairs, whereas *T. canariense* (Stearn) J.C.Manning & F.Forest, endemic to Gran Canaria Island, is glabrous. Otherwise, the flowers, fruits and general habit of these plants are similar except for the smaller flowers of *T. canariense*. That species is the only one in the genus known to be gynodioecious, i.e. with bisexual and female flowers on separate plants (Pérez de Paz & al., 2015). Stearn (1972) considered this a genus separate from *Thesium* based mainly upon its possession of isopolar pollen and drupaceous fruits. Isopolar pollen is of questionably diagnostic value as it has been found in several South African species of *Thesium* plus intermediates between heteropolar and isopolar pollen types are known (Lobreau-Callen, 1980). Fleshy fruits are also present in *T. fragile* L.f., *T. triflorum* and *T. radicans*.

Unsampled species include *T. canariense* from Gran Canaria and *T. psilotocladum* Svent. from Tenerife (extinct). Recently, a new species, *T. palmense* P.Pérez & P.Sosa, was described from La Palma, closely related to *T. retamoides* from Tenerife.

**5. *Hagnothesium* Clade (BIPP 1). –** To maintain a monophyletic *Thesium* (Forest & Manning, 2013), species previously classified in the genus *Thesidium* (Sonder, 1857) were reclassified as *Thesium* sect. *Hagnothesium* A.DC. Zhigila & al. (2020) later raised this section to the rank of subgenus (*Thesium* subgen. *Hagnothesium* (A.DC.) Zhigila & al.). These species are CFR endemics and include annual to shrubby plants with mostly tetramerous flowers and a dioecious sexual condition. Some species, including *T. microcarpum* A.DC, *T. fragile* (Fig. 3C) and *T. quartzicolum* Zhigila & al. have morphologically similar male and female plants (monomorphic) whereas *T. fruticulosum* (A.W.Hill) J.C.Manning & F.Forest, *T. minus* (A.W.Hill) J.C.Manning & F.Forest, *T. strigulosum* A.DC. (=*T. hirtum* (Sond.) Zhigila & al., nom. illeg.), *T. longicaule* Zhigila & al. and *T. leptostachyum* A.DC. are dimorphic. In the latter case, male plants usually have small herbaceous bracts and female plants have conspicuous leafy bracts. Although we only sampled five species, the ITS phylogeny (Fig. 2A) resolves two highly-supported clades separating sexually dimorphic and monomorphic species.

The nuclear and chloroplast phylogenies suggest that *T.* subgen. *Hagnothesium* is an early divergent lineage that, in spite of being endemic to the CFR, is more closely related to Eurasian, North African, and Canary Islands lineages (*T.* subgen. *Thesium*). According to our chloroplast phylogeny it is sister to *T.* subgen. *Thesium*, however this relationship is not resolved by ITS. Instead, *T.* subgen. *Hagnothesium* is nested within *T.* subgen. *Thesium* (Clade 2), making the latter paraphyletic. Unsampled species include *T. leptostachyum*, *T. longicaule*, *T. quartzicolum*, and *T. strigulosum*.

**6. *Mauritanica* Clade (BIPP 1). –** In our sampling this clade contains a single species, *T. mauritanicum* (Fig. 3D), from North Africa (Morocco and Algeria). Its position on the phylogenetic tree (Fig. 2A) indicates that it is phylogenetically distinct from the African clade (26). When Battandier (1889) described *T. mauritanicum* he suggested a relationship with *T. bergeri* Zucc. (and its synonym *T. graecum* Boiss. & Spruner) from western Asia and Greece. However, ITS and chloroplast sequences show that species as part of Clade 9. The flowers of *T. mauritanicum* differ little from Eurasian *Thesium* species but dense pubescence on the stems, leaves, bracts, abaxial surface of the petals and fruits are uncommon characters in the genus. This clade corresponds to *Thesium* ser. *Mauritanica* Hendrych that also included the Libyan endemic *T. erythronicum* Pamp. It shares with *T. mauritanicum* a subshrub habit and reticulate fruits; however, the two species differ in several vegetative and inflorescence features suggesting that it might be related to other Paleoboreal species, possibly in Clades 8 or 9.

**7. Paleoboreal Clade (BIPP 1). –** This clade (Fig. 2A) was also recovered with the chloroplast dataset (BIPP 1) and contains most of the European and Asian species included by Hendrych (1972) in subgenera *Thesium* and *Chrysothesium*. The three species sampled that were classified by Hendrych (1972) in *Chrysothesium* are not monophyletic (Nickrent & García, 2015). Our sampling includes all the series in *T.* sect. *Thesium* except for *T.* ser. *Indica* Hendrych (one species, *T. indica* Hendrych) and *T.* ser. *Macrocarpa* Hendrych (one species, *T. jarmilae* Hendrych, from the Himalayas) and all the sections except for *T.* sect. *Wightiana* Hendrych (one species, *T. wightianum* Wall. ex Wight, from southern India) and *T.* sect. *Coarctiflora* Hendrych with one species, *T. coarctiflorum* Hendrych, considered by Miller (1982) as a synonym of *T. bergeri* (Clade 9). *Thesium wightianum*, however, might be related to tropical East African species (see Discussion on Clade 64).

**8. *Humilia* Clade (BIPP 1). –**This clade (Fig. 2A) contains *Thesium humile* Vahl (Fig. 3E), one of the four species of *T.* ser. *Humilia* Hendrych. The phylogenetic position is resolved as sister to the *Macranthia* clade (9) with moderate support (BIPP 0.90) whereas it is resolved as sister to the core Eurasian clade (12) in the chloroplast phylogeny, although with low support (BIPP 0.77). *Thesium humile* is one of the few annual species of the genus. Hendrych (1972) also included in the series *T. ifrikianum* Hendrych, *T. tuneticum* Hendrych, and *T. macedonicum* Hendrych, none of which were sampled here. The first two species are restricted to small areas in North Africa and might be synonyms of *T. humile*, whereas *T. macedonicum* is characteristically a puberulent-hispid perennial species known from the mountains bordering Greece and North Macedonia (Hendrych, 1964). This species, however, might be related to species in Clade 11. In contrast, *T. humile* has a wide distribution around the Mediterranean and to the Canary Islands. All of these species have spike-like inflorescences and fruits bearing reticulate nerves, features interpreted by Hendrych (l.c.) as primitive. However, these characters are not exclusive to this series and are present in other Eurasian species.

**9. *Macranthia* Clade (BIPP 1). –** This clade (Fig. 2A), which also received the highest support with the chloroplast datasets, includes mostly SW Asian species with varied morphological features but all appear to have fruits with reticulate surface venation. The clade name derives from *Thesium* sect. *Macranthia* (Bobrov) Hendrych that in our sampling includes *T. bertramii* Azn., *T. kotschyanum* Boiss.*, T. macranthum* Fenzl, and *T. szowitsii* A.DC. (Fig. 3F). However, components of other sections and series are also resolved in this clade such as *T.* ser. *Procumbentia* Bobrov (*T. bergeri*), *T.* sect. *Compressia* Hendrych (*T. compressum* Boiss.) and *Chrysothesium* (*T. stelleroides* Jaub. & Spach). *Thesium stelleroides*, an early-branching member of this clade, is endemic to Turkey and it has a number of distinctive morphological features such as the yellow, tubular flowers that are arranged in spikes and pubescent ovaries and fruits. *Thesium bergeri* is morphologically quite different from *T. stelleroides*. Its flowers are in racemes or in cymules with the bract much longer than the flowers. The perianth is much shorter than the fruit and the corolla lobes are triangular and ca. 0.5 mm long. It is the only species in this clade distributed mostly in SE Europe (Balkan Peninsula and the Aegean). *Thesium compressum* (=*T. lycaonicum* Bornm.) differs from other members of *T.* sect. *Macranthia* by its elongated spike-like inflorescences (with axillary and sessile flowers) and bracts and bracteoles that are shorter than the fruits. The corolla lobes in this species are very short (up to 0.75 mm), triangular in shape, and shorter than the reticulate fruit.

With the addition of *Thesium stelleroides*, *T.* sect. *Macranthia* is resolved as monophyletic in our ITS phylogeny. The species are characterized by large infundibuliform flowers with a variable tube length, for example longer than the lobes in *T. kotschyanum* and *T. bertramii* or shorter in *T. szowitsii*. The inflorescences are composed of lateral cymules of 1--5 flowers with totally adnate bracts (or partially adnate in basal cymules). The corolla remnants are about as long as the reticulate fruits (Hendrych, 1972). *Thesium kotschyanum*, present in Iran, Iraq and Turkmenistan, has prostrate to ascending stems with most flowers in lateral cymules and floral tubes about as long as the lobes. Townsend (1980) was unable to differentiate collections of this species from Iraq from *T. impressum* Steud. ex A.DC. The exact identity of the specimen from Turkey, here identified as *T. bertramii*, is not clear. Miller (1982) indicates that this species has corolla lobes as long as or slightly shorter than the tube, but in our specimen the lobes are slightly longer than the tube. *Thesium bertramii* has lateral cymules (dichasia) with little branching and frequently the flowers are solitary. In our specimen (4849) the lateral cymules have 1--3 flowers.

Voucher 4842 identifed here as *T. macranthum* might also be *T*. *impressum*; fruit characters that are used to separate these species were lacking in this specimen. According to Miller (1982), *T. macranthum* has reticulate and strongly 10-ribbed fruits with convoluted venation whereas in *T. impressum* the venation is not convoluted. *Thesium macranthum* has inflorescences “widely” branched, as in this specimen, with lateral cymules (dichasia) with 3--5 flowers, bracts with denticulate margins, yellowish corolla lobes much longer than the tube, and a perianth longer than the fruit.

The taxonomy of this group is difficult because the characters used to separate species are variable and a study covering their whole geographical area does not exist. Unsampled species that might be synonyms of those sampled include *T. billardieri* Boiss., *T. impressum*, *T. maritimum* C.A.Mey. and *T. scabriflorum* P.H.Davis. *Thesium coarctiflorum*, only known from the type collection from the island of Samos (Greece), was included in the monotypic *T.* sect. *Coarctiflora* by Hendrych (1972) but considered a synonym of *T. bergeri* by Miller (1982). *Thesium tauricolum* Boiss. & Haussk. has inflorescences in spikes similar to *T. compressum* and might be related to this species.

**10. Eurasian Clade (BIPP 0.65). –** This clade, containing many species from Europe to East Asia, received very low support with ITS and was not resolved with chloroplast sequences. *Thesium minkwitzianum* (B.Fedtsch.) Hendrych is sister to the rest of the species in this clade. It is a very distinctive species from central Asia owing to its thick, coriaceous leaves, broadly lanceolate bracts, lanceolate bracteoles, and yellowish, up to ca. 2 cm long tubular flowers arranged in racemes (Fig. 3G). This species was included by Hendrych (1994) in the genus *Chrysothesium* together with *T. stelleroides* (Clade 8), *T. aureum* Jaub. & Spach, and *T. cilicicum* (the latter two in Clade 11). Except for the long tubular flowers, these two species are morphologically very different from *T. stelleroides* and all of them different from *T. minkwitzianum.* The molecular data presented here clearly indicate that *Chrysothesium* is polyphyletic.

Unsampled species that likely reside in the Eurasian clade include the following: *Thesium emodi* Hendrych from the Himalayas is a rhizomatous plant characterized by the bracts free (or in the upper flowers shortly adnate to the peduncle), funnelform to tubular flowers, and fruits with longitudinal nerves. *Thesium tongolicum* Hendrych and *T. remotebracteatum* C.Y.Wu & D.D.Tao have similar distributional areas and morphology as *T. emodi*. Their closest relative might be *T. himalense* Royle ex Edgew. (Clade 24) but in this species the bracts are completely adnate. Most Eurasian species have adnate bracts and Hendrych (1965) could not place *T. emodi* in any series based solely on morphology. As in *T. emodi, T. jarmilae* is the only species of *T.* ser. *Macrocarpa* that has free (or partly adnate at the base) bracts and fruits with conspicuous longitudinal veins (Nianhe & Gilbert, 2003; Ghosh & al., 2021). *Thesium orgadophilum* P.C.Tam is only known from immature individuals collected at the type locality of Xizang Province in China. Another endemic to this province is *T. bomiense* C.Y.Wu & D.D.Tao, a species with the bracts adnate to the peduncle.

**11. *Procumbens* Clade (BIBP 1). –** This clade (Fig. 2A) contains species formerly placed by Hendrych (1972) in *Chrysothesium* (*Thesium cilicicum*) and two series, *T. brachyphyllum* Boiss. (*T.* ser. *Micrantha*) and *T. procumbens* C.A.Mey. (*T.* ser. *Procumbentia*). In the early-branching *T. cilicicum*, the yellow flowers are almost sessile in dense terminal spikes or racemes with long (nearly 8 mm) tubular corollas. *Thesium aureum* (not sampled but possibly a synonym of *T. cilicicum*) is a similar species with a longer corolla tube and a larger fruit with reticulate surface nerves. The species in this clade are distributed mostly in montane Anatolia, the Middle East, western Iran and the Caucasus, with populations of *T. procumbens* or *T. brachyphyllum* in SE Europe. This latter species was considered as synonym of *T. procumbens* by Miller (1982). Conversely, Bobrov (1936) considered them different species based on differences in fruit reticulation (see review by Romo & al., 2004). *Thesium procumbens* has fruits longitudinally nerved, with lateral nerves absent or present and ascending, whereas *T. brachyphyllum* (Fig. 3H) has transverse nerves connecting the longitudinal ones, but these differences are frequently very subtle. The ITS phylogeny shows *T. brachyphyllum* nested within accessions of *T. procumbens*, thus suggesting conspecificity. The inclusion *T. cilicicum* within the *T. procumbens* clade is well-supported in the ITS phylogeny but not in the chloroplast phylogeny where it is resolved as sister to Clade 9, albeit with very low support. Morphologically, the clearest differences are in the flowers, tubular in *T. cilicicum* and broadly campanulate in *T. procumbens*. These are caespitose plants, sharing characters such as the short decumbent to ascending stems, recaulescent floral bracts, and flowers in spike-like or racemose inflorescences.

Unsampled species include *T. aureum*, *T. libanoticum* Ehrenb. ex A.DC (one of the few species of *Thesium* with yellow flowers), *T. krymense* Romo & al., and *T. vlachorum* Aldén. The latter two species were described from Crimea and Greece, respectively, and they share many morphological features with *T. procumbens* (Aldén, 1981; Romo & al., 2004). *Thesium oreogetum* Hendrych, a cespitose species from Turkey known only from type specimens, might be resolved here but fruits are unknown. It was included by Hendrych (1972) in *T.* ser. *Repentia* Bobrov (see Clade 19). *Thesium brachystegium* Post is an enigmatic species described from Syria from which we could not study the type material, but according to the description it might belong in this clade. See Clade 13 for more discussion on other species in *T.* ser. *Micrantha*.

**12. Core Eurasian Clade (BIPP 1). –** This clade (Fig. 2A) contains representatives of 13 of the 19 series proposed for *Thesium* sect. *Thesium* by Hendrych (1972). Over 30 of the 47 species present in these series were sampled, representing collections from Europe and Asia including China. Many of the early-diverging clades contain European species whereas many of the later-diverging clades have Asian species. Clades 13--16 (below) all arise from a polytomy. Backbone relationships within this clade are resolved with higher support using the chloroplast dataset than with ITS. The topology of the chloroplast trees shows ITS Clade 15 as a sister lineage to the rest of the species and, as suggested by ITS, the species with exclusive Asian distribution are resolved in later diverging lineages. Both nuclear and chloroplast topologies are largely congruent barring a few exceptions that are commented upon in the discussion of individual ITS clades.

**13. *Parnassi* Clade (BIPP 1). –** The two accessions of *Thesium parnassi* A.DC. (Fig. 3I), a species from southern and eastern Europe, emerge as part of the polytomy for the Core Eurasian clade (Fig. 2A) without clear phylogenetic relationships. However, the chloroplast trees resolved it in a strongly supported clade together with other European species resolved in ITS Clades 14--20 and 23. This is one of the seven species included by Hendrych (1972) in

3. *T.* ser. *Micrantha* of which only two were sampled here, *T. brachyphyllum* (Clade 11) and *T. parnasii*. Species in this series are alpine, generally prostrate, perennials (oreophytes) with short branches and short-pedunculate racemose inflorescences. The morphological features used to define the series seem to be related to their common habitat rather than their phylogenetic relationship and the series is polyphyletic. The other unsampled species in *T.* ser. *Micrantha* that might be related to *T. parnassi*, or resolved in other clades of the core Eurasian lineage, are *T. sommieri* Hendrych (Alpi Apuane and Apennines, Italy) and *T. kyrnosum* Hendrych (Corsica). *Thesium hispanicum* Hendrych, from N Spain, was treated as a synonym of *T. pyrenaicum* (Clade 15) by Pedrol & Laínz (1997).

**14. *Bavarum* Clade (BIPP 1). –** This clade (Fig. 2A) includes *Thesium bavarum* Schrank (Fig. 3J) and *T. montanum* Ehrh. ex Hoffm., two species considered as synonyms by Hendrych (1993) in Flora Europaea (the valid name at the rank of species would be *T. bavarum*). This species is a perennial non-stoloniferous plant with erect stems and broad, lanceolate leaves with 3--5 veins, branched paniculate inflorescences and flowers typically with developed auricles on the margins of the corolla lobes. Although frequently considered a subspecies of *T. linophyllon* L. (Clade 17) and included together with it in *T.* ser. *Linophylla* Bobrov by Hendrych (1972), this latter species is stoloniferous, has narrower, rigid leaves and our phylogenies suggest that they are distinct (see *Humifusum* Clade below). Although ITS resolves *T. bavarum* as part of the core Eurasian Clade polytomy, the chloroplast phylogeny places the species in a strongly supported clade together with *T. parnassi* and other species in ITS Clades 17, 19, 20, and 23.

**15. *Alpina* Clade (BIPP 0.97). –** This highly-supported clade (Fig. 2A), also resolved with chloroplast sequences, contains European and northern Asian species classified by Hendrych (1972) in three different series: *Thesium* ser. *Alpina* Hendrych (*T. alpinum* (Fig. 3K), *T. corsalpinum* Hendrych), *T.* ser. *Auriculata* Hendrych (*T. auriculatum* Vandas), and *T.* ser. *Saxatilia* Hendrych (*T. pyrenaicum* Pourr.). *Thesium pyrenaicum* and the two sampled species of *T.* ser. *Alpina* all have fruits bearing at their apex remnants of the hypanthium and corolla lobe that together are at least half the length of the nutlet. Our voucher 5142 was tentatively identified as *T. tenuifolium* Saut. ex W.D.J.Koch, which is generally treated as a subspecies or a synonym of *T. alpinum*. This specimen has many morphological similarities with *T. pyrenaicum* but it was collected in NW Russia, out of the known geographical range of that species. Morphologically it is what Tzvelev (1996) recognized as *T. tenuifolium* but its taxonomic identity needs further research. Other sampled species placed by Hendrych in *T.* ser. *Alpina* are *T. australe* R.Br. and *T. chinense* Turcz.; however, molecular data indicate they are distinct, thus making the series polyphyletic (see discussion of Clade 21).

*Thesium auriculatum* is a morphologically well-characterized species of the Dinaric Alps from Bosnia and Herzegovina to Albania which together with the Carpathian endemic *T. kernerianum* Simonk. (not sampled) constitutes Hendrych’s *T.* ser. *Auriculata*. The inclusion of both species in the series, although differing in other morphological features, was based on the presence of swollen glands between the corolla lobes (Hendrych, 1963a). Considering the phylogeny of the genus as a whole, this character lacks phylogenetic value since it is also present in many African species but is less common in the Eurasian ones (present but not so developed in e.g. *T. catalaunicum* Pedrol & M.Laínz or *T. brachyphyllum*). Some populations from the Northen Appennines identified as *T. sommieri* Hendrych (*T.* ser. *Micrantha*) present developed glands very similar to those of *T. auriculatum*.

**16. *Linophylla-Alatavica* Clade (BIPP 0.99). –** This highly-supported (Fig. 2A) clade is resolved only with ITS. It spans clades 17 through 25 and includes species with mostly European and Mediterranean distribution (Clade 17) as well as mostly Asian species in Clade 18.

**17. *Linophylla* Clade (BIPP 1). –** This clade includes *Thesium linophyllon* (=*T. linifolium* Schrank), *T. divaricatum* Jan ex Mert. & W.D.J.Koch, and *T. humifusum* DC. (Fig. 3L), three common mostly European species with branched, racemose to paniculate inflorescences. The lateral cymules normally bear 1--5 flowers and the floral bract is free from to totally adnate to the peduncle. Hendrych (1972) placed the species in *T.* ser. *Divaricata* Hendrych (*T. divaricatum* and *T. humifusum*) and *T.* ser. *Linophylla* (*T. linophyllon*). The Asian species included by Hendrych (1972) in both of these polyphyletic series are resolved in ITS Clades 24 and 25. *Thesium linophyllon* is a very variable species, especially in leaf and flower size. Some specimens from southern Europe have narrower and thicker leaves and smaller flowers, more similar to *T. divaricatum* and *T. humifusum*. The specimens with broader leaves are often confused with *T. bavarum* and are thus considered a subspecies (*T. linophyllon* subsp. *montanum* (Ehrh. ex Hoffm.) Čelak), however, our results support recognizing two different species. *Thesium divaricatum* and *T. humifusum* have been frequently treated as two different species or *T. divaricatum* as a subspecies of *T. humifusum* (e.g. Hendrych, 1972), but the characters used to separate them, such as the plant height or whether or not the bracts and upper branches are scabrid, are variable; they are probably the same species, in agreement with Pedrol & Laínz (1997). Finally, the unsampled *T. italicum* A.DC., a stoloniferous perennial from the Sardinian mountains, was placed in series *Divaricata* by Hendrych (1972) and its morphological features suggest that it might be a member of this clade.

**18. Second Asian Radiation (BIPP 0.94). –** ITS resolved this clade (Fig. 2A) with good support whereas the highest support was received with chloroplast markers (BIPP 1) but in this case excluding subclade 23 and a few other species (see discussion below). Most of the species resolved in this clade have their main areas of distribution in E Europe, N, C and E Asia and the only species present in Australia (*Thesium australe*, Clade 21). Exceptions include a few of European species that are also resolved here. The most remarkable is the NE Spanish endemic *T. catalaunicum* (Fig. 3M) which is resolved with good support as sister to the rest of species with ITS but was not supported with chloroplast markers. This species presents a very characteristic spicate inflorescence of tubular flowers and slender rhizomes. Our phylogenies do not suggest close relatives in Europe except for those in Clade 19.

**19. *Rostratum* Clade (BIPP 0.78).** This clade received low support (Fig. 2A) and is composed of a polytomy of eight clades, most of which individually received high support, as well as several species with no clear relationships on the ITS tree. This clade is a heterogeneous assemblage of taxa from 11 of the 19 series proposed for *Thesium* sect. *Thesium* by Hendrych (1972). We sampled at least 20 of the 37 species included in these series. One of the few European species is *T. rostratum* Mert. & W.D.J.Koch (Fig. 3N), represented in our sampling by two accessions from Italy. The most evident character for this species is its lack of bracteoles, a feature it shares with *T. ebracteatum* Hayne. The lack of bracteoles is probably exclusive to these two species, but it is a morphological convergence according to the ITS topology. In contrast, the combined chloroplast trees resolve these two species as sister with moderate support (BIPP 0.90) and this clade does not include *T. multicaule* Ledeb. (see below, Clade 20). Other morphological characters, like flower morphology, type of rootstock, and the relative size of the corolla lobe remnant and achene portion of the fruit, differ between *T. ebracteatum* and *T. rostratum*.

*Thesium ramosoides* Hendrych, included in *T.* ser. *Divaricata*, is a plant with numerous erect branches from a thick or branched rootstock, racemes or paniculate inflorescences, campanulate flowers and fruits with longitudinal nerves. This species is well-supported as part of the second diversification of the genus in Asia but with unresolved relationships.

However, our chloroplast trees resolved the species in a well-supported clade composed mostly of European species (except for Clade 23; see below). The chloroplast relationship is remarkable because *T. ramosoides* is endemic to Sichuan and Yunnan provinces of China (Nianhe & Gilbert, 2003).

*Thesium repens* Ledeb. is a common species in C Siberia and N Mongolia characterized by the slender rhizomes, short and thin unbranched stems and campanulate flowers in racemose inflorescences. With ITS it is not resolved as closely related to other species in Clade 19 but chloroplast markers highly support its relationship with the mostly Siberian species of Clade 22 (see below). According to Hendrych (1972), *Thesium* ser. *Repentia* Bobrov contained *T. repens* and the unsampled *T. oreogetum* Hendrych. Although *T. ebracteatum* resembles *T. repens* morphologically and was classified in this series by Hendrych (1969), ITS does not show a close relationship. Two other unsampled species in series *Repentia*, *T. hookeri* Hendrych, and *T. afghanicum* Hendrych, may reside here or in Clade 24.

**20. *Multicaule* Clade (BIPP 1).** – This clade (Fig. 2A) contains *Thesium ebracteatum* Hayne (*T.* ser. *Repentia*) from Russia and three accessions of *T. multicaule* (*T.* ser. *Caespitosa* Bobrov), all from Kazakhstan. This relationship is surprising and not immediately evident from morphology. *Thesium multicaule* is a plant of the steppes of western Siberia, Central Asia and Mongolia whereas *T. ebracteatum* (Fig. 3O) is a forest or forest margins plant from west of the Urals to Central Europe (Hendrych, 1969). They share unbranched racemose inflorescences with flowers borne on long, recaulescent peduncles, shortly tubular, campanulate flowers, and longitudinally veined fruits that are longer than the corolla remnant. Vegetatively, *T. ebracteatum* has long rhizomes with a few stems, whereas the rootstock in *T. multicaule* is thick and frequently woody, with numerous tufted stems, a character more common in tropical African species but not in the Eurasian ones. In addition, *T. multicaule* generally has bracts shorter than flowers and fruits whereas in *T. ebracteatum* they are always clearly longer. Morphologically, *T. ebracteatum* appears closer to species like *T. repens* or *T. alatavicum* Kar. & Kir., and for that reason they were included by Bobrov (1936) in his *T.* ser. *Repentia*. The chloroplast phylogeny placed *T. ebracteatum* with *T. rostratum* in a clade of mostly European species.

**21. *Australe* Clade (BIPP 0.56).** – This clade (Fig. 2A) received very low support with ITS but obtained high support with chloroplast sequences (BIPP 0.97). Two species with fused corolla lobes, *Thesium chinense* (Fig. 3P) and *T. australe*, are resolved in a well-supported subclade with *T. longiflorum* Hand.-Mazz. as sister. Both morphology and ITS sequences of *T. australe* and *T. chinense* are very similar; therefore, these two species probably should be combined under *T. australe*. Morphologically, they resemble *T. alpinum* (Clade 15) and for that reason were included in *T.* ser. *Alpina* by Hendrych (1972); however, our molecular data suggest they are not closely related. In contrast, Bobrov (1936) created the monotypic *T.* ser. *Decurrentia* Bobrov including only *T. chinense* suggesting only an outwardly resemblance to *T. alpinum*. The reticulate fruits and pentamerous flowers of *T. australe* (and *T. chinense*) are the most evident characters to differentiate them from *T. alpinum*. The Chinese endemic *T. longiflorum* (not to be confused with *T. longifolium*, Clade 22), with large tubular flowers and subreticulate fruits, was included by Hendrych (1972) in *T.* ser. *Rostrata* Bobrov. One of the characteristic features of this species is that the floral bracts on proximal flowers are not adnate to the peduncles whereas the distal ones are completely adnate (progressive recaulescence). In constrast, in most Eurasian *Thesium* species the floral bracts are adnate on all flowers. Progressive recaulescence or different degrees of fusion of the bract to the peduncle in the same individual can be seen in other Eurasian species such as *T. kotschyanum* (Clade 9) or *T. humifusum* (Clade 19) and African species with branched inflorescences (see Clade 59 below).

No sequence data currently exist for the other species of *T.* ser. *Alpina*: *T. cathaicum* Hendrych (China), *T. thomsonii* Hendrych, and *T. unicaule* Haines (both N India). Based solely on morphology these three species will likely be placed in this clade. Apparently related to both *T. australe* and *T. chinense*, they share the more or less tubular flowers in racemes and reticulate fruits with the remnant corolla about as long or longer than the gynoecial portion of the fruit.

**22. *Longifolia* Clade (BIPP 1).** – This clade (Fig. 2A) contains the species of *T.* ser. *Longifolia* Bobrov, and our data supports the monophyly of the series as proposed by Bobrov (1936) including the mostly Siberian species *T. longifolium* Turcz., *T. refractum* C.A.Mey. (Fig. 3Q), and *T. saxatile* Turcz. Hendrych (1972), however, considered *T. saxatile* in the polyphyletic *T.* ser. *Saxatilia* together with *T. pyrenaicum* (clade 15) and *T. rupestre* Ledeb. (see below). Our results do not support this division. In this clade is also resolved *T. tuvense* Kransob., a species later described from the Republic of Tuva of the Russian Federation (Krasnoborov, 1992).

Species in the *Longifolia* Clade have several morphological features in common. All of them have paniculate inflorescences on stems developing from compact rootstocks. The lateral racemes of the panicles have relatively long and wide bracts that are free or partially adnate. The flowers have a well-developed tube (shorter in *T. refractum*) and the fruits have five prominent ribs with longitudinal nerves between them. The differences are basically found on the disposition of the fruiting peduncles on the rachis (spreading to appressed) and flower and fruit size. *Thesium refractum* has the largest distributional area of this group, being widespread from Central Asia to Siberia, Mongolia, and China. According to Bobrov (1936), the eastern populations have larger flowers and are sometimes difficult to distinguish from *T. saxatile*, a species with a central and eastern Siberian distribution. *Thesium longifolium* from E Siberia and N Mongolia is easy to distinguish from *T. refractum* when fruiting, but both taxa share a number of morphological features. Psammophyte populations in NE Mongolia, W Manchuria, Tuva, and Buryatia, with flexuous inflorescences, larger flowers and fruits (with very prominent ribs) were described as *T. tuvense.* Our results support the close relationship between this species and the morphologically similar *T. saxatile*. Bobrov (1936) noted the morphological differences of plants growing on sandy soils but treated them as components of the morphologically variable *T. saxatile*.

The unsampled *T. rupestre*, known from the Altai, is probably related to species in this clade, although the few specimens available show inflorescences in racemes, not panicles. Diagnostic characteristics of this species are its long external glands and the accrescent hypanthium. *Thesium brevibracteatum* P.C.Tam (nom. illeg.), a species from Inner Mongolia (China), was described in the protologue as morphologically close to *T. longifolium* and might also be included in this clade. A new name, *T. longiperianthium* H.H.Xu & W.Jun was proposed for this species (Xu & al., 2020). *Thesium basninianum* Turcz., described from the Transbaikal, although considered as a synonym of *T. chinense* by Bobrov (1936), is morphologically similar to *T. longifolium* but with bracts much longer than fruits.

**23. *Ramosum* Clade (BIPP 1).** – This clade contains eight accessions, most of which were identified as *Thesium ramosum* (Fig. 3R), a widespread species from the Balkans to Central Asia. This taxon has frequently been called *T. arvense* (Hendrych, 1961, 1993) but as pointed out by Gutermann (2009) that name is superfluous and therefore illegitimate. Endemic to Europe (Bulgaria to Crimea), *T. moesiacum* Velen. and *T. simplex* Velen. were accepted by Hendrych (1993) as perennial subspecies of the annual *T. dollineri* Murb. We did not sample the latter, but given that these taxa and *T. ramosum* have identical ITS sequences, all might best be considered one species (*T. ramosum*) as proposed by Bobrov (1936). This group is morphologically more diverse in Europe and the Caucasus than the Asian populations, which are morphologically stable in spite of a wider area of distribution (Bobrov, 1936). Characters that would separate *T. ramosum* from *T. dollineri* s.l. are the degree of inflorescence branching and the relative length of flowers and bracts, characters of taxonomic importance but of low phylogenetic value. Our accession 5558 from USA (Montana) was identified as *T. ramosum*. This species was accidentally introduced into North America and now has an expanding range that includes North Dakota, Montana, Idaho, and Alberta (Nickrent, 2016; Macdonald & Visser, 2022). *Thesium asperulum* Boiss & Buhse and *T. laxiflorum* Trautv., both described from the Caucasus, were considered forms of *T. ramosum* by Bobrov (1936). The unsampled *T. carduchorum* Hedge from Turkey shares morphological features with *T. ramosum* and is probably in the range of morphological variation of this species.

There is a remarkable incongruence in the phylogenetic position of this clade between the ITS and chloroplast phylogenies. Whereas ITS resolved it within Clade 18, chloroplast data placed it within a well-supported clade of mostly European species. Further research is necessary to explain this incongruence, but one hypothesis is that *T. ramosum* s.l. (including the subspecies of *T. dollineri*) originated as a result of hybridization between one European (maternal) and one Asian (paternal) species in Eastern Europe followed by ITS concerted evolution towards the paternal progenitor.

**24. *Himalensia* Clade (BIPP 1).** – This clade (Fig. 2A) was recovered with the highest support with ITS and chloroplast sequences. It includes *Thesium gontscharovii* Bobrov and *T. ramosissimum* Bobrov of mountainous regions of Tajikistan and Kyrgyzstan, together as sister to *T. himalense* (Fig. 3S) from S China, NW India, and Nepal. Both *T. gontscharovii* (*T.* ser. *Divaricata*) and *T. ramosissimum* (*T.* ser. *Caespitosa*) have a woody rootstock with numerous ascending to erect branched stems and campanulate flowers in slender racemes. They differ from each other basically in the sizes of bracts, bracteoles, flowers and fruits. *Thesium himalense*, however, has slender, scarcely branched procumbent stems. It shares with the previous species the campanulate flowers and fruits with numerous longitudinal veins.

Unsampled species include *Thesium pachyrhizum* A.DC. (*T.* ser. *Himalensia* Hendrych), *T. hookeri* Hendrych, *T. afghanicum* Hendrych (both in *T.* ser. *Repentia*) and *T. indicum* Hendrych (*T.* ser. *Indica*) which are all morphologically similar to *T. himalense*.

**25. *Alatavicum* Clade (BIPP 0.99).** – This highly-supported clade (Fig. 2A) includes all five accessions of the Central Asian *Thesium alatavicum* (Fig. 3T) as well as *T. ferganense* Bobrov, a species described from near the Fergana Valley, within the area of distribution of *T. alatavicum*. It is interesting that the four accessions of *T. alatavicum* from which we obtained chloroplast sequences were not resolved as monophyletic on the chloroplast trees, although they are morphologically similar. *Thesium alatavicum* is easily identified by the stoloniferous rhizomes, the one-sided (secund) arrangement of leaves and flowers, and racemose inflorescences in which the floral bracts are markedly longer than the flowers proximally but progressively shorter distally. *Thesium ferganense*, included in *T.* ser. *Linophylla* (Clade 17) by Hendrych (1972), however, has many stems that arise from a compact rootstock and paniculate inflorescences. Although they share some morphological characters with *T. linophyllon,* they are distantly related.

Unsampled species include *Thesium brevibracteatum* Sumnev., which was described from the Tashkent Alatau Mts., and was included by Hendrych (1972) in *T.* ser. *Alatavica* Hendrych. The type specimen has secund inflorescences as in *T. alatavicum*; however, it was considered by Goloskokov (1960) as a southern race of *T. ramosum*.

**26. African Clade A (BIPP 1).** – This clade (Fig. 2B) contains 175--182 species, most of them of African origin. Those species currently known from South America, Madagascar, and SE Asia (*Thesium psilotoides*) have resulted from long-distance dispersal from Africa (see below). The early diverging member of this clade is *Thesium nautimontanum* (García & al., 2018), an endemic to the Matroosberg Mts. of South Africa (Fig. 4A). The phylogenetic position of *T. nautimontanum* as sister to the rest of African species is not recovered with the chloroplast phylogeny that resolve Clades 27--30 as the earlier divergent lineages (see discussion below). Based on their ITS analysis, Zhigila & al. (2020) erroneously assigned this species to subgenus *Frisea*.

**Fig. 4.**
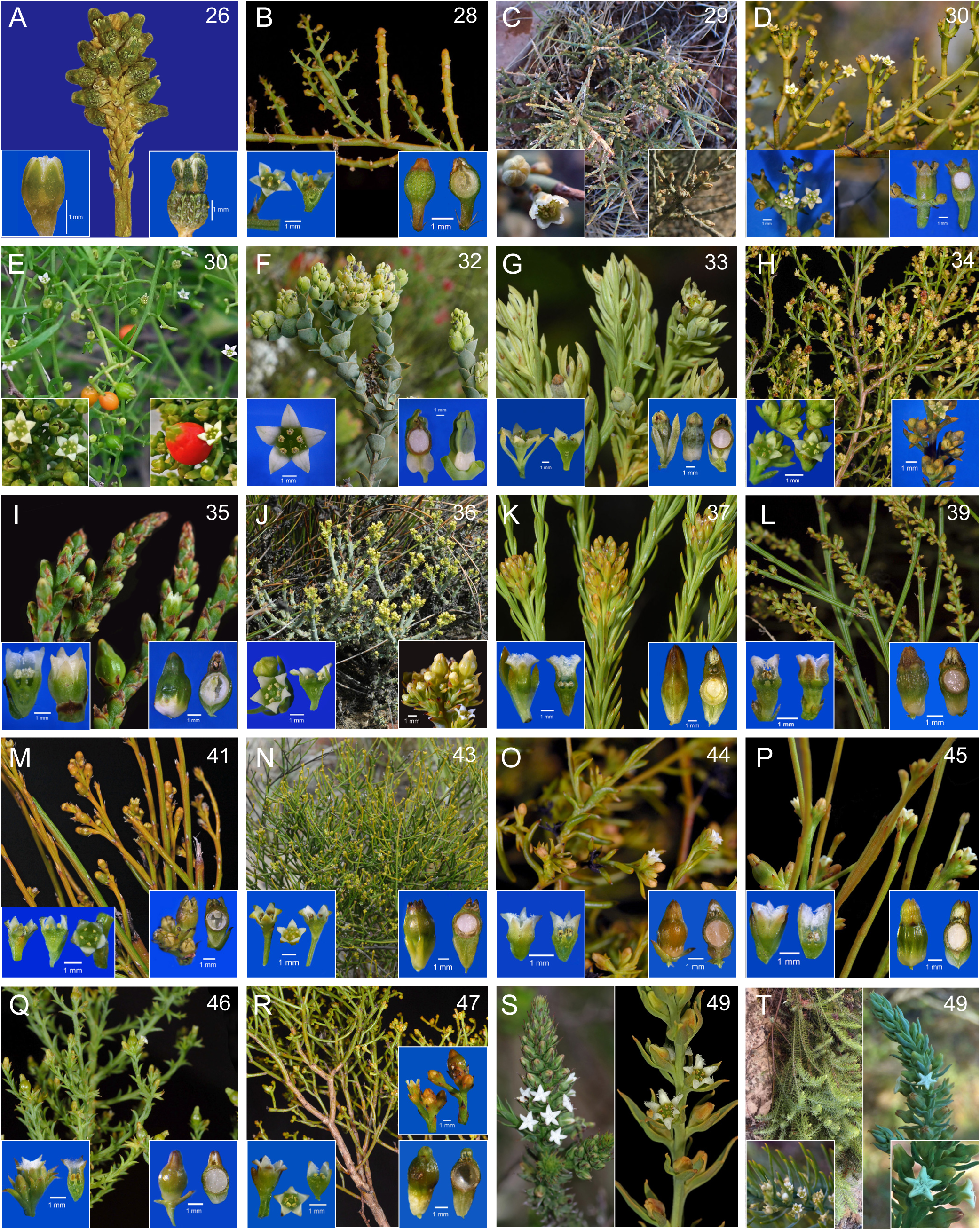
Photographs of representatives of the major clades of mostly South African *Thesium* species. The numbers in the upper right corners correspond to clade numbers on the Bayesian trees (Fig. 2B-C). L.S = longitudinal section. **A,** African clade, *T. nautimontanum*, rehydrated herbarium specimens showing flower and young fruits (*Šibík & Šibíková MS463*, MA); **B,** *Spinosa* clade, *T. spinulosum*, shoot with flowers and fruits, flower and L.S., fruit L.S.; **C,** *Namaquense* clade, *T. lacinulatum*, habit, flower and fruits; **D,** *Triflorum* clade, *T. scandens*, flowering shoots, flowers, and fruit L.S.; **E,** *Triflorum* clade, *T. triflorum* ; **F,** Core Cape clade, *T. euphorbioides*, flowering shoots, flowers, fand ruit L.S.; **G,** *Foliosum* clade, *T. foliosum*, flowering shoots, flower L.S., and fruit L.S.; **H,** *Ericaefolium* clade, *T. ericaefolium,* flowering shoots, flowers, and fruits; **I,** *Scirpioides* clade, *T. flexuosum*, flowering shoots, flower L.S., and fruit L.S.; **J,** *Strictum* clade, *T. albomontanum*, habit, flower L.S., and fruits; **K,** *Capitatum* clade, *T. carinatum,* shoots with young fruits, flower L.S., and fruit L.S.; **L,** *Frisea* clade, *T. funale*, shoots with flowers and young fruits, flower L.S., and fruit L.S; **M,** *Nigromontanum* clade, *T. nigromontanum*, shoots with flowers and young fruits, flower L.S., and fruit L.S.; **N,** *Virgatum* clade, *T. pseudovirgatum*, habit, flower L.S., and fruit L.S.; **O,** *Capitellatum* clade, *T. prostratum*, flowering shoots, flower L.S., and fruit L.S.; **P,** *Acuminatum* clade, *T. capituliflorum*, flowering shoots, flower L.S., and fruit L.S.; **Q,** *Hispidulum* clade, *T. hispidulum*, flowering shoots, flower L.S., and fruit L.S.; **R,** *Commutatum* clade, *T. commutatum*, shoots with flowers and young fruits, flower L.S., and fruit L.S.; **S,** *Gnidaceum* clade, *T. gnidiaceum* (left); *T. impeditum* (right), inflorescences; **T,** *Gnidaceum* clade, *T. oresigenum* (left) showing pendant habit and flowers; *T. phyllostachyum* (right) inflorescence and flower close-up.

**27. African Clade B (BIPP 1).** – This clade (Fig. 2B) contains all members of Clade 26 except *Thesium nautimontanum*. It contains three major clades (28, 29/30, and 31) that arise from a polytomy. Clades 28--30 includes species of *T.* subgen. *Discothesium* and Clade 31 of *T.* subgen. *Frisea*.

**28. *Spinosum* Clade (BIPP 1).** – A clade composed of *Thesium spinosum* and *T. spinulosum* (Fig. 4B) was obtained from both ITS (Fig. 2B) and the chloroplast spacer data (individually and combined). These species both occur in sandy soil areas in the north and west portion of the CFR of South Africa. The inflorescences are indeterminate spikes or racemes with pedicellate, beardless, campanulate flowers. Flowers are born in the axils of bracts that are robust and spiny in *T. spinosum*, whereas in *T. spinulosum* bracts and bracteoles are very thin and subulate. Normally the flowers are solitary but in *T. spinosum* they may be accompanied by secondary flowers in small dichasia or with flower buds that do not develop into mature flowers but retain the minute ovate and ciliolate bracteoles. In *T. pungens* A.W.Hill and *T. spinosum*, all vegetative parts are moderately succulent. The leaves and bracts subtending flowers have swollen bases but with age the tips senesce and become hardened spines. For some shoots in *T. spinosum* this process goes to an extreme such that the bract bases and even part of the stem can become quite inflated. *Thesium spinulosum* shows a marked stem dimorphism (Fig. 4B) where flowering shoots are generally less succulent than vegetative ones. This dimorphism may be a response to environmental conditions such as dry, sandy and/or salty soils. There is a clear tendency to succulent vegetative organs in species or populations in coastal and/or sandy environments and is frequently observed e.g. in members of sect. *Frisea* (Clade 39), *T. subsucculentum* (Clade 4) or in coastal populations of *T. triflorum* (Clade 30), among many other examples.

Sonder (1857) placed *T. spinosum* in group b “spinosa”. A. De Candolle (1857) named *T. spinulosum* but apparently this was unknown to Sonder, hence he did not list this species. Hill (1915) placed the two species together in *Thesium* sect. *Imberbia* A.W.Hill, subsect. *Subglabra* A.W.Hill (along with 71 other species), but their proximity in his key indicates he considered them related. In the molecular phylogenetic study of Moore & al. (2010), a collection identified as *T. spinosum* was sister to *T. virgatum* Lam., which is clearly erroneous. Unsampled taxa likely to be components of the Spinosum clade includes *T. pungens* and *T*. *aristatum* Schltr. which is here considered a synonym of *T. spinulosum*.

**29. *Namaquense* Clade (BIPP 1).** – Clades 29 and 30 (Fig. 2B) were only weakly supported as sister using ITS data, whereas in the trees from concatenated *trnDT* and *trnLF* trees this relationhip was resolved with very high or the highest BIPP, MPBS and MLBS values (1.0, 97 and 99, respectively, Figs. S4--S7). Clade 29 includes drought-tolerant species of arid areas of Namibia, Namaqualand (western South Africa), Great Karoo semi-desert and the southern Kalahari arid savanna. Most members of the clade are shrubs with spiny stems. The flowers are sessile in spike-like inflorescences but, as in *Thesium spinosum*, there can be additional flowers in each axil or numerous short axillary branches developing flowers in congested clusters with internodes elongating in fruit (as in *T. namaquense* Schltr. or the unsampled *T. archeri* Compton and *T. whitehillense* Compton). Most species in this clade have fringed corolla lobe margins with different types of projections, varying from hairs to membranous flaps (the latter reminiscent of the auricles of the Eurasian species). One exception is *T. namaquense*, which has flowers without any projections on the margins of the corolla lobes and for that reason Hill (1915a, 1915b) placed it in *T.* subsect. Subglabra. *Thesium* 4393, is probably a new species of a small shrub with flexuous and minutely scabrous stems, fleshy leaves and bracts with spiny tips, flowers with somewhat developed marginal flaps in spike-like inflorescences. Whereas our ITS phylogeny places this species in this clade, the chloroplast sequences do not support this relationship but highly support it as belonging to a clade that includes ITS Clades 29 and 30.

*Thesium hystrix* A.W.Hill and *T. lacinulatum* A.W.Hill (Fig. 4C) were included by Hill (1915a) in *T.* subsect. *Fimbriata* along with other spinescent species such as *T. horridum* Pilg., *T. hystricoides* A.W.Hill and *T. pleuroloma* A.W.Hill. *Thesium cruciatum* A.W. Hill was later included in *T.* subsect. *Fimbriata* (Hill, 1915b) suggesting a close relationship with *T. lacinulatum*, as is confirmed by our results. *Thesium xerophyticum* A.W.Hill was not included in Hill (1915a, 1915b), possibly because it was described from Namibia and considered tropical Africa. The sectional name *Fimbriata* may be confusing because *T. fimbriatum* A.W.Hill is a tropical East African species not closely related (see Clade 55), thus this name was not used here.

Unsampled taxa include *T. archeri* (Little Karoo), *T. dissitiflorum* Schltr., *T. horridum*, *T. hystricoides* A.W.Hill (Kalahari), *T. pleuroloma* A.W.Hill, and *T. whitehillense*. *Thesium rigidum* Sond., although sometimes considered a synonym of *Lacomucinaea lineata* (or *T. lineatum*), is a different plant, with scaly leaves, spiny branches, and flowers lacking any projections on the margins of the corolla lobes. It was described from north of Winterhoek Mts. of the Eastern Cape and it might be resolved in this clade.

**30. *Triflorum* Clade (BIPP 1).** – Receiving the highest support, this clade (Fig. 2B) is composed of three species, *Thesium galioides*, *T. scandens* (Fig. 4D), and *T. triflorum* (Fig. 4E), that were all placed in *T.* subsect. *Subglabra* by Hill (1915a), albeit with many other species. Both ITS and chloroplast data indicate these three species are closely related, as suggested by Hilliard (2006). These plants are scandent herbs or subshrubs with inflorescences in indeterminate thyrses whose lateral cymules have a variable number of flowers (mostly 1--7). The floral bracts are mostly non-recaulescence, but in *T. triflorum* the bract may be adnate to about half of the length of the peduncle. The most widespread species is *T. triflorum*, one of the few species in the genus with fleshy fruits (Fig. 4E). According to Brummitt (1976), its distribution is disjunct in tropical East Africa (Tanzania and Nyika Plateau) and from S Mozambique to the E and S of South Africa. The tropical African populations have some morphological and ecological differences with those from South Africa (Brummitt, 1976). This relationship and the position on the tree indicate that the tropical African populations have their origin in South Africa and the fleshy red fruits contributed to its dispersal, probably by birds, north of its original area. *Thesium triflorum* is very variable especially in terms of leaves and bracts morphology (from linear to wide elliptic, acute to obtuse) and fleshiness of vegetative organs. The other two species, *T. scandens* and *T. galioides*, have more restricted distributions and are basically plants of the Fynbos (a species-rich scrub biome on nutrient-poor soils). The characters used to separate these two species are flower and bract lengths; however, the morphology of leaves and bracts may change with age, as occurs frequently in *Thesium*. Younger plants and branches may have well-developed leaves that are lost with age, thus this variable character might not have value to recognize two species.

In the ITS phylogeny, two collections (5300, 5309) from the Anysberg and Touwsberg Mts. in the Western Cape Province (South Africa) are resolved as sister to these three species. These plants, that might belong to a new species, show similar inflorescence structure, but the lateral cymes are much reduced, lacking well-developed bracts and leaves. Chloroplast markers resolve both accessions within the ITS Cape Clade (31), showing a remarkable topological incongruence with the nuclear phylogeny.

Unsampled species might include *T. semotum* N.E.Br., an enigmatic species that was described from a single collection presumably from the former Transvaal (northern provinces of South Africa). The type is a young specimen with flowers in compact lateral cymes of three or more flowers with the apparently succulent basal bract longer than young cymules. This taxon might be an immature *T. triflorum*, which is also present in the area, or more probably a species that has not been further collected.

**31. *Frisea*-*Psilotoides* Clade (BIPP 1).** – This large and well-supported clade (Figs. 2B and 2C) is composed of two large clades, 32 (*T.* subgen. *Frisea*) and 48 (*T.* subgen. *Psilotoides*).

**32. Core Cape Clade. *Thesium* subgen. *Frisea* (BIPP 1).** – Although this clade received the highest support (Figs. 2B and 2C), it is composed of a large polytomy of clades (33--38) representing the radiation of the genus that gave rise to most of the Cape diversity of *Thesium*. All the species have sulcate stems with decurrent leaves. Anatomically, the primary phloem fibers are located deep in the cortex near the vascular cambium and are surrounded by the chlorenchyma from the fused portion of the leaves (Nickrent & García, 2015). This character is almost exclusive to this clade with some exceptions such as the east tropical *T. kilimandscharicum* Engl. in Clade 68 (Polhill, 2005; Hilliard, 2006).

Probably the only species that lacks sulcate stems in this clade is *T. euphorbioides* P.J.Bergius (Fig. 4F), a very characteristic plant for its large size, broad, cordate, thick leaves and bracts, and flowers in terminal elongated cymes with the lateral cymules protected by the basal bracts. The habit is similar to *T. strictum* (see below) and the fruits have a characteristic swollen pedicel like in that species. Because ITS places this species in the polytomy of Clade 32, further resolution may or may not confirm a relationship between *T. euphorbioides* and *T. strictum*. Chloroplast sequences resolve this species as sister to the Cape Clade (Figs. S4--S7) excluding *T.* sect. *Frisea* (Clade 39).

Within this clade other species have uncertain phylogenetic position. It is surprising that the individual identified as *T. rufescens* A.W.Hill is not resolved in any of the Cape clades with ITS. This species belongs to *T.* sect. *Barbata* (Hill, 1915a, 1915b) and morphologically might be confused with *T. pubescens* A.DC. (Clade 37) as both are densely pubescent and have similar inflorescences although in *T. rufescens* cymes are spikate (see further information on this species in Lombard & Le Roux, 2023). Chloroplast data place this species in a clade related to *T. flexuosum* A.DC. and one of the two accessions of *T. sonderianum* Schltr., two Cape species that are not closely related on the ITS trees.

It should be mentioned here that Hill’s classification recognized four sections within the group of species resolved in this clade: sects. *Annulata* (≡ *Frisea*)*, Barbata*, *Penicillata,* and *Imberbia*. *Barbata* is characterized by having a dense beard of hairs that descend from the apices of the corolla lobes. Sections *Frisea* and *Penicillata* also have such hairs and, according to our ITS tree topology, *Frisea* is a subclade within section *Barbata*. These bearded flowers occur in clades ranging from 32--80 whereas flowers without beards in *T.* sect. *Imberbia* occur in our Clades 28--80. This clearly demonstrates that the presence or absence of an apical beard has little to no phylogenetic significance.

**33. *Foliosum* Clade (BIPP 1).** – Clade 33 (Fig. 2B) includes *Thesium foliosum* A.DC. (Fig. 4G) and *T. fruticosum* A.W.Hill, the latter with short and scattered leaves and bracts shorter than flowers and the former species with dense leafy bracts that are longer than the flowers. Both are erect shrubs and have cucullate corolla lobes with papillose margins even though they were included in *T.* subsect. *Subglabra*. In some cases, as for example in the voucher specimen 4706 of *T. fruticosum*, the apex of the corolla lobes has more developed papillae and resemble a short beard. *Thesium foliosum* has ITS sequences with some polymorphic sites, not found in *T. fruticosum*.

Unsampled species include *T. susannae* A.W.Hill, which is very similar to *T. foliosum* in habit, vegetative features, flowers and fruits, however, the inflorescence has flowers in short lateral cymules rather than solitary. If this is merely a developmental difference these two taxa may be the same species.

**34. *Ericaefolium* Clade (BIPP 1).** – Clade 34 (Fig. 2B) includes six accessions of *Thesium ericaefolium* A.DC. (Fig. 4H), a leafy shrublet with small cymose inflorescences on lateral short branches and flowers that lack an apical beard on the corolla lobes (*T.* sect. *Imberbia*). The species is morphologically well characterized and resolved as monophyletic with ITS. However, the five accessions with chloroplast data are resolved in two different clades with the highest support (Figs. S7, S8).

Unsampled species include *Thesium glomeruliflorum* Sond. that differs in several vegetative characters. The axillary cymules are similar to *T. ericaefolium* but the floral bract is longer than the cymules. This species was sampled by Moore & al. (2010) and Zhigila & al. (2020) and in the combined analyses was resolved in a clade with *T. ericaefolium*. However, the individual analyses of nuclear and chloroplast sequences by Zhigila & al. (2020) resolved this species in different clades related to *T. strictum* and *T. albomontanum* Compton.

**35. *Scirpioides* Clade (BIPP 0.97).** – Our sampling includes three of the seven species recognized by Lombard & al. (2021) in their taxonomic revision of the *Thesium scirpioides* A.W.Hill species complex. All the species sampled received moderate to the highest support as monophyletic with ITS (Fig. 2B) but in the chloroplast analyses *T. flexuosum* (Fig. 4I) was not part of the *T. scirpioides* and *T. junceum* Bernh. clade. The *Scirpioides* clade includes erect plants with leaves reduced to scales (except in young specimens that may have well-developed leaves), flowers in elongated spikes, corolla lobes with woolly apical beards (*T.* sect. *Barbata*) and anthers included in the floral tube. The exception is *T. flexuosum*, a frequently straggling suffrutex with compact and branched spikes, corolla lobes with an apical beard of stiff comb-like hairs similar to many species in *T.* sect. *Frisea* (Clade 39) and anthers partially exerted from the hypanthium. The morphological differences among these species are discussed in Lombard & al. (2021). Except for *T. flexuosum*, the other species sampled are mostly distributed in eastern provinces of South Africa, Eswatini (former Swaziland) and Lesotho and *T. scirpioides* also in Mozambique (Hilliard, 2006), sharing the grassland habitat with species of Clade 62. Although the geographical distribution for some of these species is atypical for species of the Cape Clade, they have twisted placentae, a character shared with other species of the group, whereas the grassland, savanna and other tropical taxa (Clade 50) mostly have short and erect placentae (Visser & al., 2018).

Unsampled species included in this complex by Lombard & al. (2021) are *T. natalense* Sond. and the annual *T. paronychioides* Sond., both species with apical beards on the corolla lobes. A rare species from the Eastern Cape, *T. lisae-mariae* Stauffer and the more recently described *T. atratum* N.Lombard & M.M.Le Roux have similar inflorescences and vegetative characters but the corolla lobes are not bearded. *Thesium stirtonii* Zhigila & al. is another species lacking an apical beard that is probably related to species in this clade. Further molecular analyses including non-bearded species are required to test the monophyly of the species complex defined by Lombard & al. (2021). Zhigila & al. (2020) resolved *T. lisae- mariae* within the clade of *T.* sect. *Frisea* but the specimen sequenced was probably misidentified because the floral characters are very different between these two groups.

**36. *Strictum* Clade (BIPP 1).** – Clade 36 (Fig. 2B) includes *Thesium strictum*, one of the most widespread species of the Fynbos Biome (Rebelo & al., 2006). It has a distinctive growth form known as wand-plant architecture (Bailey & al., 2019), is frequently more than 1.5 meters tall and is woody at the base. The inflorescences are cymose, the corolla lobes with short marginal papillae and the fruits have swollen peduncles. There is, however, great morphological variation, especially in density and shape of the leaves and in plant robustness. The degree of compaction of the inflorescence is also highly variable in the species. At a first glance, a relationship with *T. albomontanum* is surprising. This species is a robust shrub with short and thick stems and leaves (Fig. 4J). However, the corolla lobes are also subpapillate on the margins and the inflorescence is a less branched and condensed version of the inflorescence of *T. strictum*, with a reduction in the number of flowers of partial cymes.

The specimen identified as *T. pinifolium* A.DC. was shown with ITS to be sister to the *T. strictum* and *T. albomontanum* subclade. This species has a very similar growth form but its stems are densely covered with long, terete, secund leaves. As in *T. strictum* its flowers have papillose corolla lobe margins whereas its inflorescence is sometimes very condensed. Compared to the widespread distribution of *T. strictum*, *T. pinifolium* is restricted to rocky Cape Fold Belt mountains.

Unsampled species include *T. occidentale* A.W.Hill and *T. umbelliferum* A.W.Hill. These two species are tentatively included here based on their inflorescences that are similar to *T. strictum* and *T. pinifolium*. The flowers in the former are similar to *T. strictum* as they have papillate corolla lobe margins whereas in the latter the corolla lobes bear dense beards. *Thesium penicillatum* A.W.Hill also has the apical beard and based on its growth form could be a member of the *Strictum* Clade. Support for this placement was obtained following the molecular phylogenetic analysis by Zhigila & al. (2020). It was the only species placed in *T.* sect. *Penicillata* by Hill (1915b), likely because it bears hairs behind the anthers that do not attach to the anther apices. Post-staminal hairs that connect the corolla lobes to the anther apex can be seen in many *Thesium* species of sections *Imberbia* and *Barbata*. Compton (1931) included *T. hillianum* Compton in section *Penicillata* based on its unattached anther hairs, but this plant differs from *T. penicillatum* in many other morphological characters, thus the hair feature has questionable phylogenetic value. *Thesium aspermontanum* Zhigila & al., from Skurweberg Mt of the Western Cape was resolved as related to *T. strictum* by Zhigila & al. (2020).

**37. *Capitatum* Clade (BIPP 1).** – Clade 37 (Fig. 2B) comprises leafy and slender to mat-forming shrubs or shrublets all of them in *T.* sect. *Barbata*. These species are typical Fynbos Biome plants mostly distributed in the Western Cape Province of South Africa. Some of them have a broader distribution out of that province, such as *T. imbricatum* Thunb., but they are mostly limited to fynbos vegetation areas. Their synflorescences are cymose, generally composed of 10 or more flowers but this number can be reduced in some individuals like e.g. our accession 5392 of *T. sonderianum*. It is possible to differentiate the terminal first flower of the cymose system but in those species with compact inflorescences the individual cymes cannot be discerned without dissection. Frequently, they show a sympodial growth of vegetative shoots from below the inflorescences. Depending on the growth of the floral peduncles, the inflorescences may be rounded heads as in *T. scabrum* L., corymbiform with well-developed peduncles as in *T. helichrysoides* A.W.Hill or a false spike as in *T. sonderianum*. A beard of hairs usually covers the entire inner surface the corolla lobes but is present as long papillae along the lobe margin in *T. imbricatum* and *T. sonderianum*.

The taxonomy of this group is especially difficult because many of the vegetative characters used to separate species, such as leaf and bract margins, are frequently variable. Style length is important to separate species but its infraspecific variability has not been studied. Individuals showing characters intermediate between species are very common, especially in the widespread *T. carinatum* A.DC. (Fig. 4K). Most species in this clade are glabrous but exceptions include *T. pubescens* and the more or less scabrid *T. scabrum*. Individuals of the latter species can be totally glabrous.

Unsampled species include *T. boissierianum* A.DC., *T. brachystylum* A.W.Hill, *T. ecklonianum* Sond., *T. fallax* Schltr., *T. hollandii* Compton, *T. karooicum* Compton, and *T. sawae* Zhigila & al.

**38. *Frisea*-*Commutatum* Clade (BIPP 0.6).** – This poorly supported clade (Fig. 2B) includes Clades 39 through 47 and is composed entirely of South African species.

**39. Section *Frisea* Clade (BIPP 1).** – Clade 39 (Fig. 2B) received the highest support including the species of *T.* sect. *Frisea*, a section that was renamed by Hill (1915a) as *T.* sect. *Annulata*. The latter name has been frequently used for this section, but the name *Frisea* was previously validly published at the rank of section, including the type species *T. frisea* L., and therefore has priority over *Annulata*, an illegitimate and superfluous name. The ITS phylogeny resolves this group as nested within the Core Cape Clade, whereas it is sister to the rest of species in chloroplast phylogenies. The monophyly of this group is supported by the morphological features of the flower detailed in the Introduction (free anthers, ring of golden hairs at the level of departure of the staminal filaments) and is also well-supported by chloroplast data. However, there is no internal resolution, and the clade is basically a polytomy in spite of the great variation in inflorescence types and vegetative features. The characters used to separate species, such as whether the bract margin is serrate or not, or the general morphology of the inflorescence, are frequently variable even in the same specimen (Kathleen Immelman, pers. comm.). For this reason, many of the plants in this clade are here identified only as belonging to the section but not identified to species. The inflorescences are very diverse, including elongated spike-like inflorescences, compact glomerules and cymes. Some species have true spikes and others exhibit lateral cymules on an axis with indeterminate apical growth (e.g. *T. funale* L., Fig. 4L). The great variability of the inflorescences in this section is exemplified by *T. frisea*. Even on the same specimen, inflorescences are basically cymose but the units can be branched or contracted, of which the latter can mistakenly be considered spikes.

The variability within this section is also present in vegetative structures such as prostrate to erect stems, habit of the whole plant or the size and shape of the leaves, bracts and bracteoles. The various species can also occur in very diverse habitats. The high morphological variation between and within species of *T.* sect. *Frisea* contrasts with the low internal resolution in this clade, probably as the result of a rapid and more recent diversification. In addition, the ITS sequences of many individuals present several polymorphic sites suggesting the existence of hybridization, polyploidy or other phenomena in which complete homogenization via concerted evolution has not occurred to completion.

Species in this clade include *T. aggregatum* A.W.Hill, *T. annulatum* A.W.Hill, *T. bathyschistum* Schltr., *T. brachygyne* Schltr., *T. diversifolium* Sond., *T. dmmagiae* Zhigila & al., *T. elatius* Sond., *T. frisea*, *T. funale*, *T. litoreum* Brenan, *T. macrostachyum* A.DC., *T. micropogon* A.DC., *T. neoprostratum* Zhigila & al., *T. patersonae* A.W.Hill, *T. patulum* A.W.Hill, *T. spicatum* L., *T. subnudum* Sond., and *T. urceolatum* A.W.Hill.

**40. *Nigromontanum*-*Commutatum* Clade (BIPP 1).** – Composed of Clades 41 through 47, Clade 40 (Fig. 2C) received the highest support in all ITS analyses. It includes fynbos species mostly from the Western Cape Province, less commonly from the Eastern and Northern Cape Provinces of South Africa. A common character for most of the species in this clade is that partial inflorescences bear few (up to 5) flowers, but this feature cannot be considered as a synapomorphy of the clade. A trend is observed towards the reduction in the number of flowers on the partial cymes and in some cases only the terminal flower is developed (e.g. *T. euphrasioides* A.DC., Clade 46). There is a high diversity in floral features, including species of Hill’s sections *Imberbia* and *Barbata* and taxa with intermediate features between both sections (see below).

**41. *Nigromontanum* Clade (BIPP 0.95).** – Clade 41 (Fig. 2C) received high support from Bayesian analysis but bootstrap values for MP and ML were <50%. Two species of *T.* sect. *Barbata* are successively sister to a well-supported clade of species of *T.* sect. *Imberbia*. Accession 5567 is a plant that has some features of *T. capitellatum* A.DC. but does not fully match the description. This and its position on the ITS tree suggest it is a new species. The other *Barbata* species, *T. microcephalum* A.W.Hill, is an endemic small shrub growing at high elevations (elev. ca. 2,000 m) within the Hex River Mts (Matroosberg). It is another Matroosberg endemic whose closest relationship is not well resolved in this study. As other species in Clade 40, the inflorescences in *T. microcephalum* are few flowered.

The remaining portion of Clade 41 received the highest support with both ITS and chloroplast data. It includes *T. corymbuligerum* Sond., two morphologically similar species, *T. nigromontanum* Sond. (Fig. 4M) and *T. leptocaule* Sond. as well as one unidentified species (5393). *Thesium leptocaule* has visible glands between the corolla lobes that are missing in *T. nigromontanum* (Hill 1915a, 1915b). The presence of these glands, however, is not always easy to observe and our results suggest it lacks phylogenetic value in this group. Flowers are arranged in short, mostly lateral cymose inflorescences of 3--5 flowers, and the bracts have finely serrate margins, frequently with blackened tips. Species in this group show similar floral and fruit characteristics but are very different vegetatively. The stems and leaves in *T. corymbuligerum* (5419, 5433 and 5525), a species sometimes considered as a synonym of *T. virgatum* (Clade 43), are thicker and the leaves have spreading and subspiny tips, especially in younger specimens. However, *T. sp. nov.* 5393 has slender and thin stems, and the very reduced leaves are almost completely appressed to the stems. This latter accession is the only one in clade 41 with polymorphic ITS and the ribotypes recovered after cloning are not monophyletic (i.e. resolved in two distinct subclades).

Unsampled species include *T. nigroperianthum* Zhigila & al., *T. rhizomatum* Zhigila & al. and *T*. *crassifolium* Sond. (=*T. sedifolium* A.DC.). The latter is a small, erect and densely leafy shrublet with stout stems, short succulent leaves and bracts, and solitary flowers in spike-like inflorescences that are shorter than the bracts and bracteoles. It shares flower and fruit characteristics of other species in the clade including leaves and bracts with blackened tips.

**42. *Virgatum*-*Commutatum* Clade (BIPP 0.99).** – Clade 42 (Fig. 2C) consists of Clades 43 through 47 with species formerly classified in sections *Barbata* and *Imberbia*. Clade 43 contains species of *T.* sect. *Imberbia* subsect. *Subglabra* whereas Clade 44 contains species of sect. *Barbata*. The exception to this is *T. commutatum* Sond., which Hill (1915a) placed in sect. *Imberbia*, but is here resolved in Clade 47. The species in Clade 42 show different levels of apical trichome development on the corolla lobes. Clade 43 species lack hairs (e.g. *T. quinqueflorum* Sond.) or have sparse marginal hairs. Clade 44 species range from sparse marginal (e.g. *T. commutatum*), to dense marginal (e.g. *T. selagineum* A.DC.) to dense and long hairs that cover the inner surface of the corolla lobes (e.g. *T. cuspidatum* Sond.). Within Clade 44 these trichome level categories do not map to specific subclades.

**43. *Virgatum* Clade (BIPP 1).** – Clade 43 (Fig. 2C) includes, together with other species, several accessions of *T. virgatum*, which like *T. strictum*, is a very widespread and common species (or species complex) in the Cape. This clade illustrates the great variation in vegetative features of closely related taxa in the genus. Habits range from herbs with wiry branches (e.g. *T. juncifolium* A.DC.) to robust and more or less stiff shrubs with lignified stem bases (e.g. *T. pseudovirgatum* Levyns, Fig. 4N). Vegetative variation also affects leaf development, thereby giving very different general aspects to plants. For example, species like *T. virgatum* or *T. nudicaule* A.W.Hill have leaves reduced to scales whereas *T. quinqueflorum* is a densely leafy shrub. Also variable at the species level is the relative length of bracts and flowers/fruits. The species in this clade have corolla lobes with cucullate apices, varying in development and color, but always present. There is variation in flower shape from almost rotate (*T. pseudovirgatum*, *T. quinqueflorum*) to tubular (*T. juncifolium*), style length, and corolla lobe margins (from entire to fimbriate). In general, the fruits present five (rarely more) conspicuous swollen ribs at the base that typically extend to the peduncle (Fig. 4N, right inset), a character rarely present in species of other clades. The swollen ribs are present but not as developed in *T. juncifolium* and are lacking only in accession 5377, a possible new species.

Unsampled species include *T. affine* Schltr. (probably a synonym of *T. quinqueflorum*) and *T. schumannianum* Schltr. The latter has developed leaves (but not as densely leafy as *T. quinqueflorum*) and compact inflorescences with flower and fruit characters similar to other species of the clade.

**44. *Capitellatum* Clade (BIPP 0.97).** Clade 44 (Fig. 2C) was recovered with high support by Bayesian analyses whereas bootstrap values for both MP and ML were below 50%. The structure of the inflorescence is similar to the species of the *Capitatum* Clade (37) but here the partial cymes have less than 10 flowers and only one flower is fully developed in *T. paniculatum* (Clade 45), *T. euphrasioides* and *T. sertulariastrum* A.W.Hill (both Clade 46). The bracts subtending the aborted flowers are maintained on the floral axis forming an involucre below the flower, a feature that independently appeared in the tropical clade of *T. viride* A.W.Hill (Clade 61). All the species in this clade are distributed in the Fynbos Biome (mainly in the Western Cape Province with the exception of *T. polycephalum* Schltr., a characteristic species of the Namaqualand Hardeveld (Mucina & al., 2006). Several subclades were recovered with the highest support but some species were not included in any of them. *Thesium prostratum* A.W.Hill (Fig. 4O) and *T. selagineum*, two species of spreading shrublets with exserted anthers, share some features with species of the *Acuminatum* Clade (45, see below).

Unsampled species include *Thesium conostylum* Schltr., probably a synonym of the variable *T. hispidulum* Lam. The following may reside in subclades 45 or 47: *T. glomeratum* A.W.Hill, *T. glaucescens* A.W.Hill, *T. micromeria* A.DC., *T. rariflorum* Sond. (=*T. maximilliani* Schltr.), and *T. repandum* A.W.Hill. The remaining species are resolved in subclades 45 and 47.

**45. *Acuminatum* Clade (BIPP 1).** – The clade of *T. acuminatum* A.W.Hill (Fig. 2C) includes herbs or shrublets, all of them in *T.* sect. *Barbata*. With the exception of *T. paniculatum*, these plants have a dense woolly apical beard and acuminate corolla lobes. There are no evident morphological characters that define this clade because other species of Clade 44 not resolved here share similar features. Some species have characteristic long and subulate apices of the corolla lobes, but this character is also present in other species such as *T. hispidulum* or *T. prostratum*. The separation of species is based on the relative size of bracts and flowers or the presence of more or less developed leaves. In shrublets the latter character is probably variable with the age of the plant but those specimens that are clearly herbaceous have developed acicular and terete leaves (*T. acuminatum*, *T. paniculatum*) whereas in others, such as *T. capituliflorum* Sond. (Fig. 4P), the leaves are short and mostly apressed to the stems.

**46. *Hispidulum* Clade (BIPP 0.57). –** This clade (Fig. 2C) received very low support in the Bayesian analysis of ITS and was not recovered by MP and ML. The four accessions of *T. hispidulum* (Fig. 4Q) are monophyletic but this species shows great variation in vegetative features, from soft prostrate herbs to erect and stiff subshrubs. Variation exists in the stiffness of leaves and stems and the number of flowers of the terminal cymes. The unsampled *T. muasyae*, which was compared among other species with *T. hispidulum* (Zhigila & Muasya, 2022), is probably closely related to *T. sonderianum* (Clade 37). Sister to *T. hispidulum* a clade was recovered with the highest support, including *T. sertulariastrum* and *T. euphrasioides*, two species with a single flower fully developed in the inflorescence.

**47. *Commutatum* Clade (BIPP 0.69).** – This clade (Fig. 2C) received very low support when the accession 5238 identified as *T. densiflorum* A.DC. is included. The remaining accessions were recovered in a clade with high support (BIPP 0.97). *Thesium densiflorum* and *T. capitellatum* are very similar species distinguished by longer leaves and lanceolate bracts as long as flowers in *T. capitellatum* compared to shorter leaves and bracts shorter than flowers in *T. densiflorum*. These characters might be of low taxonomic value and further studies might show both taxa to be one variable species. Both taxa show corolla lobe tips, bracts and bracteoles that blacken with age. Because they share so many characters, identifying a particular individual as either of these species is frequently difficult. The three accessions of *T. commutatum* were monophyletic and resolved in a clade with *T. capitellatum* with the highest support. The flowers of *T. commutatum* (Fig. 4R) differ from that species in the papillose corolla lobe margins, lacking a dense apical beard. As mentioned above, Hill included it in *T.* sect. *Imberbia*; however, the vegetative characters are more similar to species in sect. *Barbata*, such as *T. capituliflorum* (Clade 45).

**48. Subgen. *Psilothesium* Clade (BIPP 1).** – Whereas the Core Cape Clade (32) represents most of the radiation of the genus in the CFR, the sister Clade 48 (Figs. 2C, 2D) represents the diversification of the genus in other Southern African and tropical African biomes, including independent dispersals to South America, Madagascar, and SE Asia. A few Cape species are also included here (see discussion below). This clade was also recovered with high support by chloroplast data (BIPP=1, MPBS=97, MLBS=98, Figs. S4--S7).

**49. *Gnidiaceum* Clade (BIPP 1).** – This group of South African species is sister to all remaining *Thesium* (Clade 50) that occur mainly in grassland and tropical areas (Fig. 2C). Most members of this clade were classified in the extremely heterogeneous *T.* sect. *Barbata*. It mostly includes species from the SE Great Escarpment and eastern Cape Fold Belt mountains: *T. impeditum* A.W.Hill (Fig. 4S, right image), *T. phyllostachyum* Sond. (Fig. 4T, right), *T. griseum* Sond., *T. gnidiaceum* A.DC. (Fig. 4S, left) and *T. durum* Hilliard & B.L.Burtt. The latter is a widespread species in eastern South Africa and Lesotho (Mashego & Le Roux, 2018). However, the two sampled species in *T.* sect. *Imberbia*, *T. oresigenum* Compton (Fig. 4T, left) and *Thesium* sp. 5574 are plants originating from the Cedarberg Mts and Matroosberg Mt in the Cape Fold Belt. The specimen 5574, from the Matroosberg, shows some morphological similarities with *T. congestum* R.A.Dyer (not sampled), but this latter species is endemic to the Drakensberg Alpine Centre (Carbutt & Edwards, 2006). Except for *T. durum*, species in *T.* sect. *Barbata* were recovered in a subclade with the highest support. They are characterized by the spicate inflorescences that, after fruiting, continue apical growth and elongate as a vegetative shoot. This feature is uncommon in *Thesium* (other examples are in Clade 52) since normally, when the apical meristem is terminated by forming an inflorescence, the vegetative growth continues from lateral branches at the base of the inflorescence.

Despite the strong support received for this clade, the topology of the chloroplast trees is different. *Thesium oresigenum* and *Thesium* 5574 are resolved as successively sisters to the remaining *Thesium* species of Clade 50, whereas the *T.* sect. *Barbata* species are highly supported as monophyletic but not as sister to the species in Clade 50. Both ITS and chloroplast sequences reveal the biogeographical importance of the Hex River and Cedarberg Mts in Western Cape for the two main radiations of the genus in Africa. The ITS phylogeny suggests a link between these ranges and the Drakensberg via the southern section of the Great Escarpment, a pattern that has been previously documented for other plant groups (Clark & al., 2011, 2012).

Unsampled species include *T. congestum*, *T. transvaalense* Schltr.?, and *T. confine* Sond. (= *T. spartioides* A.W.Hill). *Thesium confine* could belong to this clade because, together with *T. durum,* it was recognized as part of the *T. confine* complex by Mashego & Le Roux (2018) based on morphological features. *Thesium transvaalense* in tentatively included because it has some similarities with the *Barbata* species in this clade, such as the spike-like inflorescences. However, the inflorescence in this species in monotelic, ending in a flower with the vegetative branches developed from the base not from the tip of the inflorescence, and might be as well resolved together with other South African species in Clade 62. *Thesium rasum* (A.W.Hill) N.E.Br. was originally described as a variety of *T. impeditum* with the only difference being the short, sparse apical perianth hairs. Brown (1932) described several other differential characters and combined Hill’s variety to species. *Thesium rasum* has indeterminate inflorescences, apparently true spikes, and lacks the characteristic vegetative shoots at the tip as in *T. impeditum* and related species. Additionally, it is distributed in the Transvaal (not in the SE Great Escarpment) and might, as with *T. transvaalense*, be related to species in Clade 62.

**50. Tropical African Clade (BIPP 1).** – This clade (Figs. 2C and 2D) is mostly composed of the remaining species of the genus from Tropical Africa as well as grasslands and savannas in Southern Africa (excluding the Core Cape species). The well-supported monophyly of these species suggests a unique dispersal event from the Cape followed by diversification in the tropical parts of the continent and the rest of Southern Africa. The backbone relationships within this clade are poorly supported. Only the ITS Bayesian analyses resolve two major clades (51 and 62) but in both cases with low and very low support (0.76 and 0.53, respectively). Clade 51 is not resolved by the chloroplast genes whereas Clade 62 received a slightly higher BIPP value (0.66). All the species examined have short and straight placentas compared to the twisted placentas observed in the Core Cape species. The leaves are not decurrent and the stems are terete or ribbed by vascular strands ending in midribs of leaves and bracts. Exceptions are *T. kilimandscharicum* and *T. nigricans* Rendle, with sulcate stems and decurrent leaves typical of the Cape species (Hilliard, 2006).

**51. *Fimbriata* Clade (BIPP 0.76).** – This clade, which includes clades 52 through 61 (Figs. 2C and 2D), received low suppport with ITS and no support with chloroplast data. In general, their flowers lack apical beards on the corolla lobes, but more or less developed hairs or flaps are present on the margins. The degree of development of these structures varies greatly between and sometimes within species. Other characters shared are stamens with long filaments that are attached to the linear or lanceolate corolla lobes, variable length hypanthia, developed styles, and short and straight placentas. Many of these species have adaptations to occasional fires, such as well-developed woody rootstocks and stolons. In habit they are profusely branched and have buds densely covered with hard scales. Perennials predominate with annual species being less common. All of them have a tropical distribution, mostly in Africa, but also in Madagascar, South America and one in SE Asia (see below). Two clades with the highest support (52, 60) are resolved with ITS, but not with chloroplast data.

**52. LDD1 Clade (BIPP 1).** – The Long-Distance Dispersal (LDD) name for this clade (Fig. 2C) derives from the movement of propagules to South America and Madagascar from Tropical Africa. It has a good internal resolution with several subclades receiving strong support with BI, ML and MP. These clades were also seen with chloroplast data but with lower support. Two main subclades were recovered, the Usssanguense and Brachyblast Clades.

**53. *Ussanguense* Clade (BIPP 1).** – This clade (Fig. 2C) includes plants from the areas treated by Flora of Tropical East Africa, Flora Zambesiaca, and Flora of Central Africa. They are morphologically distinctive by the long corolla lobes with hairy margins (*T.* subsect. *Fimbriata*), flowers in elongated apparently true spikes (lacking both peduncle and pedicel), well-developed hypanthia, and long styles and staminal filaments. Many branches are born from a thick rootstock covered with numerous bracts from which basal flowering shoots arise. Based on these morphological features Polhill (2005) and Hilliard (2006) separated *T. ussanguense* Engl. in one of the six and nine informal groups in which they divided the genus for Flora of Tropical East Africa and Flora Zambesiaca, respectively. The other sampled Central African species resolved in this clade, *T. lewallei* Lawalrée and *T. passerinoides* Robyns & Lawalrée (Fig. 5A), are distinguished (among other characters) by the relative length of flowers and floral bracts. *Thesium ussanguense*, described from Tanzania, is distinguished from the Central African species by its broadly lanceolate basal leaves that are clearly membranous on the margins. Despite morphological differences that distinguish the species, ITS sequences showed the three sampled species emerging from a polytomy. Unsampled species include *T. malaissei* Lawalrée and *T. robynsii* Lawalrée.

**Fig. 5.**
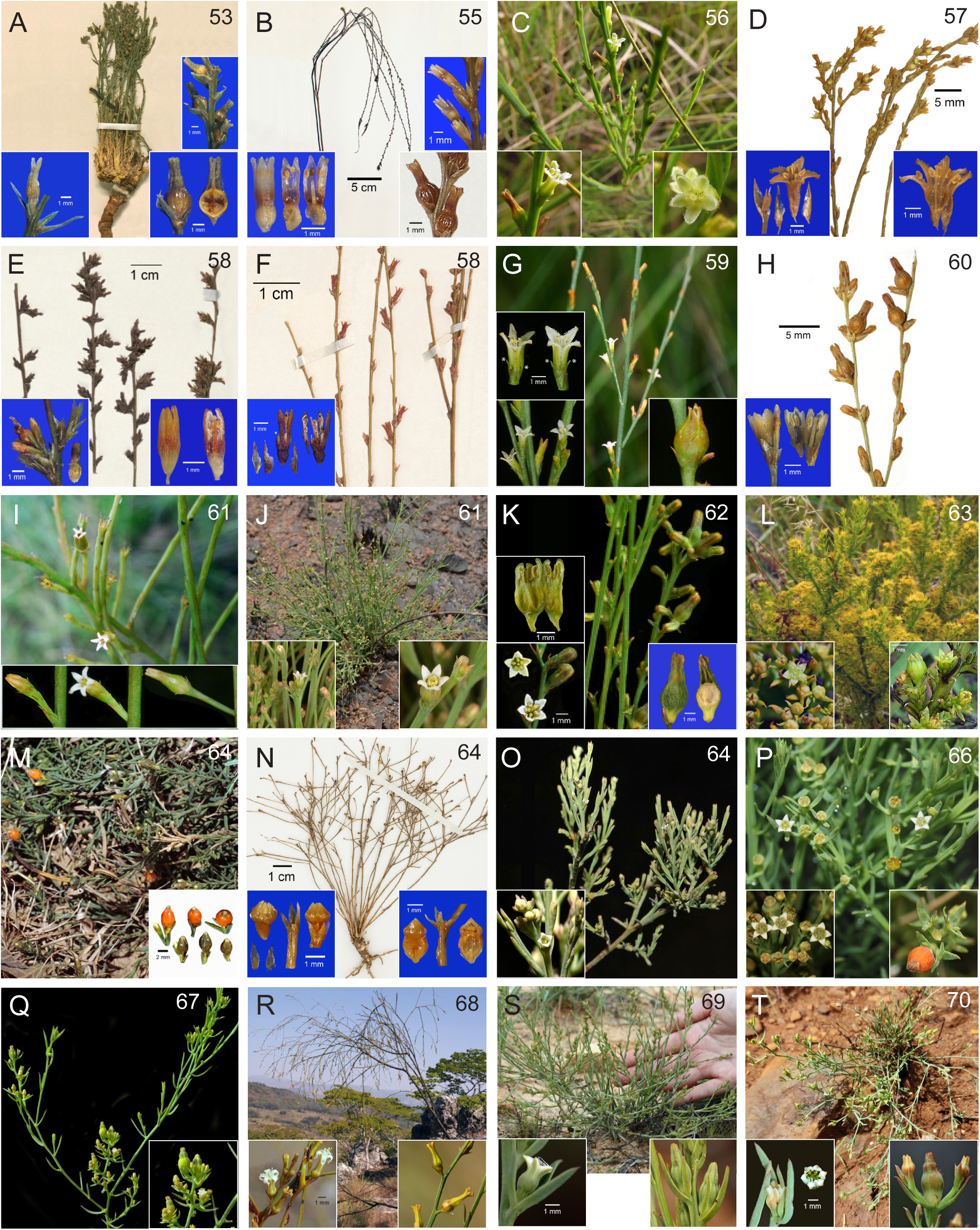
Photographs of representatives of the major clades of *Thesium* species. The numbers in the upper right corners correspond to clade numbers on the Bayesian trees (Fig. 2C-D). L.S = longitudinal section. **A,** *Ussanguense* clade, *T. passerinoides,* herbarium specimen (*Christiaensen 2495*, BR) with rehydrated flowers and fruits; **B,** *Stuhlmannii* clade, *T. fimbriatum,* herbarium specimen (*Goldblatt 8056*, MO) with rehydrated flowers and fruits; **C,** *Austroamericium* clade, *T. aphyllum*, habit and flowers; **D,** *Tenuissimum* clade, *T. tenuissimum*, herbarium specimen (*Meijer 15407*, BR) with rehydrated flowers; **E,** *Reekmansii* clade, *T. madagascariense*, herbarium specimen (*Evrard 11284*, BR) with rehydrated flower; **F,** *Reekmansii* clade, *T. wilczekianum* Lawalrée, herbarium specimen (*Milne-Redhead 3581*, BR), dissected flower from rehydrated herbarium specimen (*Malaisse 8407*, BR). Asterisk denotes calyx lobe; **G,** *Filipes* clade, *T. filipes*, flowering shoot, flowers, fruit. Asterisks denote calyx lobes; **H,** *Amicorum* clade, *T. amicorum*, herbarium specimen and rehydrated flowers (*Lisowski et al. 5736*, BR); **I,** *Viride* clade, *T. equisetoides*, flower shoots and flower developmental series; **J,** *Viride* clade, *T. fastigiatum*, habit and flowers; **K,** *Angulosum* clade, *T. angulosum*, shoot with floral buds, flowers, fruit L.S.; **L,** *Cupressoides* clade, *T. cupressoides*, habit, flowers, young fruits; **M,** LDD2 clade, *T. radicans*, habit, fruits; **N,** LDD2 clade, *T. psilotoides*, herbarium specimen and rehydrated flowers and fruits (*Williams 1310*, NY); **O,** LDD2 clade, *T. decaryanum,* flowering shoot and closer view of flowers; **P,** *Pallidum* clade, *T. pallidum*, flowering shoots, flowers, fruit; **Q,** *Cornigerum* clade, *T. cornigerum*, shoot with flowers and young fruits, close-up of same; **R,** *Kilimandscharicum* clade, *T. dolichomeras*, habit, flowers, fruits; **S,** *Resedoides* clade, *T. resedoides*, habit, flower, young fruits; **T,** *Gracile* clade, *T. gracile*, habit, flowers, fruit.

**54. Brachyblast Clade (BIPP 1).** – This clade (Fig. 2C) represents the first of two with members that have undergone long-distance dispersal, i.e. Tropical Africa to Madagascar and South America. The second LDD example (Clade 64) also involves Tropical Africa and Madagascar but also includes the enigmatic eastern Asian species *T. psilotoides*. Clade 54 includes species of Group 4 (excluding the annual species) of Flora of Tropical East Africa (Polhill, 2005) and Group 6 of Flora Zambesiaca (Hilliard, 2006), together with the three South American species and *T. madagascariense*. Most of the species in this clade inhabit temporary marshes and wetlands and few species (i.e. *T. stuhlmannii* Engl.) can grow in drier areas. ITS resolved three major subclades with the highest support that all arise from a polytomy.

In most species the stems are erect from narrow rootstocks and the leaves are reduced to scales. The flowers typically appear in short lateral shoots or brachyblasts with one to several flowers with a spike-like aspect (Fig. 5E). Bracts are frequently fimbriate and keeled, subulate or with an acute apex, and persist even when the flower is not developed, thus giving a scaly aspect to the lateral flowering shoots. The corolla lobes in most species are linear, sometimes fimbriate on the margins (*T.* subsect. *Fimbriata*), and the styles and filaments are long. In some species projections that could be interpreted as traces of a calyx alternate with the corolla lobes, thus challenging the concept that these are monochlamydous flowers. In some species such as *T. libericum* or *T. wilczekianum* Lawalrée these projections are quite developed (up to 1 mm long, Fig. 5G, asterisks in the upper left inset). In other species they are reduced to swollen enlargements of the corolla tube between the lobes. These structures are unique to this clade but might be homologous to the so-called perianth glands which are well developed in many African species. The taxonomy of this group is difficult because the structure of the inflorescences, including the development of the brachyblasts, may change with age, as discussed by Hilliard (2006: 223).

Unsampled species include *T. helodes* Hilliard, *T. inonoense* Hilliard, *T. leucanthum* Gilg, *T. libericum*, *T. masukense* A.W.Hill, *T. matteii* Chiov., and *T. perrieri* Cavaco & Keraudren.

**55. *Stuhlmannii* Clade (BIPP 1).** – This clade (Fig. 2C) includes two morphologically close species, *T. fimbriatum* (Fig. 5B) and *T. stuhlmannii*. The latter has glabrous margins on the corolla lobes and smaller fruits than the former (Hilliard, 2006) and is distributed in the area of Flora of Tropical East Africa and Ethiopia (Polhill, 2005). *Thesium fimbriatum* has a more southern distribution in the area of Flora Zambesiaca and D.R. Congo.

Our accession 4819 is a creeping plant from Ethiopia that Miller (1989) identified as *T. matteii* in his treatment for Flora of Ethiopia and Eritrea. This plant, however, is clearly different from the holotype of that species, which has well developed and elongated brachyblasts and flowers and bracts densely grouped at the tips, whereas 4819 has mostly sessile, solitary flowers in elongated spike-like inflorescences. Miller stated that he didn’t see the type of *T. matteii* and this plant might be a new species.

Unsampled species include *T. shabense* Lawlarée, which was considered a synonym of *T. fimbriatum* by Polhill (2005) and Hilliard (2006).

**56. *Austroamericium* Clade (BIPP 1).** – This clade (Fig. 2C) suggests a unique long-distance dispersal episode from Africa followed by speciation in two separated areas, the savannas of the Guiana Shield in Venezuela and Guyana (*T. tepuiense* Steyerm.) and the highlands of C, E and S Brazil (*T. aphyllum* Mart. ex A.DC., and *T. brasiliense* A.DC.). These species were segregated by Hendrych (1963b) in the genus *Austroamericium*, based on characters that are either nonexistent or also present in the African species of Clade 53. Hendrych considered that the cucullate corolla lobes, the stamens with long filaments, the shape of the fruit, the scale-like leaves and the annual life cycle of the South American species were sufficiently unique to warrant generic segregation. However, all those characters are also present in many Old World species of *Thesium*. The annual life cycle, although not common among the tropical species, does exist as demonstrated by some members of Clade 60 (see below). Moreover, *T. aphyllum* (Fig. 5C) is a perennial with stems from a thick rootstock or a rhizome and therefore the statement that all the South American species are annual is incorrect. Hendrych indicated that the clearest character to recognize *Austroamericium* is the dehiscence of the lobes of the perianth in fruit, a character that is frequently observed in herbarium specimens of *T. aphyllum* but not in *T. brasiliense* and *T. tepuiense*. These species share many other morphological characters with African species of the Brachyblast Clade, including the short and straight placenta.

**57. *Tenuissimum* Clade (BIPP 1).** – This clade (Fig. 2C) includes most of the sampled species with brachyblasts and consists of a polytomy of *T. tenuissimum* Hook f., *Thesium* sp. 4825 and Clades 58 and 59. *Thesium tenuissimum* (Fig. 5D) has its area of distribution in tropical West Africa and is characterized by numerous profusely branched thin stems that arise from a woody rootstock. *Thesium* sp. 4825, from Ethiopia, was identified by Miller (1989) as *T. stuhlmannii* but it differs from this species in several features and is more similar to the holotype of *T. matteii*. If this specimen is shown to be *T. mattei*, the Madagascan *T. perrieri* might be part of Clade 57 given their morphological similarity.

**58. *Reekmansii* Clade (BIPP 1).** This clade (Fig. 2C) contains three species named by Lawalrée (1985) from Tropical Africa as well as *Thesium madagascariense* from Madagascar (Fig. 5E). *Thesium wilczekianum* is characterized by its conspicuous (approaching 1 mm long) tooth-like protruberances (calyx lobes) in the sinuses of the corolla lobes (Fig. 5F). *Thesium schaijesii* Lawalrée that has shorter protruberances, flowers in long spike-like inflorescences composed mostly of single flowers but many with additional bracts at the base. According to Hilliard (2006), *T. schaijesii* is a synonym of *T. schliebenii* Pilg. but this species is resolved here in Clade 59. *Thesium reekmansii* Lawalrée differs from *T. schliebenii* by its mostly 4-merous flowers and smaller fruits with an inconspicuous network of nerves on the pericarp. Morphologically, *Thesium madagascariense* is clearly a member of the Brachyblast Clade, but its closest relatives among the African species are different according to ITS or chloroplast sequences. For the latter it is resolved with the highest support (BIPP=1) with *T. fimbriatum* and *T. stuhlmanii* (Clade 55). The ITS topology suggests that the dispersal from the continent to Madagascar occurred several times independently. Other Madagascan species sampled for this study come out in different clades: *T. decaryanum* in Clade 64 and *T. pseudocystoseiroides* in Clade 74. Morphological features of these species also support at least three independent dispersals (see below).

**59. *Filipes* Clade (BIPP 0.91).** Clade 59 (Fig. 2C) includes *T. vimineum* Robyns & Lawalrée from Central Africa with long spikes lacking brachyblasts, sessile pentamerous flowers, and fruits with up to 20 longitudinal ribs. *Thesium aellenianum* Lawalrée has the persistent perianth shorter than the not reticulate fruit. *Thesium filipes* A.W.Hill (Fig. 5G), distributed in tropical regions of Central and West Africa, has mostly tetramerous flowers in elongate spike-like inflorescences with short brachyblasts and developed tooth-like protruberances, here interpreted as calyx lobes. The unsampled *T. libericum* has been considered as a synonym of this species.

**60. *Amicorum* Clade (BIPP 1).** – Clade 60 (Fig. 2D) includes some of the few annual tropical species of the genus: *T. subaphyllum* Engl., *T. symoensii* Lawalrée and *T. amicorum* Lawalrée (Fig. 5H). ITS resolved the two latter species in a very poorly supported clade with *T. microphyllum* Robyns & Lawalrée and the three of them sister to the *Viride* Clade (Clade 61). *Thesium microphyllum* is not annual but has long and thin rhizomes, not a thickened rootstock. The annual species have a pair of subopposite developed leaves at the base (probably the persistent cotyledons) with the others being scale-like. Inflorescences are elongated spikes, with solitary flowers or with one or two additional ones. Bracts and bracteoles have the typical naviculate shape of species in the *Fimbriata* Clade (51). Corolla lobes margins are not fimbriate but have inflexed, membranous, erose flaps, more similar to the species in the *Viride* Clade (61). Fruits mostly have reticulate nerves between the ribs whereas in most of the species in the *T. fimbriatum* clade the fruits are not reticulate or reticulations are few and often obscure.

Unsampled species include *T. annuum* Lawalrée, *T. brachyanthum* A.W.Hill and *T. lisowskii* Lawalrée as well as *T*. *singulare* Hilliard that might belong here or in the *Viride* Clade (61). *Thesium rectangulum* Welw. ex Hiern, described from Angola, is an annual herb (Baker & Hill, 1911) with developed leaves (not reduced to scales) and flowers axillary or located at the terminus of lateral leafy branches (as in the species of the *Viride* Clade).

**61. *Viride* Clade (BIPP 0.99).** This clade 61 (Fig. 2D) includes a group of at least 15 species that are all closely related and that emerge from a polytomy. This subclade is sister to *Thesium subaphyllum* (=*T. andongense* Hiern), an annual species distributed from Angola to tropical East Africa. Its sister relationship with the remaining members of the *Viride* Clade is resolved with ITS but not with chloroplast markers. Morphologically, it shares features with the annual species of Clade 60 such as the presence of a pair of opposite leaves up to 3 cm long or leaf scars at the base of the flattened and somewhat alate stem. Inflorescences are spikes but sometimes lateral branches with few flowers and several bracts are present. The bracts and bracteoles are small and are reminiscent of those of species in the *Viride* Clade. The fruits are rounded with a very reticulate surface.

Sister to *T. subaphyllum* is a clade of exclusively tropical African perennial species that was included in group 7 in Flora Zambesiaca (Hilliard, 2006) and group 5 in Flora of Tropical East Africa (Polhill, 2005). Although ITS sequence analyses result in a polytomy, higher internal resolution in this clade is obtained with combined chloroplast markers (Figs. S4--S7). Members of this clade are readily characterized by shoots that arise from a more or less compact and branched woody rootstock bearing subulate and frequently spinous leaves as well as by the presence of an involucre of 4–6 bracts at the base of the flowers.

Vegetatively they are similar to species in the *Ussanguense* Clade (53) and both groups are pyrophytes. Some species are metallophytes, tolerant of high concentrations of copper and cobalt in the soil. The perianth lobes are not bearded and have entire margins or margins inflexed and more or less erose, especially at the base. The flowers may be terminal, but frequently a lateral branch may develop from the base of one of the bracts subtending the flower. The characters used for species delimitation include the presence or absence of hairs or cilia on the whole plant, size and shape of the floral bracts, presence or absence of developed leaves at least at the base of the stems, length and number of flowers on the flowering branches, shape of leaves, angle of lateral branches, presence or not of bract-like leaves on the peduncles subtending the terminal flower, diameter of the branches and color of the plant when dry. Infraspecific variation related to the age of the plant may confound species identification.

Clade 61 includes 15 species with identical or nearly identical ITS sequences: *T. doloense* Pilg., *T. equisetoides* Welw. ex Hiern (Fig. 5I), *T. fastigiatum* A.W.Hill (Fig. 5J), *T. germainii* Robyns & Lawalrée, *T. hockii* Robyns & Lawalrée, *T. lopollense* Hiern, *T. luembense* Robyns & Lawalrée, *T. myriocladum* Baker ex A.W.Hill, *T. pawlowskianum* Lawalrée, *T. setulosum* Robyns & Lawalrée, *T. subaphyllum*, *T. tamariscinum* A.W.Hill, *T. thamnus* Robyns & Lawalrée, *T. unyikense* Engl., and *T. viride* A.W.Hill. The lack of internal resolution of this clade resembles the section *Frisea* Clade (39), but these taxa are not so diverse morphologically, thus some of the species described might be synonyms.

Unsampled species includes *T. atrum* A.W.Hill, *T. bangweolense* R.E.Fr., *T. bequaertii* Robyns & Lawalrée, *T. cinereum* A.W.Hill, *T. crassipes* Robyns & Lawalrée, *T. fanshawei* Hilliard, *T. fulvum* A.W.Hill, *T. fuscum* A.W.Hill, *T. katangense* Robyns & Lawalrée*, T. laetum* Robyns & Lawalrée, *T. lycopodioides* Gilg, *T. lynesii* Robyns & Lawalrée, *T. magnifructum* Hilliard, *T. manikense* Robyns & Lawalrée, *T. nutans* Robyns & Lawalrée, *T. pilosum* A.W.Hill, *T. quarrei* Robyns & Lawalrée, *T. strigulosum* Welw. ex Hiern (nom. illeg.), *T. tetragonum* A.W.Hill, *T. triste* A.W.Hill, and *T. wittei* De Wild.

**62. Grassland Clade (BIPP 0.53).** – Clade 62 (Fig. 2D) is recovered in ITS and combined chloroplast Bayesian analyses with very low support (0.53 and 0.63 BIPP, respectively). Although including also species of savannas and other biomes, most species occur in grasslands. Some are also present in Tropical Africa, S Arabian Peninsula, Madagascar, and E and SE Asia. Except for *T. angulosum* A.DC. (Fig. 5K), these species are resolved with ITS in three major clades, two of them (63 and 64) with the highest support and Clade 65 with very low support.

**63. *Cupressoides* Clade (BIPP 1).** – Clade 63 (Fig. 2D) consists of prostrate herbs to erect shrublets, all from *Thesium* sect. *Imberbia*, that are distributed in the Drakensberg, the Sneeuberg and neighboring regions of Lesotho and South Africa (Eastern Cape, KwaZulu-Natal and SE Free State Provinces). These are densely leafy plants, with leaves frequently recurved and more or less scabrid. All the species in this clade have twisted placentae. It is a taxonomically difficult group, especially the complex of *T. acutissimum* A.DC., which is morphologically variable in the degree of inflorescence branching. It is often difficult to separate from the closely related *T. disciflorum* A.W.Hill and *T. squarrosum* L.f. (not sampled), both of which have similar flower morphology and mucronate to subulate leaves. Additionally, these taxa showed polymorphic ITS and the two sequences of *T. disciflorum* obtained by TA cloning were succesively sister to the clones of *T. acutissimum*. *Thesium cupressoides* A.W.Hill (Fig. 5L) is an erect shrublet frequent seen in the Drakensberg and lower areas of KwaZulu-Natal. It is sister to the remaining species in this clade with ITS but not with chloroplast data. Its shrubby habit is useful to separate it from *T. acutissimum* but when flowers are crowded in dense inflorescences (*T. acutissimum* var. *corniculatum* A.W.Hill) these two species are not easily differentiated on herbarium specimens.

The plant with accession number 4818 is one of several Drakensberg collections of creeping plants with thin rhizomes. They are sometimes misidentified on herbarium labels as *T. racemosum* Berh. ex Krauss but likely represent a new species. *Thesium racemosum* is known from lower areas of Eswatini and E South Africa, differing in several morphological features, such as the erect stems and a different inflorescence. The flowers of accession 4818 are cup-shaped on lateral cymules of 1--5 flowers, with a short corolla tube, short style and conspicuous external glands. Morphologically it is distinctive among the species in this clade but its phylogenetic position is supported by both ITS and chloroplast sequences.

In addition to *T. squarrosum*, unsampled species may include *T. cystoseiroides* Baker, from Madagascar, which is a profusely branched and prostrate shrublet that might be related to species in this clade given its habit, mucronate leaves and morphology of flowers and fruits that are similar to *T. acutissimum*. *Thesium leandrianum* Cavaco & Keraudren, also from Madagascar, has similar floral features, with bracts, bracteoles and leaves strongly recurved, as shown on the type specimens.

**Clades 64**--**65.** – The combined chloroplast analyses resolved a major clade containing all the species of clades 64 and 65 with the highest support (Figs. S4, S5), suggesting that excluding the species in Clade 63 these species are monophyletic. This topology was also obtained in a more reduced sampling by Moore & al. (2010), in which a clade formed by *T. radicans* and *T. schweinfurthii* Engl. (our Clade 64) were sister to a clade containing six species, four of which are in our Clade 65. Unfortunately, we could not obtain chloroplast sequences from the unplaced *T. angulosum*, which was resolved in neither clade with ITS but is in a polytomy with Clades 63--65. This shurblet in *T.* sect. *Imberbia* from E KwaZulu-Natal is distinctive given the combination of erect and winged stems that branch above, elongated spike-like inflorescences with lateral cymules composed of 1--3 flowers, a hypanthium half the length of the flower, and corolla lobes with papillae or fimbriae on the margins (Fig. 5K). An unsampled species, *T. decipiens* Hilliard & B.L.Burtt, also from Natal but found at higher elevations, is morphologically very similar to *T. angulosum* and might be closely related, but the corolla lobes in that species are bearded (Hilliard & Burtt, 1983) and could be misidentified as *T. alatum* Hilliard & B.L.Burtt (Clade 72).

**64. LDD2 Clade (BIPP 1).** – This clade was recovered with the highest support (Fig. 2D) including species from Tropical Africa and S Arabian Peninsula (*T. radicans*, *T. schweinfurthii*, *T. schmitzii* Robyns & Lawalrée), East Asia (*T. psilotoides*), and Madagascar (*T. decaryanum*). The clade was named based on the long-distance dispersal that characterizes the last two members. *Thesium decaryanum* (Fig. 5O) is sister to the entire clade, *T. radicans* is sister to the Asian *T. psilotoides* and that clade sister to *T. schweinfurthii* and *T. schmitzii*. Chloroplast data also placed members of this clade together with the highest support. However, it also included species resolved with ITS in Clade 69 (see below): *T. resedoides* A.W.Hill, *T. mukense* A.W.Hill, *T. burkei* A.W.Hill and *T. hararensis* A.G.Mill. In the chloroplast phylogenies *T. decaryanum* is strongly supported as sister to all of them. *Thesium schmitzii* is generally considered a synonym of *T. schweinfurthii* and our results support this.

Despite being resolved in Clade 64 with the highest support, these species exhibit quite different vegetative and floral morphologies. For example, *Thesium schweinfurthii* and *T. schmitzii*, have an apical beard of hairs at the apex of the corolla lobes. In *T. decaryanum* and *T. radicans* the beard is present but very reduced and in *T. psilotoides* it is completely lacking. *Thesium radicans* is one of the few species in the genus with fleshy orange-red fruits (Fig. 5M), a syndrome that has arisen at least two other times independently in the evolution of the genus (see discussion of the *Triflorum* Clade). It is also distinctive in having thin, creeping rhizomes which produce short, mat-forming stems. Both ITS and chloroplast phylogenies resolved *T. psilotoides* (Fig. 5N) in this clade, a species described from S China and later collected in SE Asia and the Philippines, where it is the only native member of the genus in these areas. Hendrych (1972) considered it to be related to the Eurasian species despite possessing unusual characters for that group (e.g., scale leaves). Our results indicate that *T. psilotoides* originated following a long-distance dispersal event. In addition to being distinct from Eurasian species morphologically, the fruit of *T. psilotoides* is more similar to species in the *Viride* Clade (Fig. 5N). The presence of a short and straight placenta, instead of the apically twisted placentae of the Eurasian species, is maybe the morphological feature that more clearly shows its relationship with tropical African taxa.

Unsampled species includes *Thesium wightianum* (= *T. nilagiricum* Miq.), from SW India, which apparently has ciliate corolla lobes and a short and erect placenta, characters that approach tropical African rather than Eurasian taxa. Morphologically it is reminiscent of *T. radicans* but the fruits apparently are not fleshy. Its relationship with *T. radicans* was first suggested by Candolle (1857) who considered *T. radicans* a variety of *T. wightianum*; DNA sequence information of this species would conform a biogeographical connection with species from tropical East Africa, Arabian Peninsula, Madagascar and also with *T. psilotoides*.

**65. *Pallidum*-*Costatum* Clade (BIPP 0.65).** – This clade (Fig. 2D), which contains the remainder of *Thesium* species analyzed here, received very low Bayesian support with ITS sequences but received the strongest support in chloroplast analyses (BIPP = 1). It lacked internal resolution with chloroplast data whereas a number of well-supported clades were recovered with ITS. Clades 66--71 all arise from a polytomy as do subclades within Clade 73. The geographical distribution of its component species is broad including the savanna and grassland biomes of South Africa, Eswatini, the area covered in Flora Zambesiaca, Tropical Africa and Madagascar. The species of Hilliard’s (2006) groups 2, 3 and 8 (except *T. subaphyllum*) and Polhill’s (2005) groups 2 and 6 (except *T. schweinfurthii*) are resolved here. With a few exceptions, all these species have more or less developed leaves and leafy floral bracts that are not scale-like. Inflorescences are diverse as shown by Visser & al. (2018) for the *T. goetzeanum* Engl. complex. Frequently they are branched with cymose or racemose patterns, often with lateral cymes bearing a variable number of flowers. Some species exhibit spike-like inflorescences probably as the result of the reduction of number of flowers since some specimens may show short lateral cymules with two or three flowers. Another important character is the degree of metatopic displacement of the floral bracts along the peduncle (recaulescence). This character appears to be variable within moderately or well-supported clades and may include species with different states of this character. However, it is constant in well-developed inflorescences of the same species, thus having taxonomic value. The species derive from both sections *Imberbia* and *Barbata* and thus show variation in the ornamentation of the corolla lobes. Most of the species resolved in this clade have corolla lobes bearded and most of the subclades are exclusively formed by species in *T.* sect. *Barbata*. Species of both sections may resemble each other vegetatively, independent of this floral character.

All the species of the *T. goetzeanum* complex as defined by Visser & al. (2018) that were sampled for this study were resolved in different subclades clades within Clade 65. That taxonomic revision of the complex was based on plants from South Africa, Lesotho, and Eswatini and many species names were reduced to synonyms. However, species with similar morphological characters are not exclusive to those countries and are also present in tropical Africa and Madagascar. To investigate the monophyly of the species complex as were defined by Visser & al. (2018) and in older revisions, we have tried to identify the plants used in this work following other taxonomic treatments and at the same time compare these plants with the type material of species accepted by Baker & Hill (1911), Hill (1915a, 1915b), Brown (1932), Thulin (1999), Polhill (2005) and Hilliard (2006).

Unsampled species with bearded corolla lobes that probably would be resolved in Clade 65 are as follows:

- *Thesium celatum* N.E.Br. is a species that was only known from the type specimen collected at the Waterberg in Limpopo Province (South Africa). Lombard & Le Roux (2023) studied more recent collections and provided a detailed description of the species comparing it morphologically with *T. burchellii* A.W.Hill (see discussion of Clade 70). Instead of straight placental column it is “slightly to prominently undulate but not twisted” in this species and, as for *T. procerum* N.E.Br. (see below), molecular data is neccesary to establish its phylogenetic relationships.
- *Thesium dinteri* A.W.Hill, from Namibia, has been treated as a synonym of *T. megalocarpum* A.W.Hill (Roessler, 1969). However, Hilliard (2006) recognized them as two different species and Brown (1932), when describing *T. dumale* N.E.Br., concluded it was very close to the type of *T. dinteri* but with smaller flowers. Brown’s *T. dumale* has been reduced to a synonym of *T. resedoides* by several authors.
- *Thesium hillianum*, a small shrub from Whitehill near Lainsburg (Witterberg) in the Western Cape Province, has indeterminate inflorescences with axillary solitary flowers or in 3-flowered cymules and the bract free or only partially fused to the peduncle. The flowers have showy external glands and the corolla lobes have long hairs along the whole margin rather than an apical beard. More evidence that this species might be resolved in Clade 65 are that its stems are terete and non-sulcate, as in most species in the clade. We could not study the placentation in the type specimens but this character might help to clarify its relationship. Some collections from the Namaqualand misidentified as *T. hillianum* have clearly sulcate stems and differ in several other characters. The Namaqualand plants are probably *T. urceolatum* and would thus be placed in Clade 39 (*T.* sect. *Frisea*). Compton (1931) included his species in *T.* sect. *Penicillata* because the anthers are not attached to the post-staminal corolla hairs; however, *T. penicillatum* differs in many vegetative and reproductive characters (see discussion of Clade 36).
- *Thesium hispidifructum* N.Visser & M.M.leRoux is a recently described subshrub or shrub endemic to the Limpopo Province in South Africa (Lombard & Le Roux, 2023). It was compared in the diagnosis with *T. disparile* N.E.Br. (Clade 80), a minutely scabrid herbaceous species only known from the type collection and the specimen sampled for this study, collected in the Rustenburg District of the North West Province. Although also compared with *T. rufescens* (Clade 32), this new species would probably be resolved in Clade 65 for its straight placental column.
- *Thesium infundibulare* N.Visser & M.M.leRoux, from KwaZulu-Natal and Eswatini is a species in the *T. goetzeanum* species complex characterized by its stipitate fruits. Visser & al. (2018) compare their new species with *T. resedoides*. Other species in Clade 65 have stipitate fruits including *T. vahrmeijeri* Brenan (Clade 67), *T. procerum*, and *T. mossii* N.E.Br. (see below).
- *Thesium junodii* A.W.Hill, from the Transvaal, was described based on its slender habit with recurved leaves and prominent external glands. Brown (1932) suggested a relationship with *T. breyeri* N.E.Br. whereas Visser & al. (2018) included the species as a synonym of *T. resedoides*. The lectotype of *T. junodii* (*Junod 1301*, K!) has flowers with a tube ca. 1.5 mm long and style 1.5--1.75 mm long, above the range given by Visser & al. (2018) for *T. resedoides*. The overall vegetative (erect stems with raised vascular bundles) and floral (long hypanthium and style) morphology suggests its relationship with species of *T.* sect. *Imberbia* in Clade 75.
- *Thesium mossii* was included in the synonymy of *T. resedoides* by Visser & al. (2018), but the type of *T. mossii* has clearly stipitate flowers and fruits, a character considered of taxonomic importance by the authors, whereas *T. resedoides*, according to their description, has sessile flowers and fruits.
- *Thesium ovatifolium* N.Lombard & M.M.leRoux, from the grasslands of KwaZulu- Natal, has broadly ovate leaves and bracts with reticulate secondary venation, a character uncommon in the genus. Lombard & al. (2019) compared this species in the diagnosis with *T. goetzeanum* and *T. racemosum*.
- *Thesium pottiae* N.E.Br., described from the former Transvaal (northern provinces of South Africa), is among the species with elongated spike-like inflorescences similar to e.g.,
- *T. magalismontanum* Sond. (Clade 76), *T. asterias* A.W.Hill (Clade 80) or *T. polygaloides* A.W.Hill (Clade 79). The floral bract is either partly or fully adnate to the short (1--2 mm) peduncle. Different style lengths in the species suggested heterostyly to Brown (1932). It resembles *T. cornigerum* A.W.Hill from KwaZulu-Natal (Clade 67) but in this species flowers are mostly in lateral cymules rather than solitary. It is probably a synonym of *T. multiramulosum* Pilg. (Clade 74) as suggested by Hilliard (2006).
- *Thesium procerum* was included in the *T. goetzeanum* species complex (Visser & al., 2018) as the only species in the complex with a twisted placenta. This type of placentation is uncommon within species of Clade 65 and further molecular data would establish the relationship with other species of the clade.
- *Thesium rogersii* A.W.Hill was considered as a synonym of *T. goetzeanum* by Hilliard (2006) and Visser & al. (2018). The type specimen is a glaucous plant with bracts shorter than the flowers and angulate stems but the rest of morphological characters match with *T. goetzeanum*.

**66. *Pallidum* Clade (BIPP 1).** – This clade (Fig. 2D) includes two collections of *T. pallidum* A.DC., a grassland species of South Africa (Fig. 5P). Features include flowers borne in cymose branched inflorescences, lateral cymes with floral bracts fused up to half or less to the peduncle, well-developed floral disks, and very short hypanthia and styles. These plants are commonly tinted orange when rubbed or cut and the fruits can be of that color when mature. This species can be easily mistaken for *T. acutissimum* (Clade 63) because of similar inflorescences and floral characters; however, our molecular data indicate that they are not closely related. One character that might differentiate the two clades is that in Clade 63 species the placenta is twisted, whereas in the specimens of *T. pallidum* sampled for this study the placenta is straight. Further morphological and molecular studies including more samples of this widespread species, are necessary to find additional morphological characters to separate them and to confirm whether *T. floribundum* A.W.Hill is a synonym of *T. pallidum* as suggested by Polhill (2005).

The unsampled *Thesium asperifolium* A.W.Hill is a scabrid species that share many morphological features with *T. pallidum* and could be closely related.

**67. *Cornigerum* Clade (BIPP 0.99).** – This well-supported clade (Fig. 2D) includes the annual *T. vahrmeijeri*, a species included in the *T. goetzeanum* complex by Visser & al. (2018), which is sister to the clade of *T. cornigerum* and *Thesium* sp. 4686. Vegetatively *T. vahrmeijeri* and *T. cornigerum* are clearly different, the former an annual that is profusely branched from the base whereas the latter generally has erect, virgately branching stems. Also *T. cornigerum* has long inflorescences, frequently with lateral cymules containing three or more flowers, and the bracts are free or only adnate at the base of the peduncle (Fig. 5Q). Accession 4686, from the former Transvaal, is clearly different morphologically from both *T. vahrmeijeri* and *T. cornigerum*, having elongated spike-like inflorescences with apparently indeterminate growth, very shortly pedicellate flowers and corolla lobes with apical dendritic hairs. Although the key published by Brown (1932) leads to *T. pottiae*, not all the characters described match with this plant.

**68. *Kilimandscharicum* Clade (BIPP 1).** – This clade (Fig. 2D) contains two species, *T. kilimandscharicum* and *T. dolichomeras* Brenan. The sister relationship of these two species is surprising given their strikingly different vegetative and floral morphologies. *Thesium kilimandscharicum* is a prostrate to suberect annual or perennial with ribbed stems that are densely covered with decurrent leaves. Its flowers are small, mostly tetramerous with a very short style. *Thesium dolichomeras*, however, is an erect tall shrublet with sparse leaves, and large pentamerous flowers on long spike-like inflorescences, very long styles and corolla lobes with long hairs on the edges but lacking an apical beard (Fig. 5R). An additional specimen of *T. dolichomeras* was sequenced and its ITS sequence was identical to accession 4695 and therefore, a sequencing error can be ruled out. Conversely, chloroplast phylogenies do not resolve *T. dolichomeras* as sister to *T. kilimandscharicum* but to a group that includes most of the leafy tropical species. Additional phylogenetic data including species morphologically similar to both *T. kilimandscharicum* (e.g. *T. nigricans*, *T. pygmaeum* Hilliard) and *T. dolichomeras* (e.g. *T. chimanimaniense* Brenan) are needed to confirm their phylogenetic relationship.

Unsampled species include *T. nigricans* (=*T. scabridulum* A.W.Hill), *T. pygmaeum*, and *T. chimanimaniense*.

**69. *Resedoides* Clade (BIPP 0.99).** Both ITS (Fig. 2D) and chloroplast data strongly support a clade composed of *T. resedoides*, *T. mukense*, and *T. hararensis*. *Thesium resedoides* is a species with a broad geographical distribution from Angola to southern

Mozambique, Eswatini, and NE KwaZulu-Natal in South Africa. The other two species occur north of that distribution and east of the Great Rift Valley from Mozambique to Kenya (*T. mukense*) and Ethiopia and Somalia (*T. hararensis*). All are much branched shrublets with generally short racemose inflorescences at the tips of the branches. The morphology of *T. resedoides* in South Africa was studied in detail by Visser & al. (2018) who described the species as a rhizomatous suffrutex. Our accession number 4754, one of the specimens listed by these authors as *T. resedoides*, has the typical vegetative and floral features of the species except for the stems developed from an apparently non-rhizomatous rootstock. ITS did not place this accession in clade 69 (with accessions 4424 and 4823) but with low support in Clade 70 whereas the chloroplast analyses included it in the *Resedoides* Clade with strong support.

*Thesium mukense* is morphologically similar to *T. resedoides* in vegetative and floral characters and both have woody rhizomes. In the SW of the Flora Zambesiaca area, these two species can be distinguished by the shorter, broader, and strongly recurved bracts of *T. resedoides* (Hilliard, 2006). Bract lengths in *T. resedoides* are variable and further studies might show that *T. mukense* are northern populations of a widespread *T. resedoides*. Visser & al. (2018) did not include *T. mukense* among the synonyms of *T. resedoides* but the morphological features of both species overlap. *Thesium mukense* has also been placed in synonymy with *T. whyteanum* Rendle, however, our ITS data place that taxon in Clade 73.

*Thesium hararensis* shares morphological features with *T. resedoides* and *T. mukense* but the bracts are much shorter than the flowers and fruits and the plants appear to lack woody rhizomes, at least in the herbarium samples studied. In addition, the lower leaves in young specimens are terete and subsucculent such as in the individual sampled here (accession number 4814).

The combined chloroplast trees (Figs. S4--S7) show a well-supported sister relationship between the *Resedoides* Clade (69) and the LDD2 Clade (64), with *T. decaryanum* as sister to both clades. Possibly an ancient hybridization event in the common ancestor of the species from both clades might explain this discordance between data sets. Further discussion on the morphological similarities between species of the *T. resedoides* group with *T. schweinfurthii* is provided by Polhill (2005) and Hilliard (2006).

Unsampled species include *T. angolense* Pilg. (from Angola) which might belong in this clade. It has some features similar to *T. resedoides*, such as the adnate, recurved and subulate bracts but is also similar to other species in the *T. goetzeanum* complex. It was not included in the study by Visser & al. (2018).

**70. *Gracile* Clade (BIPP = 0.68).** This clade (Fig. 2D) is composed of *T. deceptum* N.E.Br., *T. gracile* A.W.Hill (Fig. 5T), *T. panganense* Polhill and one accession of *T. resedoides* (4754). If this latter accession is excluded the three first species were resolved in a clade with the highest support. *Thesium panganense*, endemic to the Pangani District in Tanzania, has well developed dichasial cymes and minutely ciliolate corolla lobes. According to Polhill (2005) this species is morphologically similar to *T. pallidum* (=*T. floribundum*, see Clade 66) but they differ in the shape of hypanthium tube and style length. In our phylogenies these species are not resolved as closely related. The other two species, *T. gracile* (=*T. palliolatum* A.W.Hill) and *T. deceptum*, both in *T.* sect. *Barbata*, were included by Visser & al. (2018) in the *T. goetzeanum* species complex (*T. deceptum* as a synonym of *T. goetzeanum*). This relationship was not resolved by the chloroplast dataset. *Thesium panganense* and *T. gracile* have similar cymose inflorescences, with the bracts free or adnate only at the base. The sampled *T. gracile* showed polymorphic ITS but both sequences were resolved in the same clade. *Thesium deceptum* is distinguished by the markedly scabrid and cartilaginous margins of leaves, bracts and bracteoles. Its inflorescence is less branched than in the other species of this clade and its flowers have prominent perianth glands. According to Hilliard (2006) *T. gracile* is closely related to *T. brevibarbatum* Pilg. (Clade 80) but its flowering branches appear at the base of the plant and the flowers are mostly single in the axils of the bracts.

Unsampled species include *T. inhambanense* Hilliard from southern Mozambique, which has similar branched cymose inflorescences with long peduncles, bracts adnate at the base to the peduncle and flowers with bearded corolla lobes. Hilliard (1991) compared this species with *T. gracile* in the diagnosis, thus this species might be a member of Clade 70. Two of the syntypes of *T. megalocarpum* (*Dinter* 726 and 912) have inflorescences similar to *T. gracile* with lateral cymules of mostly 1--3 flowers and bracts free or shortly adnate to the base of the peduncle, features clearly observed in the duplicates at SAM. The fruits that develop then have long stipes up to 3 mm long. However, in the protologue of *T. megalocarpum,* Hill described the inflorescences as racemose with floral bracts adnate to very short pedicels [peduncles]. Hilliard (2006) also described the inflorescences as racemose with completely adnate bracts. Unfortunately, she selected *Dinter* 585 (K), a poor specimen without inflorescences, as the lectotype. Further studies are required to clarify the taxonomic and phylogenetic relationships of this species that could be part of Clades 70 or 78. *Thesium welwitschii* Hiern, a species described from Angola, has similar cymose inflorescences with lateral monochasia and dichasia and floral bracts only partially adnate to the peduncles. *Thesium burchellii* was reduced to a synonym of *T. megalocarpum* by Hilliard (2006), but comparing the type material and geographical distribution they might be different species, in agreement with Lombard & Le Roux (2023).

**71. *Alatum*-*Costatum* Clade (BIPP 0.63).** This clade (Fig. 2D) includes species from South Africa or Tropical Africa except *T. pseudocystoseiroides*, from Madagascar. It was split into Clades 72 and 73, although with low or very low support and not resolved in the chloroplast analyses. The majority of species in Clade 71 have external glands between the corolla lobes. These can be quite conspicuous (e.g., *T. asterias*) or poorly-developed and inconspicuous (e.g., *T. polygaloides*). However, such glands occur outside of Clade 71 (e.g., Clades 61, 64, 69, 70) and even in distant clades such as Clade 30 and a few Eurasian species. This scattered presence indicates that it may be useful for species recognition but overall has low phylogenetic value.

**72. *Alatum* Clade (BIPP 0.79).** The *Alatum* Clade (Fig. 2D) contains two species from Southern Africa, *T. alatum* (Fig. 6A) and *T. utile* A.W.Hill (=*T. cytisoides* A.W.Hill, Fig. 6B), and one species, *T. whyteanum*, from the Chimanimani Mts straddling the border of Zimbabwe and Mozambique, and Mulanje Mt in Malawi. The relation of the latter species with the other two is resolved with low support only in Bayesian analyses (<50% with MPBS). *Thesium whyteanum* plants are shrubs with one or few stems up to 3 m tall branching above with linear leaves densely arranged along the upper part of the stems. The flowers occur in elongated racemose inflorescences of lateral cymules (mostly dichasia) or less frequently as solitary flowers. The bracts are free or adnate to the base of the peduncle, with the basal one longer than the flowers (when solitary) and about as long as the cymules. The relationship of *T. alatum* and *T. utile* was suggested by Hilliard & Burtt (1983). *Thesium alatum* is a more robust plant, with winged stems, corolla lobes twice the length of the hypanthium, and sessile stigmas. While the inflorescence of *T. utile* is more similar to that of *T. whyteanum*, *T. alatum* has mostly solitary flowers along the inflorescence axis with the bracts shorter than the flowers and adnate to the peduncle. Although the phylogenetic relationship of these two species was resolved with good support with ITS, it was not resolved with chloroplast sequences. Our accession number 4710 was identified on the herbarium sheet as *T. gracile* by O. Hilliard; it is a sterile specimen, but vegetative features are more similar to *T. utile* despite the fact that this individual was collected in the Nyanga Mts (Zimbabwe), north of the known distribution area of *T. utile* which is mostly South Africa.

**Fig. 6.**
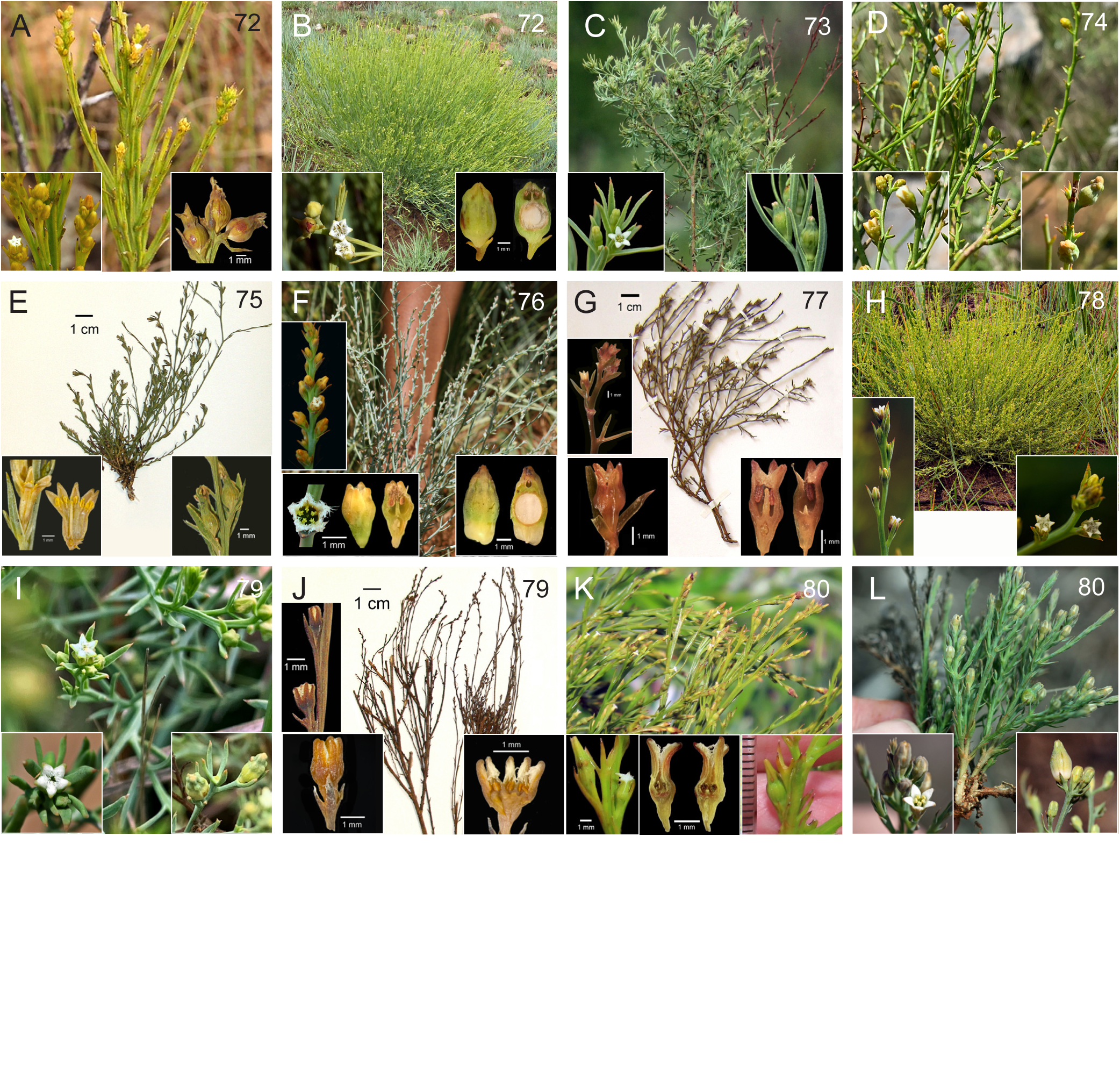
Photographs of representatives of the major clades of *Thesium* species. The numbers in the upper right corners correspond to clade numbers on the Bayesian trees (Fig. 2D). L.S = longitudinal section. **A,** *Alatum* clade, *T. alatum*, flowering shoots, flowers, fruits (*Goetghebeur 4512*, BR); **B,** *Alatum* clade, *T. utile*, habit, flowers, fruit L.S.; **C,** *Davidsoniae* clade, *T. davidsoniae*, habit, flower, fruits; **D,** *Multiramulosum* clade, *T. multiramulosum*, shoots with flowers and fruits; **E,** *Virens* clade, *T. subsimile*, herbarium specimen, rehydrated and dissected flowers, and fruit (*Balkwill & Balkwill 11245*, MO); **F,** *Nigrum* clade, *T. magalismontanum*, fruiting plant, inflorescence, flowers, fruit L.S.; **G,** *Lobelioides* clade, *T. lobelioides*, herbarium specimen and rehydrated and dissected flowers (*Brenen 14920*, UPS); **H,** *Goetzeanum* clade, *T. goetzeanum* habit, inflorescence, flowers; **I,** *Gracilarioides* clade, *T. gracilarioides*, flowering shoots, flower, young fruits; **J,** *Gracilarioides* clade, *T. polygaloides*, herbarium specimen and rehydrated and dissected flowers (*Hobson 442*, MO); **K,** *Costatum* clade, *T. asterias*, flowering shoots, flower, flower L.S., and young fruits; **L,** *Costatum* clade, *T. costatum*, excavated plant, flower, fruit.

**73. *Davidsoniae* Clade (BIPP 0.61).** This very poorly supported clade (Fig. 2D) includes Clades 74--80, resolved with varying degrees of support. It is composed of a polytomy involving 36 acessions of 21 species and four unidentified to species. Four species considered part of the *T. goetzeanum* complex by Visser & al. (2018) occur in this clade: *T. goetzeanum,*

4. *T. gracilarioides* A.W.Hill*, T. gypsophiloides* A.W.Hill, and *T. lobelioides* A.DC. Four species considered synonyms of *T. goetzeanum* by Visser & al. (2018) also occur in Clade 73: *T. coriarium* A.W.Hill*, T. nigrum* A.W.Hill and *T. orientale* A.W.Hill (Clade 76). Because these three species are not resolved within Clade 78, we consider them distinct species. Two South African species, *T. davidsoniae* Brenan (Fig. 6C) and accession 4711 of *T. gypsophiloides* are not resolved in subclades but appear as individual components of the polytomy. The former was classified in *T.* sect. *Imberbia* and is a dolomite endemic from E Transvaal with corolla lobes shorter than the tube. It is a densely leafy perennial herb with profuse branching above. The latter is found in South Africa (KwaZulu-Natal) and Eswatini. Other accessions of *T. gypsophiloides* are resolved in Clade 79. For features that distinguish this species from *T. gracilarioides*, see Visser & al. (2018).

**74. *Multiramulosum* Clade (BIPP 1).** This well-supported clade (Fig. 2D) contains *Thesium multiramulosum* and *T. pseudocystoseiroides*. The former is a widespread species from South Africa to Eswatini, Malawi, Zimbabwe, and Mozambique. The latter is endemic to Madagascar. Both are shrubs with flowers placed mostly singly in the axils with bracts and bracteoles shorter than the flowers.

**75. *Virens* Clade (BIPP 1).** This well-supported clade (Fig. 2D) includes *Thesium subsimile* N.E.Br. (Fig. 6E) and *T. virens* E.Mey. ex A.DC., two species with long hypanthial tubes and long styles. Inflorescences are spike-like (but with terminal flowers) and the floral bract is fully recaulescent along the length of the peduncle. Hairs may (e.g. *T. subsimile*) or may not be present on the margins of the corolla lobes. The stems are more or less erect from a branched rootstock and are sulcate via raised vascular bundles. A marked midrib is present on the leaves and bracts.

Unsampled species with these morphological features that might be phylogenetically related include *T. breyeri*, *T. densum* N.E.Br., *T. gracilentum* N.E.Br., *T. hirsutum* A.W.Hill (might be also related to species in Clade 49), *T. nationae* A.W.Hill, *T. racemosum* and *T. resinifolium* N.E.Br. Most of these species have papillate margins on the corolla lobes.

**Clades 76--78**. Species resolved in Clades 76, 77 and 78 share some of the morphological features of the *T. goetzeanum* complex. They all have racemose inflorescences with solitary lateral flowers or with lateral cymes of 1--3 flowers, bracts adnate to more than 2/3 the length of the peduncle and styles of variable length but never sessile or subsessile. As pointed out by Visser & al. (2018), some characters that were used to segregate species are variable including flower size, leaf and bract shape and texture, leaf and bract margins and the presence or absence of glands between corolla lobes. We agree that these characters, being so variable, present problems in recognizing species based solely on morphological features, especially when studying a limited number of herbarium specimens. Molecular data are suggesting that convergence of morphological features might be contributing to the taxonomic difficulties of this group. On the other hand, differences in ecology, age of the plant and maybe even hosts contribute to increasing the infraspecific morphological variation, thus making it difficult to differentiate convergence and variability. One clear example is *T. resedoides*, a species that shares many features with *T. goetzeanum* but was resolved as more distantly related in our phylogenies.

**76. *Nigrum* Clade (BPP 0.84).** Clade 76 (Fig. 2D) received moderate support with ITS and here includes six named species (*T. dissitum* N.E.Br., *T. lesliei* N.E.Br., *T. magalismontanum*, *T. coriarium*, *T. nigrum*, and *T. orientale*) as well as two unidentified taxa. The three latter species were considered synonyms of *T. goetzeanum* (Clade 78) by Visser & al. (2018) and all share the conspicuous external glands between the corolla lobes.

*Thesium dissitum*, a species known only from the Flora Zambesiaca area, has inconspicuous external glands on the flowers and long, narrow floral bracts fused the entire length of the peduncle. This inflorescence is similar to those in the *T. goetzeanum* group. Among the specimens of *T. goetzeanum* resolved in Clade 78, accession 5250 resembles *T. dissitum* for its narrow bracts much longer than flowers and fruits, supporting the view of Visser & al. (2018) that this character is variable in the species. Hilliard (2006) suggested the relationship of *T. dissitum* with *T. mukense* but it is not supported either by nuclear nor by chloroplast sequences. In our ITS phylogeny, *T. dissitum* is resolved as sister to accession 4807, a plant with much shorter bracts and peduncles. The plant is somewhat scabrid and both leaves and bracts are strongly recurved, having characters of both *T. goetzeanum* and *T. resedoides*. *Thesium lesliei* has flowers in dense spike-like or racemose inflorescences with very short peduncles. It presents all the characters of the *T. goetzeanum* species complex but was neither recognized nor included as a synonym by Visser & al. (2018), possibly because this species is somewhat scabrid. It was compared to *T. orientale* by Brown (1932) but *T. lesliei* lacks visible perianth glands and is a shorter plant (<15 cm) with numerous erect stems growing from a rhizome. Type specimens and the plant sampled for this study were collected around Johannesburg (South Africa).

*Thesium magalismontanum* (Fig. 6F) is a South African species (occurring mostly north of the Vaal River) with the indeterminate spike-like inflorescences and other characters shared by the *T. scirpioides* complex (Lombard & al., 2021). However, it was not included in that complex, probably because it may have developed leaves (not scales) at the base of young plants. The individuals of *T. magalismontanum* studied by us have flexuose instead of short and straight placental columns.

One of the two unidentified taxa (accession 5308) has well-developed external glands but differs from other members of the clade vegetatively. It is a densely leafy plant, with short acicular leaves with sharp points that are reminiscent of *Cryptomeria japonica* D.Don. Its inflorescences are at the ends of vegetative shoots that terminate in branched cymes and lateral cymules of 1--3 flowers. The bracts are fused to the peduncles, the flowers bear dense apical beards, and the hypanthium is longer than the corolla lobes and long style.

**77. *Lobelioides* Clade (BPP 1).** In Clade 77 (Fig. 2D), our sample 4817 identified here as *T. lobelioides* (Fig. 6G), was included by Visser & al. (2018) in the list of specimens of *T. resedoides*. Our phylogenies did not resolve this specimen in the *Resedoides* Clade (69) with either nuclear and chloroplast markers. This individual is very similar to the type of *T. lobelioides*, with recurved leaves with acute apices, and the corolla lobes that are 1.4--1.6 mm long, within the range given by Visser & al. (2018) for the species. However, other characters are out of the range given by these authors. Also resolved in Clade 77 is our accession 4709, a possible new species that is a short plant (below 20 cm high) with very broad bracts (c. 2 mm), much longer than the flowers. The flowers are solitary or disposed in lateral cymules (dichasia) with the floral bract longer than the cymules. The flowers are similar to those in *T. asterias* and also *T. lobelioides* but in this specimen the style is ca. 0.8 mm, not absent as in *T. asterias* and shorter than the sizes given for *T. lobelioides* by Visser & al. (2018). Corolla lobes are up to 2.5 mm long much longer than the hypanthium, out of the range given by Visser & al. (2018) for *T. goetzeanum*.

**78. *Goetzeanum* Clade (BIPP 1).** In our sampling, individuals with typical features of *T. goetzeanum* (Fig. 6H) are resolved in Clade 78 with the highest support (Fig. 2D). Some of the individuals lack the vegetative shoots overtopping the highest inflorescences, a character that helps one distinguish this species from *T. resedoides* (Visser & al. 2018). This character might be related to the development of the individual but apparently such vegetative shoots are never present in *T. resedoides*.

**79. *Gracilarioides* Clade (BPP 0.62).** This very poorly supported clade (Fig. 2D) includes *T. gracilarioides* (Fig. 6I) and *T. gypsophiloides*, two species of the *T. goetzeanum* complex with very short styles or sessile stigmas and anthers inserted on a short hypanthium. Inflorescences in *T. gypsophiloides* are mostly cymose whereas in *T. gracilarioides* they are racemose. Other morphological differences between these two species are described in detail by Visser & al. (2018). Accession 4711 (part of the polytomy of Clade 73) has morphological features of *T. gypsophiloides* including the short hypanthium and almost sessile stigma but is not resolved in Clade 79. Conversely, chloroplast sequences resolved this accession in a highly supported clade with *T. polygaloides* of Clade 79. Flowers in *T. polygaloides* also have a short hypanthium and the stigma is almost sessile, like in *T. gracilarioides* and *T. gypsophiloides*, but the branching is virgate above and the inflorescences are elongated and spike-like, lacking an elongated peduncle (Fig. 6J). The species has been collected growing in a marsh at sea level, not in open grasslands or steep slopes. The type of *T. polygaloides* was also collected in a marsh, an uncommon habitat for the genus, and apparently this species might be restricted to wetland habitats of the KwaZulu-Natal seaboards. With chloroplast sequences both 4711 and *T. polygaloides* were resolved in a well-supported clade with *T. multiramulosum* and the Madagascan *T. pseudocystoseiroides* (ITS Clade 74).

Also resolved with low support in Clade 79 is accession 5237, a possible undescribed species. This plant lacks an apical beard but has developed hairs on the margins of the corolla lobes. According to Hill’s classification it would be included in *T.* subsect. *Fimbriata*. The inflorescence is racemose with the bracts totally adnate to the peduncle, the styles are short (< 1 mm), and the hypanthium is much shorter that the corolla lobes (c. 2 mm). The morphology of bracts and leaves, together with short style are characters shared with *T. gypsophiloides*.

**80. *Costatum* Clade (BIPP = 0.76).** This poorly supported clade (Fig. 2D) includes four South African species and one tropical African species (*T. brevibarbatum*). One subclade formed by *T. brevibarbatum*, *T. costatum* A.W.Hill (Fig. 6L), and *T. inversum* N.E.Br. received the highest support. The latter two species share several morphological features including woody stolons, ribbed or angular erect stems, acute leaves, flowers mostly in cymules, and bracts with prominent midribs that are adnate to below half the length of the peduncle (or fully fused on upper solitary flowers). Several species or varieties have been described as morphologically similar to *T. costatum*, but further taxonomic studies might prove them to be different growth forms of a very variable species in terms of size, branching of the inflorescence (i.e., *T. costatum* var. *paniculatum* N.E.Br.), and density and length of leaves and bracts.

*Thesium asterias* (Fig. 6K) has elongated spike-like inflorescences, frequently with lateral cymules of up to three flowers. It is very recognizable by its elongated flower buds with prominent external glands, short hypanthium, long corolla lobes with apical beards, and sessile stigmas. *Thesium disparile* was resolved in a subclde with *T. asterias* but with only moderate support, and these two species are very different morphologically.

Unsampled species include *T. cymosum* A.W.Hill which is mostly distributed in highlands and montane areas of Malawi and around the border of Zimbabwe and Mozambique. It was considered to be closely related to the South African *T. costatum* and *T. exile* N.E.Br. by Hilliard (2006). *Thesium fenarium* A.W.Hill from Malawi could be related to these species or it is a synonym of *T. cymosum* as suggested by Hilliard (2006). Brown (1932) described several new species of the genus, nine of them in *Thesium* sect. *Imberbia*. Important features that Brown used to separate the species included plant height, inflorescence (spike-like, racemose, pedunculated cymes or in dense clusters), inflorescences overtopped or not with sterile branches and the shape and size of the hypanthium and corolla lobes. Brown’s unsampled species of *T.* sect. *Imberbia* that might be resolved in Clade 80 or in other parts of Clade 65 include: *T. gracilentum*, *T. resinifolium*, *T. breyeri*, *T. densum* and *T. exile*. More recently, Brenan (1979) described *T. jeaniae* Brenan a species possibly related to *T. costatum* or *T. exile* but with angular stems, rigid leaves and bracts with marked midribs terminating in a spiny apex, and short spike-like inflorescences, most of them reduced to a terminal flower.

### Biogeography of *Thesium*

The extensive sampling in this study throughout the entire distributional area of *Thesium* (Fig. 1) allows the reconstruction of the historical biogeography of the genus. Further studies including estimation of absolute divergence times will help to understand the dispersal and speciation events within the genus and allow comparison with the large-scale floristic patterns observed in other groups of African origin.

The monotypic genus *Lacomucinaea* has a SW African distribution (Nickrent & García, 2015) and *Osyridicarpos* is more common in SE Africa with populations extending north to Ethiopia along the Eastern Arc African Mountains (EAAM). Both genera are the extant lineages most closely related to *Thesium*. Moore & al. (2010), using ancestral range reconstruction, suggested a Southern African origin of *Thesium* in the Late Eocene, with a crown age of 39.1 ± 11.9 My. *Lacomucinaea* and *Osyridicarpos* were not included in their analyses, but their current distribution supports their common origin in Southern Africa. The topology of both ITS and chloroplast trees resolves, with the highest support in all the analyses, two main lineages in the genus, the CMPB lineage (Clade 2), mostly including the Eurasian species, and the African lineage (Clade 26).

#### Biogeography of the CMPB lineage

The CMPB lineage has a remarkable disjunct distribution pattern because it includes the Cape endemic species of *T.* subg. *Hagnothesium* (former genus *Thesidium*) nested within Mediterranean, Canarian, and Eurasian species (*T.* subg. *Thesium*), not within the main African lineage. Both chloroplast and ITS phylogenies highly support this relationship.

Moore & al. (2010) dated the divergence of both subgenera to the Oligocene (33.7--23.8 Mya), preceding other plant groups with origin in Southern Africa that dispersed north via East Africa with the Late-Miocene uplift of the EAAM (Pokorny & al., 2015).

The early diverging lineages in the type subgenus include the NW African *T. mauritanicum* and *T.* sect. *Kunkelliela*, endemic to the Canary Islands. Even though *T. mauritanicum* might be the closest extant relative of *Kunkeliella,* their sister relationship is not resolved in our phylogenies. Our data indicate that *T.* sect. *Kunkeliella* is monophyletic and its presence in the Canary Islands is the result of a single dispersal event followed by local *in situ* diversification in the islands with one species in Gran Canaria, three in Tenerife and one in La Palma (Rodríguez-Rodríguez & al., 2022).

A second lineage, the Paleoboreal Clade (Clade 7), is well-supported with both ITS and chloroplast markers and includes the remainder species of *T.* subgen. *Thesium*. Well-supported backbone relationships in this lineage are not always seen, but among the well-supported clades the core Eurasian Clade (12), that includes most of the diversity in terms of number of species, emerges as an independent lineage with a mostly W Asian distribution. The ITS phylogeny shows that *T. humile*, a circum-Mediterranean and Canarian species, is moderately supported as sister to the *Macranthia* Clade (9), which includes species distributed in the E Mediterranean region and W Asia. A second lineage, the *Procumbens* Clade (11), is present in W Asia and SE Europe and finally, the C Asian *T. minkwitzianum* whose closest relatives are not resolved. This scenario suggests that after the migration from the Cape region to the Mediterranean, one or several dispersals to W Asia and subsequent speciation generated the extant diversity of species of *Thesium* with mostly E Mediterranean and W Asian distributions, especially in Anatolia and other regions of the Middle East.

The Core Eurasian Clade (12) represents the dispersal and subsequent speciation in Europe that produced many of the endemic species of the continent, especially in southern mountain areas. In the ITS phylogeny, the *Linophylla* Clade (17) together with *T. catalaunicum* and *T. rostratum* are the closest European relatives of a group of species of Asian distribution. However, chloroplast markers resolve the *Linophylla* Clade as being more closely related to other European species. Independent of the markers used, the phylogenies within the Core Eurasian Clade are congruent with the concept of one to several dispersals from Europe into Asia and subsequent speciation in Siberia, C Asia, China, and the Himalayas. This represents the second Asian radiation and subsequently migration into E Australia (*T. australe*).

#### Biogeography of the African lineage

The topology of ITS and chloroplast trees resolves three early diverging lineages (Clades 28--30) including species with current distributions in the Succulent Karoo and Nama-Karoo semi-deserts (Clade 29) and Fynbos (Clades 28 and 30). One species, *T. triflorum*, is also present in areas of the East African mountains, probably as the result of a later dispersal to the north from South Africa (Brummitt, 1976). Mountains of the Cape Fold Belt were important for the further diversification of *Thesium* in the Cape as revealed by the phylogenetic position of *T. nautimontanum*, an endemic from the Matroosberg Mt (García & al., 2018), as sister to the entire African Clade in the ITS phylogeny or in the chloroplast phylogeny as sister to the African species (with the exception of Clades 28--30 that were resolved as earlier diverging lineages). The rapid diversification in the Fynbos between the Great Escarpment and the coast generated the current diversity of many endemic species, all of which are included *T.* subgen. *Frisea* (Clade 32).

Both nuclear and chloroplast phylogenies resolved subgenus *Psilothesium* (Clade 48) as sister to the Core Cape Clade, representing the broad diversification of the genus in the grassy biomes of E and N South Africa and the radiation into Tropical Africa. The ITS topology resolves the *Gnidiaceum* Clade (49) as sister to the rest of the subgenus. This clade includes species from the SW ranges of the Cape Fold Belt (*T. oresigenum* and *T. sp nov.* 5574) as sister to others from the SE Great Escarpment and E Cape Fold Belt, supporting the biogeographical link between both areas (Clark & al., 2011, 2012). Although the topology of the chloroplast phylogeny does not place *T. oresigenum* and *T. sp. nov.* 5574 in the *Gnidiaceum* Clade, both species were resolved as successive sisters to the grassland and tropical African radiation.

Subsequent radiation in the Tropics is, in part, represented by Clades 52 and 60 that include species lacking an apical beard and mostly distributed in savannas in regions spanning East and West Africa. Our data suggest at least two long-distance dispersals in Clade 52, one to South America and at least one to Madagascar. The monophyly of the South American species suggests a single long-distance dispersal from tropical Africa. Although we only have sequence data of *T. madagascariense*, the morphological features typical of the Brachyblast Clade (54) suggest that another Madagascan endemic species, *T. perrieri*, belongs in this group. Because the latter could not be sampled it is unknown whether the presence of these two species in Madagascar is the result of one or two independent dispersal events from the African continent.

Although not well-supported, the topologies of ITS and chloroplast trees suggest an independent diversification of the leafy species of subgenus *Psilothesium* in the grass-dominated biomes of Southern Africa with further range expansion to the Tropical Africa. The chloroplast phylogenies highly support these species as monophyletic and with lower support its sister relationship with *T.* sect. *Barbata* species of the *Gnidiaceum* Clade and other species mostly distributed around the Drakensberg mountains. This area has been considered a “stepping-stone” between the CFR and the so-called Afrotemperate Region (White, 1978; Galley & al., 2007) encompassing high elevations of mountain ranges of Southern and East Africa (and to a minor extent Central and West Africa). Migration northwards is supported by the presence in this clade of species endemic to East African mountains and highlands such as *T. kilimandscharicum*, *T. dolichomeres*, *T. whyteanum*, etc. and its expansion to Ethiopia and SE of the Arabia Peninsula (species such as *T. radicans* or *T. hararensis*). Expansion and diversification took place also in lowland areas of the subtropical and tropical grasslands of SE Africa and the tropical Central and West Africa as well as more recent *in situ* speciations suggested by the overlap of morphological characters among species in this taxonomically difficult group. Further events of long-distance dispersal are seen in Clade 64, one to Madagascar (*T. decaryanum*) and another one to E and SE Asia (*T. psilotoides*). *Thesium wightianum* (unsampled) from SE India, shares morphological characters with *T. radicans* and molecular data from this species might reveal another case of dispersal to Asia from NE Africa in this clade (see discussion of Clade 64).

### Conflict between the ITS and chloroplast datasets

Although gene trees derived from the two subcellular compartments were generally congruent, ITS and chloroplast trees showed remarkable incongruences at the specific or even infraspecific levels. These incongruences are not observed equally along the main lineages. The Eurasian, grassland and tropical African lineages show, in general, high congruence between datasets with particular exceptions previously discussed. However, and in agreement with the results obtained by Zhigila & al. (2020), the Cape radiation (*T.* subgen. *Frisea*) is characterized by the high incongruence of chloroplast and nuclear trees. Within the subgenus this incongruence is not detected in the monophyletic sect. *Frisea* (Clade 39) because of the lack of internal resolution in this clade but it is interesting that none of the species of this section are resolved among other clades of the Cape radiation on the chloroplast trees except for accession 5460. Biological factors known to cause phylogenetic incongruence include paralogy, hybridization, incomplete lineage sorting, and horizontal gene transfer (HGT).

*Paralogy* cannot explain the incongruences detected because there is very low ITS intraspecific variation and in those cases with polymorphic ribotypes the paralogous sequences obtained after cloning were closely related to each other. Moreover, clades resolved for subgen. *Frisea* in the ITS trees largely correspond with morphological features and species are mostly resolved as monophyletic, unlike in the chloroplast phylogeny.

*Hybridization* has been identified as one of the most common sources of incongruence between nuclear and plastid phylogenies. Although hybridization probably occurs in *Thesium*, it has yet to be demostrated and by itself cannot explain the widespread incongruences found in the Cape Clade. Hendrych (1972) discussed three cases described in the literature as possible interspecific hybrids and his conclussion was that hybrid origin could not be proven. We sequenced cpDNA from two or more individuals in 25 species of subgenus *Frisea* and 14 of them were resolved as polyphyletic on the chloroplast but not on the ITS trees. They were resolved in two or more well-supported clades together with individuals of species very different in vegetative, inflorescence, floral and fruit morphology, thus ruling out the introgression of nuclear genes. As an example, *T. strictum*, a common and widespread species in the Fynbos region, is distinctive given its wand-plant architecture and other morphological features. It was resolved as monophyletic with strong support on the ITS tree (Clade 36). In contrast, on the chloroplast trees these accessions were placed among several well-supported clades with species such as *T. pubescens*, *T. pycnanthum*, *T. hispidulum*, and *T. prostratum*, all morphologically quite different. If hybridization and successive backcrosses were involved, without convergent evolution of the nuclear ribosomal cistron, one would expect to find cases of ITS sequence polymorphisms with ribotypes of the same species resolved on different clades on the ITS trees. In the case of convergent evolution, it should have occurred in all the species and always homogenized towards the paternal progenitor because species are monophyletic on the ITS trees and clades are congruent with morphology.

*Stochastic incomplete lineage sorting of ancestral polymorphisms* might explain some incongruences between nuclear and chloroplast phylogenies, especially under high rates of diversification. This biological factor was considered by Zhigila & al. (2020) as the most probable explanation of the incongruences in *T.* subgen. *Frisea*. This hypothesis is supported by the fact that incongruences seem random and not geographically biased. However, it is difficult use incomplete lineage sorting to explain the incongruence found by Zhigila & al. (2020) of one individual of *T. minus* (subgenus *Hagnothesium*) resolved in subgenus *Frisea* on their concatenated chloroplast tree. It is more probable that the cpDNA in this individual was acquired after the diversification of the four subgenera. To our knowledge no cases have been documented in plants of maintenance of ancestral polymorphism long after speciation or at least with such a wide sequence diversity among individuals of the same species as we found in subgenus *Frisea*. In this work we have detected polyphyletic cpDNA haplotypes among individuals of different populations of the same species, but it remains to be studied if it occurs also within the same population. Although incomplete lineage sorting in subgenus *Frisea* or even deep reticulation might be involved, it alone cannot explain the incongruences found in subgenus *Frisea* by Zhigila & al. (2020) and in this work.

*Horizontal Gene transfer (HGT)* is recognized to have contributed to the genome plasticity and adaptive evolution of prokaryotes and unicellular eukaryotes, and played an important role in the origin and evolution of green plants (Chen & al., 2021). HGT has been documented to be frequent in several groups of parasitic plants (Davis & Xi, 2015). Normally HGT involves particular genes but these transfers can be large, such as the acquisition of entire mitochondrial genomes from three green algae, one moss and fragments of other angiosperms in *Amborella* (Rice & al., 2013). Although HGT of mitochondrial genes is frequent, especially in parasitic plants, it only rarely affects plastid genomes (Sánchez-Puerta, 2014). We sequenced two regions of the chloroplast genome from different locations. The topologies recovered with each of them are largely congruent (see e.g., Figs. S8 and S11) and similar results were obtained by Zhigila & al. (2020) comparing their *trnLF* and *mat*K phylogenies. This suggests that whole plastomes, and not just fragments, have been transmitted horizontally between species. Chloroplast capture in plants is typically associated with reticulation events. However, asexual chloroplast capture in the absence of nuclear introgression has been demonstrated to occur through natural grafts (Stegemann & al., 2012). Plastids can dedifferentiate into motile ameboid organelles and move into neighboring cells through connective pores after cell wall disintegration (Hertle & al., 2021). In parasitic plants, intimate connections between cells of the host and parasite are formed during haustoria development, which mechanistically resemble grafting. In *Thesium*, autoparasitism through self-haustoria and connections between different individuals are known to be common (Melnyk & Meyerowitz, 2015). Subgenus *Frisea* is greatly diversified, and several species commonly grow in the same area in close proximity to each other. This might allow the formation of haustorial connections among individuals of different species, thus in theory HGT of whole plastids could occur between them. *Thesium* offers an attractive group in which to investigate whether this phenomenon is actually taking place and whether it explains the unprecedented level of incongruence documented here between nuclear and chloroplast gene trees.

## Acknowledgements

We are grateful to the curators of the following herbaria for the access to their collections and granting permission for DNA extractions: BR, LE, MA, MO, NY and UPS. Loans for the revision of morphological features were obtained from BRLU, LG, NBG, NU, NH, and POZG. This work was partially supported by grant CGL2006-00300/BOS from the Consejo Superior de Investigaciones CSIC, Real Jardín Botánico (Madrid) and the contract SOLAUT_00033574 of the Spanish Ministry of Science and Innovation to MAG. Some of the accessions from Spain, Italy, Morocco, and Turkey, used for DNA extraction and sequencing, were sampled during field trips for the project *Flora iberica*. We are very grateful to V.R. Clark, E. Pienaar, J. Šibík and I. Šibíková for providing collections from South Africa. Thanks to Dagmar Mucina for her assistance during our field work in South Africa in 2007 and to Ginés L. González for his contribution in our collecting trip and providing photographs. LM appreciates the logistic support of the Iluka Chair in Vegetation Science & Biogeography at the Murdoch University, Perth, Western Australia. Many of the photographs used to prepare Figs. 3--6 were obtained under the Creative Commons Licence from the following web pages: iNaturalist, Phytoimages, Biodiversity Data Bank of the Canary Islands, Plant Biodiversity of South-Western Morocco, Acta plantarum, Encyclopedia of Life, Wikimedia Commons, Tropicos and GBIF. The authors of those images are listed in Appendix S5.

## Appendix 1. GenBank accession numbers and voucher specimen information

***Taxon*** and authorship, DNA extraction number, Country, *Collector(s)* plus *collection number* (herbarium code), ITS, trnLF, trnDT (“--“ indicates missing sequence). All GenBank accession numbers starting with “OP” were newly generated for this work.

***Lacomucinaea lineata*** (L.f.) Nickrent & M.A.García, DLN 4413, South Africa, *K. Balkwill & L. McDade 11765* (MO), KP318956, --, OP171945; DLN 4725, South Africa, *E. Coppejans 1016* (BR), KP318957, --, --; DLN 4726, South Africa, *P. Goldblatt 6526* (BR), KP318958, --, --; DLN 4800, South Africa, *P. Bamps 9427* (BR), KP318959, --, --; DLN 5509, South Africa, *L. Mucina 020906/12* (MA), KP318960, KP318970, OP171946; ***Osyridicarpos schimperianus*** A.DC., DLN 4110, South Africa, *D.L. Nickrent 4110* (BH), KP318955, KP318968, OP171947; ***Thesium acuminatum*** A.W.Hill, DLN 5488, South Africa, *M.A. García & al. 5488* (MA, BH, NBG), OP184088, OP172214, OP171948; ***T. acutissimum*** A.DC., DLN 5585, South Africa, *V. R. Clark & al. 46* (GRA), OP184089 (ITS clone A), OP184090 (ITS clone B), OP172215, OP171949; ***T. aellenianum*** Lawalrée, DLN 4682, Democratic Republic of the Congo, *F. Malaisse 13854* (BR), OP184091, OP172216, OP171950; ***T. aggregatum*** A.W.Hill, DLN 4812, South Africa, *J. Acocks 16887* (UPS), OP184092, --, --; DLN 5486, South Africa, *M.A. García & al. 5486* (MA, BH, NBG), OP184093, OP172217, OP171951; DLN 5514, South Africa, *L. Mucina 090306/11* (MA), OP184094, OP172218, OP171952; ***T. alatavicum*** Kar. & Kir., DLN 5099, Kazakhstan, *Popov & Rubtzov 4111* (LE), OP184095, OP172219, OP171953; DLN 5100, Kazakhstan, *V.P. Goloskokov s.n.* (LE), OP184096, OP172220, OP171954; DLN 5101, Kyrgyzstan, *Aydarova s.n.* (LE), OP184097, OP172221, --; DLN 5102, Uzbekistan, *R.V. Kamelin 48* (LE), OP184098, --, OP171955; DLN 5164, China, *G. Ke-Isen 955* (LE), OP184099, OP172222, OP171956; ***T. alatum*** Hilliard & B.L.Burtt, DLN 4684, South Africa, *P. Goetghebeur 4512* (BR), OP184100, OP172223, OP171957; DLN 5306, South Africa, *O. Hilliard 5113* (MO), OP184101, OP172224, --; ***T. albomontanum*** Compton, DLN 5376, South Africa, *M.A. García & al. 5376* (MA, BH, NBG), OP184102, OP172225, OP171958; DLN 5569, South Africa, *E. Pienaar M380* (MA), OP184103, OP172226, OP171959; ***T. alpinum*** L., DLN 4835, Spain, *G. López & al. 10027GL* (MA), OP184104, OP172227, OP171960; DLN 5128, Russian Federation, *G. Konechnaya & E. Litvinova s.n.* (LE), OP184105, OP172228, OP171961; ***T. amicorum*** Lawalrée, DLN 4683, Democratic Republic of the Congo, *S. Lisowski & al. 5736* (BR), OP184106, OP172229, OP171962; ***T. angulosum*** A.DC., DLN 5223, South Africa, *A. Pegler 176* (MO), OP184107, --, --; ***T. aphyllum*** Mart. ex A.DC., DLN 4850, Brazil, *Hatschbach 20060* (NY), OP184108, OP172230, OP171963; DLN 4851, Brazil, *Irwin & al. 12166* (NY), OP184109, --, --; ***T. asterias*** A.W.Hill, DLN 5224, South Africa, *A. Nicholas & L. Smook 2328* (MO), OP184110, OP172231, OP171964; ***T. auriculatum*** Vandas, DLN 5157, Albania, *A. Baldacci 1900* (LE), OP184111, OP172232, --; ***T. australe*** R.Br., DLN 5225, Australia, *A.B. Costin s.n.* (MO), OP184112, OP172233, OP171965; ***T. bavarum*** Schrank, DLN 4836, Italy, *C. Navarro & al. 4327* (MA), OP184113, OP172234, OP171966; DLN 5158, Italy, *A. Charpin & al. 12* (LE), OP184114, --, --; ***T. bergeri*** Zucc., DLN 5159, Turkey, *W. Siebe s.n.* (LE), OP184115, OP172235, --; ***T. bertramii*** Aznav., DLN 4849, Turkey, *J.J. Aldasoro & al. 2693* (MA), OP184116, OP172236, OP171967; ***T. brachyphyllum*** Boiss., DLN 5192, Armenia, *L. Medina & al. LM-2549* (MA), OP184117, OP172237, OP171968; ***T. brasiliense*** A.DC., DLN 4852, Brazil, *Irwin & al. 24645* (NY), OP184118, OP172238, OP171969*; **T. brevibarbatum*** Pilg., DLN 5228, Zimbabwe, *E. Best 1719* (MO), OP184119, OP172239, --; ***T. capitatum*** L., DLN 5411, South Africa, *M.A. García & al. 5411* (MA, BH, NBG), OP184120, OP172240, OP171970; ***T. capitellatum*** A.DC., DLN 4780, South Africa, *J. Leonard 7351* (BR), OP184121, OP172241, OP171971; DLN 5387, South Africa, *M.A. García & al. 5387* (MA, BH, NBG), OP184122, OP172242, OP171972; DLN 5441, South Africa, *M.A. García & al. 5441* (MA, BH, NBG), OP184123, OP172243, OP171973; DLN 5564, South Africa, *E. Pienaar M856* (MA), OP184124, --, --; DLN 5573, South Africa, *E. Pienaar M266* (MA), OP184125, OP172244, OP171974; DLN 5578, South Africa, *E. Pienaar M896* (MA), OP184126, --, --; ***T. capituliflorum*** Sond., DLN 5371, South Africa, *M.A. García & al. 5371* (MA, BH, NBG), OP184127, OP172245, OP171975; DLN 5455, South Africa, *M.A. García & al. 5455* (MA, BH, NBG), OP184128, OP172246, OP171976; ***T. carinatum*** A.DC., DLN 4094, South Africa, *D.L. Nickrent & al. 4094* (BH), OP184129, --, --; DLN 4813, South Africa, *E. Esterhuysen 19884* (UPS), OP184130, --, --; DLN 5454, South Africa, *M.A. García & al. 5454* (MA, BH, NBG), OP184131, OP172247, OP171977; DLN 5463, South Africa, *M.A. García & al. 5463* (MA, BH, NBG), OP184132, OP172248, OP171978; DLN 5476, South Africa, *M.A. García & D. Nickrent 5476* (MA, BH, NBG), OP184133, OP172249, OP171979; ***T. catalaunicum*** Pedrol & M.Laínz, DLN 4837, Spain, *J. Pedrol 5605JP* (MA), OP184134, OP172250, OP171980; ***T. chinense*** Turcz., DLN 4390, China, *D. Chen 1999-9* (MO), OP184135, OP172251, OP171981; DLN 5153, Russian Federation, *T.I. Neczaeva s.n.* (LE), OP184136, KP318977, OP171982; ***T. cilicicum*** Hausskn.ex Bornm., DLN 4838, Turkey, *F. Muñoz Garmedia & al. 4566* (MA), KP318965, KP318975, OP171983; ***T. commutatum*** Sond., DLN 4441, South Africa, *S.L. Williams 3097* (MO), OP184137, --, --; DLN 5449, South Africa, *M.A. García & al. 5449* (MA, BH, NBG), OP184138, OP172252, OP171984; DLN 5527, South Africa, *L. Mucina 211006/50* (MA), OP184139, OP172253, OP171985; ***T. compressum*** Boiss. & Heldr., DLN 5176, Armenia, *A.M. Barsegyan s.n.* (LE), OP184140, OP172254, OP171986; ***T. coriarium*** A.W.Hill, DLN 5266, Lesotho, *L. Smook 7265* (MO), OP184141, OP172255, --; ***T. cornigerum*** A.W.Hill, DLN 5270, South Africa, *K. Balkwill 5299* (MO), OP184142, OP172256, OP171987; ***T. corsalpinum*** Hendrych, DLN 4839, Corse, *Serra & Bott 4812* (MA), OP184143, OP172257, OP171988; ***T. corymbuligerum*** Sond., DLN 5419, South Africa, *M.A. García & al. 5419* (MA, BH, NBG), OP184390, OP172454, OP172153; DLN 5433, South Africa, *M.A. García & al. 5433* (MA, BH, NBG), OP184391, OP172455, OP172154; DLN 5525, South Africa, *L. Mucina 141006/62* (MA), OP184392, OP172456, OP172155; ***T. costatum*** A.W.Hill, DLN 5234, South Africa, *P. du Toit 23* (MO), OP184144, OP172258, --; DLN 5251, South Africa, *K. Balkwill 10238* (MO), OP184145, OP172259, OP171989; ***T. cruciatum*** A.W.Hill, DLN 4691, South Africa, *H. Merxmüller & W. Giess 3225* (BR), OP184146, OP172260, OP171990; ***T. cupressoides*** A.W.Hill, DLN 4692, South Africa, *P. Bamps 7226* (BR), OP184147, OP172261, OP171991; ***T. cuspidatum*** A.W.Hill, DLN 5444, South Africa, *M.A. García & al. 5444* (MA, BH, NBG), OP184148, OP172262, OP171992; ***T. davidsoniae*** Brenan, DLN 4694, South Africa, *J. Brenan 14931* (BR), OP184149, OP172263, OP171993; ***T. decaryanum*** Cavaco & Keraudren, DLN 5303, Madagascar, *R. Bayer MAD-4054* (MO), OP184150, OP172264, OP171994; ***T. deceptum*** N.E.Br., DLN 5261, South Africa, *K. Balkwill 11316* (MO), OP184151, OP172265, OP171995; DLN 5290, South Africa, *K. Balkwill 11285* (MO), OP184152, OP172266, OP171996; ***T. densiflorum*** A.DC., DLN 5238, South Africa, *W. Hanekom 2737* (MO), OP184153, OP172267, OP171997; DLN 5438, South Africa, *M.A. García & al. 5438* (MA, BH, NBG), OP184154, OP172268, OP171998; ***T. disciflorum*** A.W.Hill, DLN 5592, South Africa, *V.R. Clark & T.W. Naude 153* (GRA), OP184155 (ITS clone A), OP184156 (ITS clone B), OP172269, OP171999; ***T. disparile*** N.E.Br., DLN 5288, South Africa, *K. Balkwill & McCallum 11151* (MO), OP184157, OP172270, OP172000; ***T. dissitum*** N.E.Br., DLN 5240, Zambia, *N. Zimba & al. 939* (MO), OP184158, OP172271, OP172001; ***T. dolichomeras*** Brenan, DLN 4695, Zimbabwe, *P. Bamps & al. 798* (BR), OP184159, OP172272, OP172002; ***T. doloense*** Pilg., DLN 4394, Republic of the Congo, *D. Thomas & al. 8699* (MO), OP184160, OP172273, --; ***T. durum*** Hilliard & B.L.Burtt, DLN 5274, South Africa, *M. Werger 1805* (MO), OP184161, OP172274, --; ***T. ebracteatum*** Hayne, DLN 5137, Russian Federation, *V.N. Tikorov & al. s.n.* (LE), OP184162, OP172275, OP172003; ***T. equisetoides*** Welw. ex Hiern, DLN 4395, Gabon, *L. White & K. Abetnathy 1294* (MO), OP184163, OP172276, OP172004; DLN 4398, Gabon, *M. Sosef & al. 538* (MO), OP184164, OP172277, OP172005; ***T. ericaefolium*** A.DC., DLN 5232, South Africa, *E. Oliver 8507* (MO), OP184165, --, --; DLN 5379, South Africa, *M.A. García & al. 5379* (MA, BH, NBG), OP184166, OP172278, OP172006; DLN 5380, South Africa, *M.A. García & al. 5380* (MA, BH, NBG), OP184167, OP172279, OP172007; DLN 5420, South Africa, *M.A. García & al. 5420* (MA, BH, NBG), OP184168, OP172280, OP172008; DLN 5443, South Africa, *M.A. García & al. 5443* (MA, BH, NBG), OP184169, OP172281, OP172009; DLN 5499, South Africa, *M.A. García & al. 5499* (MA, BH, NBG), OP184170, OP172282, OP172010; ***T. euphorbioides*** P.J.Bergius, DLN 5435, South Africa, *M.A. García & al. 5435* (MA, BH, NBG), OP184171, OP172283, OP172010; ***T. euphrasioides*** A.DC., DLN 5493, South Africa, *M.A. García & al. 5493* (MA, BH, NBG), OP184172, OP172284, OP172012; ***T. fastigiatum*** A.W.Hill, DLN 4397, Tanzania, *R.E. Gereau & al. 6060* (MO), OP184173, --, --; DLN 4433, Zambia, *D.K. Harder & al. 3078* (MO), OP184174, OP172285, OP172013; DLN 4700, Tanzania, *S. Bidgood & al. 1287* (BR), OP184175, --, --; DLN 4795, Malawi, *P. Bamps 8971* (BR), OP184176, OP172286, OP172014; ***T. ferganense*** Bobrov, DLN 5107, Kyrgyzstan, *O. Knorring & Z. Minkwitz 317* (LE), OP184177, OP172287, --; ***T. filipes*** A.W.Hill, DLN 4701, Democratic Republic of the Congo, *S. Lisowski B-6951* (BR), OP184178, OP172288, --; ***T. fimbriatum*** A.W.Hill, DLN 5245, Tanzania, *P. Goldblatt & al. 8056* (MO), OP184179, OP172289, OP172015; ***T. flexuosum*** A.DC., DLN 4715, South Africa, *P. Goldblatt 1772* (BR), OP184180, OP172290, OP172016; DLN 5246, South Africa, *P. Bohnen 4921* (MO), OP184181, OP172291, --; DLN 5395, South Africa, *M.A. García & al. 5395* (MA, BH, NBG), OP184182, OP172292, OP172017; ***T. foliosum*** A.DC., DLN 5248, South Africa, *P. Goldblatt 2077* (MO), OP184183, OP172293, OP172018; ***T. fragile*** L.f., DLN 4102, South Africa, *D. Nickrent & A. Wolfe 4102* (BH), OP184184, --, --; ***T. frisea*** L., DLN 4799, South Africa, *R. Bayliss 8037* (BR), OP184185, --, --; ***T. fruticosum*** A.W.Hill, DLN 4115, South Africa, *D. Nickrent & E. Brink 4115* (BH), OP184186, --, --; DLN 4706, South Africa, *R. Bayliss 7694* (BR), OP184187, OP172294, OP172019; DLN 4798, South Africa, *R. Bayliss 7179* (BR), OP184188, OP172295, OP172020; ***T. fruticulosum*** A.W.Hill, DLN 4677, South Africa, *J. Leonard 7314* (BR), OP184189, OP172296, --; DLN 5504, South Africa, *M.A. García & al. 5504* (MA, BH, NBG), OP184190, OP172297, OP172021; ***T. funale*** L., DLN 4399, South Africa, *S. Williams 1307* (MO), OP184191, OP172298, OP172022; DLN 5506, South Africa, *L. Mucina 171205/14* (MA), OP184192, OP172299, OP172023; DLN 5512, South Africa, *L. Mucina 100906/18* (MA), OP184193, OP172300, OP172024; DLN 5518, South Africa, *L. Mucina 030406/04* (MA), OP184194, OP172301, OP172025; ***T. galioides*** A.DC., DLN 4400, South Africa, *K. Dahlstrand 2322* (MO), OP184195, --, --; DLN 4708, South Africa, *R. Bayliss 8393* (BR), OP184196, --, --; DLN 5521, South Africa, *L. Mucina 190206/4* (MA), OP184197, OP172302, OP172026; ***T. germainii*** Robyns & Lawalrée, DLN 4804, Democratic Republic of the Congo, *C. Lefebvre & al. 35* (BR), OP184198, OP172303, OP172027; ***T. gnidiaceum*** A.DC., DLN 4402, South Africa, *R. Bayliss 6901* (MO), OP184199, OP172304, OP172028; ***T. goetzeanum*** Engl., DLN 4386, Zimbabwe, *R. Bayliss 9020* (MO), OP184200, OP172305, --; DLN 4387, South Africa, *J. Pienaar 637* (MO), OP184201, OP172306, OP172029; DLN 4401, Zimbabwe, *E. Mshasha 103* (MO), OP184202, OP172307, OP172030; DLN 4753, South Africa, *H. Merxmüller 405* (BR), OP184203, OP172308, --; DLN 4808, Zimbabwe, *T. Baudesson & al. 62* (BR), OP184204, OP172309, OP172031; DLN 4810, Zambia, *B. Leteinturier & al. 317* (BR), OP184205, OP172310, OP172032; DLN 5250, Zambia, *D. Harder & M. Bingham 3511* (MO), OP184206, OP172311, OP172033; ***T. gontscharovii*** Bobrov, DLN 5108, Tajikistan, *R.V. Kamelin s.n.* (LE), OP184207, OP172312, OP172034; ***T. gracilarioides*** A.W.Hill, DLN 4420, South Africa, *S.D. Williamson 207* (MO), OP184208, OP172313, OP172035; DLN 5253, South Africa, *L. Dreyer 203* (MO), OP184209, --, --; ***T. gracile*** A.W.Hill, DLN 5280 clone A, South Africa, *B. Pienaar & P. Kok 1302* (MO), OP184210 (ITS clone A), OP184211 (ITS clone B), OP172314, OP172036; ***T. griseum*** Sond., DLN 5579, South Africa, *V.R. Clark & al. 113* (GRA), OP184212, OP172315, OP172037; ***T. gypsophiloides*** A.W.Hill, DLN 4711, South Africa, *P. Goetghebeur 4438* (BR), OP184213, OP172316, OP172038; DLN 5255, South Africa, *K. Balkwill & al. 10873* (MO), OP184214, OP172317, OP172039; DLN 5256, South Africa, *J. Brenan 14274* (MO), OP184215, OP172318, --; ***T. hararensis*** A.G.Mill., DLN 4814, Somalia, *M. Thulin & A. Warfa 6041* (UPS), OP184216, OP172319, OP172040; ***T. helichrysoides*** A.W.Hill, DLN 5439, South Africa, *M.A. García & al. 5439* (MA, BH, NBG), OP184217, OP172320, OP172041; ***T. himalense*** Royle ex Edgew., DLN 4815, Nepal, *S. Einarsson & al. 376* (UPS), OP184218, OP172321, OP172042; ***T. hispidulum*** Lam., DLN 5286, South Africa, *L. Hugo 643* (MO), OP184219, OP172322, --; DLN 5464, South Africa, *M.A. García & al. 5464* (MA, BH, NBG), OP184220, OP172323, OP172043; DLN 5479, South Africa, *M.A. García & al. 5479* (MA, BH, NBG), OP184221, --, --; DLN 5480, South Africa, *L. Mucina 5480* (MA, BH, NBG), OP184222, OP172324, OP172044; ***T. hockii*** Robyns & Lawalrée, DLN 4714, Democratic Republic of the Congo, *A. Schmitz 7797* (BR), OP184223, OP172325, OP172045; ***T. humifusum*** DC., DLN 4174, Spain, *M. Chase 1881* (K), OP184224, OP172326, --; DLN 4175, UK, M. Chase 3687 (K), OP184225, OP172327, --; DLN 5194, Spain, *M.A. García MAG-3917* (MA), OP184226, OP172328, OP172046; ***T. humile*** Vahl, DLN 4176, Europe, *M. Chase 1858* (K), OP184227, --, --; DLN 4841, Spain, *C. Navarro & al. CN 1879* (MA), OP184228, OP172329, OP172047; ***T. hystrix*** A.W.Hill, DLN 5257, South Africa, *C. Peeters & al. 260* (MO), OP184229, OP172330, --; DLN 5258, South Africa, *P. Burgoyne & N. Snow 4845* (MO), OP184230, OP172331, OP172048; ***T. imbricatum*** Thunb., DLN 5584, South Africa, *V.R. Clark & R. Cerros 386* (GRA), OP184231, OP172332, OP172049; DLN 5591, South Africa, *V.R. Clark & T.W. Naude 194* (GRA), OP184232, OP172333, OP172050; ***T. impeditum*** A.W.Hill, DLN 5583, South Africa, *V.R. Clark & R. Cerros 222* (GRA), OP184233, --, --; DLN 5587, South Africa, *V.R. Clark & al. 183* (GRA), OP184234, OP172334, OP172051; DLN 5589, South Africa, *V.R. Clark & I. Crause 224* (GRA), OP184235, OP172335, OP172052; DLN 5593, South Africa, *V.R. Clark & T.W. Naude 315* (GRA), OP184236, OP172336, OP172053; ***T. inversum*** N.E.Br., DLN 5259, South Africa, *K. Balkwill & al. 7069* (MO), OP184237, OP172337, --; ***T. junceum*** Bernh. ex Krauss, DLN 4716, South Africa, *R. Bayliss 7636* (BR), OP184238, --, --; DLN 5414, South Africa, *M.A. García & al. 5414* (MA, BH, NBG), OP184239, OP172338, OP172054; DLN 5588, South Africa, *V.R. Clark. & G. Coombs 95* (GRA), OP184240, OP172339, OP172055; ***T. juncifolium*** A.DC., DLN 5470, South Africa, *L. Mucina & G. López González 5470* (MA, BH, NBG), OP184241, OP172340, OP172056; DLN 5507, South Africa, *L. Mucina 171205/20* (MA), OP184242, OP172341, OP172057; ***T. kilimandscharicum*** Engl., DLN 4741, Tanzania, *J. Lovett & al. 3876* (BR), OP184243, OP172342, OP172058; DLN 4816, Tanzania, *R. Brummitt & P. Goldblatt 18091* (UPS), OP184244, OP172343, OP172059; ***T. kotschyanum*** Boiss., DLN 4404, Iran, *K. Rechinger 53599* (MO), OP184245, OP172344, OP172060; DLN 4405, Turkmenistan, *D. Kurbanov 1550* (MO), OP184246, OP172345, OP172061; DLN 5111, Turkmenistan, *A.A. Meseryakov s.n.* (LE), OP184247, OP172346, OP172062; ***T. lacinulatum*** A.W.Hill, DLN 5260, Namibia, *P. Burgoyne & N. Snow 3610* (MO), OP184248, OP172347, OP172063; ***T. leptocaule*** Sond., DLN 4719, South Africa, *R. Bayliss 6774* (BR), OP184249, OP172348, OP172064; DLN 4720, South Africa, *M. Wells 2700* (BR), OP184250, --, --; ***T. lesliei*** N.E.Br., DLN 5289, South Africa, *K. Balkwill & Reddy 7191* (MO), OP184251, OP172349, --; ***T. lewallei*** Lawalrée, DLN 4722, Burundi, *J. Lewalle 1326* (BR), OP184252, OP172350, OP172065; ***T. linophyllon*** L., DLN 4843, Italy, *C. Aedo & al. CA 7988* (MA), OP184253, OP172351, OP172066; DLN 5138, Ukraine, *V.M. Vinogradova & V.N. Gladkova s.n.* (LE), OP184254, OP172352, OP172067; ***T. lobelioides*** A.DC., DLN 4817, South Africa, *J.P.M. Brenen 14920* (UPS), OP184255, OP172353, OP172068; ***T. longiflorum*** Hand.-Mazz., DLN 4407, China, *D. Boufford & al. 28954* (MO), OP184256, OP172354, --; ***T. longifolium*** Turcz., DLN 4408, China, *Q.R. Wu 90-84* (MO), OP184257, --, --; DLN 5144, Russian Federation, *I.M. Krasnoborov 516* (LE), OP184258, OP172355, OP172069; ***T. lopollense*** Hiern, DLN 5264, Botswana, *P.A. Smith 2282* (MO), OP184259, --, --; ***T. luembense*** Robyns & Lawalrée, DLN 4806, Democratic Republic of the Congo, *S. Lisowski & al. 8027* (BR), OP184260, OP172356, OP172070; ***T. macranthum*** Fenzl, DLN 4842, Turkey, *S. Castroviejo & S. Nissa 15890SC* (MA), OP184261, OP172357, OP172071; ***T. macrostachyum*** A.DC., DLN 4409, South Africa, *P. Goldblatt & D. Snijman 6975* (MO), OP184262, --, --; ***T. madagascariense*** A.DC., DLN 4793, Madagascar, *C. Evrard 11284* (BR), OP184263, OP172358, OP172072; ***T. magalismontanum*** Sond., DLN 4728, South Africa, *G. Germishuizen 3355* (BR), OP184264, --, --; DLN 5267, South Africa, *G. Germishuizen 3355* (MO), OP184265, OP172359, OP172073; ***T. mauritanicum*** Batt., DLN 4844, Morocco, *C. Aedo & al. CA4296* (MA), KP318966, OP172360, OP172074; DLN 5193, Morocco, *S. Castroviejo & al. SC17994* (MA), KP318967, KP318976, OP172075; ***T. microcarpum*** A.DC., DLN 4410, South Africa, *M. Wells 2699* (MO), OP184266, OP172361, --; DLN 5389, South Africa, *M.A. García & al. 5389* (MA, BH, NBG), OP184267, --, --; DLN 5522, South Africa, *L. Mucina & G. Jakubowsky 190206/21* (MA), OP184268, --, OP172076; ***T. microcephalum*** A.W.Hill, DLN 5565, South Africa, *E. Pienaar M1054* (MA), OP184269, OP172362, OP172077; DLN 5570, South Africa, *E. Pienaar M378* (MA), OP184270, OP172363, OP172078; ***T. microphyllum*** Robyns & Lawalrée, DLN 4733, Burundi, *P. Auquier 4310* (BR), OP184271, OP172364, OP172079; ***T. minkwitzianum*** B.Fedtsch., DLN 5112, Uzbekistan, *R. Kamelin & al. 18* (LE), KP318964, KP318974, OP172080; ***T. minus*** (A.W.Hill) J.C.Manning & F.Forest, DLN 4411, South Africa, *S. Williams 355* (MO), OP184272, OP172365, --; ***T. moesiacum*** Velen., DLN 5139, Ukraine, *S.N. Miliutin s.n.* (LE), OP184273, OP172366, OP172081; ***T. montanum*** Ehrh. ex Hoffm., DLN 5160, Czech Republic, *R. Hendrych s.n.* (LE), OP184274, OP172367, OP172082; ***T. mukense*** A.W.Hill, DLN 4820, Kenya, *C. Fries & Th. Fries 313* (UPS), OP184275, OP172368, OP172083; ***T. multicaule*** Ledeb., DLN 5115, Kazakhstan, *G.I. Tscherkasova s.n.* (LE), OP184276, OP172369, OP172084; DLN 5117, Kazakhstan, *E.V. Karamysheva & al. 624* (LE), OP184277, OP172370, OP172085; DLN 5119, Kazakhstan, *R.V. Kamelin 1534* (LE), OP184278, --, --; ***T. multiramulosum*** Pilg., DLN 5284, South Africa, *L. Dreyer 251* (MO), OP184279, OP172371, --; ***T. myriocladum*** Baker ex A.W.Hill, DLN 4737, Zambia, *B. Simon & C. Williamson 1395* (BR), OP184280, --, --; ***T. namaquense*** Schltr., DLN 4738, South Africa, *W. Hanekom 2396* (BR), OP184281, OP172372, --; ***T. nautimontanum*** M.A. García, Nickrent & Mucina, DLN 5562, South Africa, *J. Šibík & I. Šibíkova MS463* (MA), MH807489, OP172373, OP172086; ***T. nigromontanum*** Sond., DLN 4415, South Africa, *M. Thompson 2118* (MO), OP184282, --, --; DLN 4742, South Africa, *R. Bayliss 6579* (BR), OP184283, --, --; DLN 5407, South Africa, *M.A. García & al. 5407* (MA, BH, NBG), OP184284, OP172374, OP172087; DLN 5421, South Africa, *M.A. García & al. 5421* (MA, BH, NBG), OP184285, OP172375, OP172088; ***T. nigrum*** A.W.Hill, DLN 4854, South Africa, *D. Morris 47* (NY), OP184286, OP172376, OP172089; ***T. nudicaule*** A.W.Hill, DLN 4098, South Africa, *D. Nickrent & al. 4098* (BH), OP184287, --, --; DLN 5373, South Africa, *M.A. García & al. 5373* (MA, BH, NBG), OP184288, OP172377, OP172090; ***T. oresigenum*** Compton, DLN 5575, South Africa, *E. Pienaar C31* (MA), OP184289, OP172378, OP172091; ***T. orientale*** A.W.Hill, DLN 5279, South Africa, *O. Hilliard & B. Burtt 13219* (MO), OP184290, OP172379, --; ***T. pallidum*** A.DC., DLN 4680, South Africa, *R. Bayliss 8244* (BR), OP184291, OP172380, OP172092; DLN 5221, South Africa, *A. Nicholas & L. Smook 2377* (MO), OP184292, OP172381, OP172093; ***T. panganense*** Polhill, DLN 4775, Tanzania, *E. Milne-Redhead & P. Taylor 7314* (BR), OP184293, OP172382, OP172094; ***T. paniculatum*** L., DLN 5282, South Africa, *E. Esterhuysen 35382a* (MO), OP184294, OP172383, --; ***T. parnassi*** A.DC., DLN 5161, Bosnia Herzegovina*, L. Adamovic s.n.* (LE), OP184295, OP172384, OP172095; DLN 5162, Italy, *G. Loreto s.n.* (LE), OP184296, --, --; ***T. passerinoides*** Robyns & Lawalrée, DLN 4744, Burundi, *D. Van de Ben T894* (BR), OP184297, OP172385, OP172096; DLN 4745, Burundi, *M. Reekmans 6566* (BR), OP184298, --, --; DLN 4746, Burundi, *A. Christiaensen 2495* (BR), OP184299, --, --; ***T. pawlowskianum*** Lawalrée, DLN 4747, Democratic Republic of the Congo, *F. Malaisse 9579* (BR), OP184300, OP172386, OP172097; ***T. phyllostachyum*** Sond., DLN 4437, South Africa, *P. Forbes 828* (MO), OP184301, OP172387, OP172098; ***T. pinifolium*** A.DC., DLN 4417, South Africa, *E. Esterhuysen 35475* (MO), OP184302, OP172388, --; ***T. polycephalum*** Schltr., DLN 5510, South Africa, *L. Mucina 030903/36* (MA), OP184303, OP172389, OP172099; ***T. polygaloides*** A.W.Hill, DLN 5252, South Africa, *S. Hobson 442* (MO), OP184304, OP172390, --; ***T. procumbens*** C.A.Mey., DLN 4846, Turkey, *J.J. Aldasoro & al. 6260* (MA), OP184305, --, --; DLN 5140, Ukraine, *N.N. Tzvelev s.n.* (LE), OP184306, --, --; DLN 5178, Russian Federation, A*.I. Leskov & A.P. Rusalev s.n.* (LE), OP184307, OP172391, --; DLN 5181, Russian Federation, *L.V. Averyanov & al. 3341* (LE), OP184308, OP172392, OP172100; ***T. prostratum*** A.W.Hill, DLN 5467, South Africa, *M.A. García & al. 5467* (MA, BH, NBG), OP184309, OP172393, OP172101; ***T. pseudocystoseiroides*** Cavaco & Keraudren, DLN 5285, Madagascar, *T. Croat 29041* (MO), OP184310, OP172394, --; ***T. pseudovirgatum*** Levyns, DLN 5406, South Africa, *M.A. García & al. 5406* (MA, BH, NBG), OP184311, OP172395, OP172102; DLN 5491, South Africa, *M.A. García & al. 5491* (MA, BH, NBG), OP184312, OP172396, OP172103; DLN 5519, South Africa, *L. Mucina & G. Jakubowsky 120206/01* (MA), OP184313, OP172397, OP172104; DLN 5576, South Africa, *E. Pienaar C163* (MA), OP184314, --, --; ***T. psilotoides*** Hance, DLN 4856 clone A, Philippines, *R. Williams 1310* (NY), OP184315 (ITS clone A), OP184316 (ITS clone B), OP172398, --; ***T. pubescens*** A.DC., DLN 5304, South Africa, *J. Lavranos 22972* (MO), OP184317, --, --; DLN 5485, South Africa, *M.A. García & al. 5485* (MA, BH, NBG), OP184318, OP172399, OP172105; ***T. pycnanthum*** Schltr., DLN 5375, South Africa, *M.A. García & al. 5375* (MA, BH, NBG), OP184319, OP172400, OP172106; DLN 5520, South Africa, *L. Mucina & G. Jakubowsky 120206/02* (MA), OP184320, OP172401, OP172107; ***T. pyrenaicum*** Pourr., DLN 5195, Spain, *P. Escobar & al. 492/04* (MA), OP184321, OP172402, OP172108; DLN 5196, Spain, *M.A. García & al. MAG-3915* (MA), OP184322, OP172403, OP172109; ***T. quinqueflorum*** Sond., DLN 5447, South Africa, *M.A. García & al. 5447* (MA, BH, NBG), OP184323, OP172404, OP172110; DLN 5495, South Africa, *M.A. García & al. 5495* (MA, BH, NBG), OP184324, OP172405, OP172111; ***T. radicans*** Hochst. ex A.DC., DLN 4822, Ethiopia, *M. Gilbert. & al. 8275* (UPS), OP184325, OP172406, OP172112; ***T. ramosissimum*** Bobrov, DLN 5120, Tajikistan, *R.V. Kamelin s.n.* (LE), OP184326, OP172407, OP172113; ***T. ramosoides*** Hendrych, DLN 5170, China, *E. Maire s.n.* (LE), OP184327, OP172408, --; ***T. ramosum*** Hayne, DLN 4419, Georgia, *R. Gagnidze & al. 1011* (MO), OP184328, --, --; DLN 5105, Kazakhstan, *V.I. Vasilievich & al. 2046* (LE), OP184329, --, OP172114; DLN 5132, Ukraine, *B.N. Klopotov s.n.* (LE), OP184330, --, --; DLN 5134, Ukraine, *S.V. Yuzepchuk 1378* (LE), OP184331, OP172409, OP172115; DLN 5182, Russian Federation, *Yu.L. Menitsky & al. 122* (LE), OP184332, OP172410, OP172116; DLN 5558, United States, *M. Lavin s.n.* (BH), OP184333, --, --; ***T. reekmansii*** Lawalrée, DLN 4752, Burundi, *M. Reekmans 2957* (BR), OP184334, --, --; ***T. refractum*** C.A.Mey., DLN 4422, Russian Federation, *I. Krasnoborov & M. Danilov 129* (MO), OP184335, --, --; DLN 4423, Russian Federation, *I. Krasnoborov 35* (MO), OP184336, --, --; DLN 5145, Russian Federation, *I. Pshenichnaya & G. Liventsova s.n.* (LE), OP184337, OP172411, OP172117; DLN 5148, Russian Federation, *E. Korotkova & M. Danilov 812* (LE), OP184338, OP172412, OP172118; DLN 5155, Russian Federation, *G.A. Abrosimova & A.E. Matsenko 433* (LE), OP184339, OP172413, OP172119; ***T. repens*** Ledeb., DLN 5167, Mongolia, *A.L. Budantsev & al. 230* (LE), OP184340, OP172414, OP172120; ***T. resedoides*** A.W.Hill, DLN 4424, South Africa, *H.F. Glen 2480* (MO), OP184341, OP172415, --; DLN 4754, South Africa, *P. Goetghebeur 4390* (BR), OP184342, OP172416, OP172121; DLN 4823, Botswana, *O. Hansen 3456* (UPS), OP184343, OP172417, OP172122; ***T. retamoides*** (A.Santos) J.C.Manning & F.Forest, DLN 5559, Spain: Canary Islands, *E. Hernández s.n.* (MA), KP318961 (ITS clone A), OP184344 (ITS clone B), KP318971, OP172123; ***T. rostratum*** Mert. & W.D.J.Koch, DLN 4847, Italy, *L. Poldini s.n.* (MA), OP184345, OP172418, OP172124; DLN 5163, Italy, *A. Ylimartimo s.n.* (LE), OP184346, --, --; ***T. rufescens*** A.W.Hill, DLN 5517, South Africa, *L. Mucina 030406/19* (MA), OP184347, OP172419, OP172125; ***T. saxatile*** Turcz. ex A.DC., DLN 4425, Russian Federation, *A. Skvortsov s.n.* (MO), OP184348, OP172420, OP172126; DLN 5152, Russian Federation, *Mijno s.n.* (LE), OP184349, OP172421, OP172127; ***T. scabrum*** L., DLN 4426, South Africa, *K. Steiner 2256* (MO), OP184350, OP172422, OP172128; DLN 5492, South Africa, *M.A. García & G. López González 5492* (MA, BH, NBG), OP184351, --, --; DLN 5503, South Africa, *M.A. García & al. 5503* (MA, BH, NBG), OP184352, OP172423, OP172129; ***T. scandens*** Sond., DLN 4857, South Africa, *R. Bayliss 1423* (NY), OP184353, --, --; DLN 5399, South Africa, *M.A. García & al. 5399* (MA, BH, NBG), OP184354, OP172424, OP172130; ***T. schaijesii*** Lawalrée, DLN 4755, Democratic Republic of the Congo, *M. Scheijes 2735* (BR), OP184355, OP172425, OP172131; ***T. schliebenii*** Pilg., DLN 4756, Zambia, *H. Richards 5214* (BR), OP184356, OP172426, --; DLN 4805, Democratic Republic of the Congo, M. *Schaijes 1806* (BR), OP184357, OP172427, OP172132; ***T. schmitzii*** Robyns & Lawalrée, DLN 4757, Democratic Republic of the Congo, *F. Malaisse 13645* (BR), OP184358, --, --; DLN 4758, Burundi, *P. Auquier 4337* (BR), OP184359, OP172428, OP172133; ***T. schweinfurthii*** Engl., DLN 4759, Tanzania, *R. Gereau & al. 4976* (BR), OP184360, OP172429, OP172134; DLN 4797, Uganda, *P. Rwaburindore 3639* (BR), OP184361, --, --; DLN 4824, Ethiopia, *M. Gilbert & M. Thulin 619* (UPS), OP184362, OP172430, OP172135; ***T. scirpioides*** A.W.Hill, DLN 5233, Mozambique, *D. Zunguze & C. Boane 96* (MO), OP184363, OP172431, --; DLN 5292, South Africa, *O. Hilliard & B. Burtt 18535* (MO), OP184364, OP172432, --; ***T. selagineum*** A.DC., DLN 5378, South Africa, *M.A. García & al. 5378* (MA, BH, NBG), OP184365, OP172433, OP172136; ***T. sertulariastrum*** A.W.Hill, DLN 5425, South Africa, *M.A. García & al. 5425* (MA, BH, NBG), OP184366, OP172434, OP172137; DLN 5511, South Africa, *L. Mucina 100906/16* (MA), OP184367, OP172435, OP172138; ***T. setulosum*** Robyns & Lawalrée, DLN 4760, Zambia, *A. Angus 398* (BR), OP184368, --, --; ***T. simplex*** Velen., DLN 5141, Moldova, *A. Arvat s.n.* (LE), OP184369, OP172436, --; ***T. sonderianum*** Schltr., DLN 5392, South Africa, *M.A. García & al. 5392* (MA, BH, NBG), OP184469, OP172517, OP172213; DLN 5394, South Africa, *M.A. García & al. 5394* (MA, BH, NBG), OP184370, OP172437, OP172139; ***Thesium sp.***, DLN 4393, South Africa, *E. Cloete s.n.* (MO), OP184371, OP172438, --; ***Thesium sp.***, DLN 4686, South Africa, *P. Goetghebeur 4332* (BR), OP184372, OP172439, OP172140; ***Thesium sp.***, DLN 4709, South Africa, *G. Germishuizen 3389* (BR), OP184373, OP172440, OP172141; ***Thesium sp.***, DLN 4710, Zimbabwe, *C. Vanden Berghen 91* (BR), OP184374, OP172441, OP172142; ***Thesium sp.***, DLN 4796, Burkina Faso, *J. Lejoly 82/382* (BR), OP184375, OP172442, OP172143; ***Thesium sp.***, DLN 4807, Zimbabwe, *T. Baudesson & al. 471* (BR), OP184376, OP172443, OP172144; ***Thesium sp.***, DLN 4818, Leshoto, *O. Hedberg 82120* (UPS), OP184377, OP172444, OP172145; ***Thesium sp.***, DLN 4819, Ethiopia, *M. Thulin & A. Hunde 3975* (UPS), OP184378, OP172445, OP172146; ***Thesium sp.***, DLN 4825, Ethiopia, *M. Gilbert & al. 8287* (UPS), OP184379, OP172446, OP172147; ***Thesium sp.***, DLN 5237, South Africa, *K. Balkwill 7868* (MO), OP184380, OP172447, OP172148; ***Thesium sp.***, DLN 5263, China, *Y. Shu-Xiu 160* (MO), OP184381, OP172448, --; ***Thesium sp.***, DLN 5300, South Africa, *C. Bredenkamp 717* (MO), OP184382, OP172449, OP172149; ***Thesium sp.***, DLN 5308, South Africa, *K. Balkwill & Balkwill 4555* (MO), OP184383 (ITS clone A), OP184384 (ITS clone B), OP172450, --; ***Thesium sp.***, DLN 5309, South Africa, *E. Oliver 10324* (MO), OP184385, OP172451, OP172150; ***Thesium sp.***, DLN 5377, South Africa, *M.A. García & al. 5377* (MA, BH, NBG), OP184386, OP172452, OP172151; ***Thesium sp.***, DLN 5393, South Africa, *M.A. García & al. 5393* (MA, BH, NBG), OP184387 (ITS clone A), OP184388 (ITS clone B), OP184389 (ITS clone C), OP172453, OP172152; ***Thesium sp.***, DLN 5566, South Africa, *E. Pienaar M336* (MA), OP184393, OP172457, OP172156; ***Thesium sp.***, DLN 5567, South Africa, *E. Pienaar T65* (MA), OP184394, OP172458, --; ***Thesium sp.***, DLN 5574, South Africa, *E. Pienaar M1184* (MA), OP184395, OP172459, OP172157; ***Thesium sp.* sect. *Frisea***, DLN 5384, South Africa, *M.A. García & al. 5384* (MA, BH, NBG), OP184396, OP172460, OP172158; ***Thesium sp.* sect. *Frisea***, DLN 5416, South Africa, *M.A. García & al. 5416* (MA, BH, NBG), OP184397, OP172461, OP172159; ***Thesium sp.* sect. *Frisea***, DLN 5422, South Africa, *M.A. García & al. 5422* (MA, BH, NBG), OP184398, --, --; ***Thesium sp.* sect. *Frisea***, DLN 5426, South Africa, *M.A. García & al. 5426* (MA, BH, NBG), OP184399, OP172462, OP172160; ***Thesium sp.* sect. *Frisea***, DLN 5431, South Africa, *M.A. García & al. 5431* (MA, BH, NBG), OP184400, OP172463, OP172161; ***Thesium sp.* sect. *Frisea***, DLN 5437, South Africa, *M.A. García & al. 5437* (MA, BH, NBG), OP184401, OP172464, OP172162; ***Thesium sp.* sect. *Frisea***, DLN 5453, South Africa, *M.A. García & al. 5453* (MA, BH, NBG), OP184402, OP172465, OP172163; ***Thesium sp.* sect. *Frisea***, DLN 5460, South Africa, *M.A. García & al. 5460* (MA, BH, NBG), OP184403, OP172466, OP172164; ***Thesium sp.* sect. *Frisea***, DLN 5482, South Africa, *M.A. García & al. 5482* (MA, BH, NBG), OP184404, OP172467, OP172165; ***Thesium sp.* sect. *Frisea***, DLN 5494, South Africa, *M.A. García & al. 5494* (MA, BH, NBG), OP184405, --, --; ***Thesium sp.* sect. *Frisea***, DLN 5523, South Africa, *L. Mucina 190206/3* (MA), OP184406, OP172468, OP172166; ***Thesium sp.* sect. *Frisea***, DLN 5577, South Africa, *E. Pienaar M87* (MA), OP184407, --, --; ***T. spicatum*** L., DLN 4095, South Africa, *D. Nickrent & al. 4095* (BH), OP184408, --, --; DLN 4761, South Africa, *R. Bayliss 7748* (BR), OP184409, --, --; ***T. spinosum*** L.f., DLN 4762, South Africa, *E. Coppejans 1476* (BR), OP184410, OP172469, OP172167; DLN 5513, South Africa, *L. Mucina 090306/07* (MA), OP184411, OP172470, OP172168; ***T. spinulosum*** A.DC., DLN 5295, South Africa, *E. Esterhuysen 34460* (MO), OP184412, --, --; DLN 5424, South Africa, *M.A. García & al. 5424* (MA, BH, NBG), OP184413, KP318978, OP172169; DLN 5498, South Africa, *M.A. García & al. 5498* (MA, BH, NBG), OP184414, OP172471, OP172170; ***T. stelleroides*** Jaub. & Spach, DLN 4848, Turkey, *F. Muñoz Garmedia & al. 4525* (MA), KP318963, KP318973, OP172171; ***T. strictum*** P.J.Bergius, DLN 4092, South Africa, *D. Nickrent & al. 4092* (BH), OP184415, --, - -; DLN 4763, South Africa, *P. Bohnen 9003* (BR), OP184416, OP172472, OP172172; DLN 4764, South Africa, *E. Coppejans 459* (BR), OP184417, OP172473, OP172173; DLN 4811, South Africa, *H. Breyne 5414* (BR), OP184418, OP172474, OP172174; DLN 5505, South Africa, *L. Mucina 021106/7* (MA), OP184419, OP172475, OP172175; DLN 5508, South Africa, *L. Mucina 171205/02* (MA), OP184420, OP172476, OP172176; DLN 5515, South Africa, *L. Mucina 090206/27* (MA), OP184421, OP172477, OP172177; DLN 5516, South Africa, *L. Mucina 090206/10* (MA), OP184422, OP172478, OP172178; DLN 5526, South Africa, *L. Mucina 211006/51* (MA), OP184423, OP172479, OP172179; DLN 5568, South Africa, *E. Pienaar M646* (MA), OP184424, OP172480, OP172180; ***T. stuhlmannii*** Engl., DLN 4765, Tanzania, *R. Gereau & C. Kayombo 4664* (BR), OP184425, OP172481, OP172181; DLN 4766, Tanzania, *J. Lovett & C. Kayombo 3402* (BR), OP184426, OP172482, OP172182; ***T. subaphyllum*** Engl., DLN 4767, Tanzania, *R. Gereau & C. Kayombo 3899* (BR), OP184427, OP172483, OP172183; ***T. subsimile*** N.E.Br., DLN 5254, South Africa, *K. Balkwill & M-J. Balkwill 11245* (MO), OP184428, OP172484, OP172184; ***T. subsucculentum*** (Kämmer) J.C.Manning & F.Forest, DLN 4374, Spain: Canary Islands, *A. Santos Guerra s.n.* (MA), KP318962 (ITS clone A), OP184429 (ITS clone B), OP184430 (ITS clone C), KP318972, OP172185; ***T. symoensii*** Lawalrée, DLN 4768, Democratic Republic of the Congo, *S. Lisowski & al. 4276* (BR), OP184431, OP172485, --; ***T. szowitsii*** A.DC., DLN 4434, Iran, *K. Rechinger 41333* (MO), OP184432, OP172486, OP172186; DLN 5191, Armenia, *C. Navarro & al. CN-5602* (MA), OP184433, OP172487, OP172187; ***T. tamariscinum*** A.W.Hill, DLN 4769, Tanzania, *J. Lovett & C. Kayombo 3401* (BR), OP184434, OP172488, OP172188; ***T. tenuifolium*** Saut. ex W.D.J.Koch, DLN 5142, Russian Federation, *N.N. Tzvelev 95* (LE), OP184435, OP172489, OP172189; ***T. tenuissimum*** Hook. f., DLN 4772, Cameroon, *W. Meijer 15407* (BR), OP184436, OP172490, OP172190; ***T. tepuiense*** Steyerm., DLN 4858, Guyana, *B. Maguire & D. Fanshawe 32543* (NY), OP184437, OP172491, --; ***T. thamnus*** Robyns & Lawalrée, DLN 4773, Democratic Republic of the Congo, *A. Schmitz 8124* (BR), OP184438, OP172492, OP172191; DLN 4792, Democratic Republic of the Congo, *M. Schaijes 2025* (BR), OP184439, OP172493, OP172192; ***T. translucens*** A.W.Hill, DLN 5500, South Africa, *M.A. García & al. 5500* (MA, BH, NBG), OP184440, OP172494, OP172193; ***T. triflorum*** L.f., DLN 4438, Tanzania, *J.A. Mlangwa & W. Kindeketta 717* (MO), OP184441, OP172495, OP172194; DLN 4679, South Africa, *R. Bayliss 7876* (BR), OP184442, --, --; DLN 4774, Tanzania, *D. Goyder & al. 3850* (BR), OP184443, OP172496, OP172195; DLN 5291, South Africa, *P. Phillipson & T. Dold 4188* (MO), OP184444, --, --; ***T. tuvense*** I.M.Krasnoborov, DLN 5172, Mongolia, *R.V. Kamelin & al. 1476* (LE), OP184445, OP172497, OP172196; ***T. unyikense*** Engl., DLN 4392, Malawi, *I.F. LaCroix 4756* (MO), OP184446, OP172498, OP172197; DLN 4776, Tanzania, *R. Brummit & al. 18192* (BR), OP184447, OP172499, OP172198; ***T. ussanguense*** Engl., DLN 4777, Zimbabwe, *P. Bamps & al. 691* (BR), OP184448, OP172500, --; DLN 4778, Malawi, *P. Tyrer 821* (BR), OP184449, OP172501, OP172199; ***T. utile*** A.W.Hill, DLN 4689, South Africa, *G. Germishuizen 3380* (BR), OP184450, OP172502, OP172200; DLN 5298, South Africa, *N. Kroon 11710* (MO), OP184451, OP172503, OP172201; ***T. vahrmeijeri*** Brenan, DLN 4779, Mozambique, *J. Koning 7869* (BR), OP184452, --, --; ***T. vimineum*** Robyns & Lawalrée, DLN 4783, Democratic Republic of the Congo, *S. Lisowski & al. 7720* (BR), OP184453, OP172504, OP172202; ***T. virens*** E.Mey. ex A.DC., DLN 5299, South Africa, *O. Hilliard & B. Burtt 9061* (MO), OP184454, OP172505, OP172203; ***T. virgatum*** Lam., DLN 5293, South Africa, *P. Goldblatt 3737* (MO), OP184455, --, --; DLN 5448, South Africa, *M.A. García & al. 5448* (MA, BH, NBG), OP184456, OP172506, OP172204; DLN 5481, South Africa, *M.A. García & al. 5481* (MA, BH, NBG), OP184457, OP172507, OP172205; DLN 5594, South Africa, *L. Mucina 130808/07* (MA), OP184458, OP172508, --; ***T. viride*** A.W.Hill, DLN 4770, Cameroon, *A. Jacques-Georges 27654* (BR), OP184459, OP172509, OP172206; DLN 4784, Togo, *J. Brunel 708* (BR), OP184460, OP172510, --; DLN 4785, Nigeria, *R. Meikle 1198* (BR), OP184461, OP172511, OP172207; ***T. viridifolium*** Levyns, DLN 4388, South Africa, *E.R. Orchard 571* (MO), OP184462, --, --; DLN 5442, South Africa, *M.A. García & al. 5442* (MA, BH, NBG), OP184463, OP172512, OP172208--; DLN 5451, South Africa, *M.A. García & al. 5451* (MA, BH, NBG), OP184464, --, --; DLN 5456, South Africa, *M.A. García & al. 5456* (MA, BH, NBG), OP184465, OP172513, OP172209; ***T. whyteanum*** Rendle, DLN 4787, Malawi, *J. Chapman & I. Patel 5779* (BR), OP184466, OP172514, OP172210; ***T. wilczekianum*** Lawalrée, DLN 4789, Democratic Republic of the Congo, F. *Malaisse 8407* (BR), OP184467, OP172515, OP172211; ***T. xerophyticum*** A.W.Hill, DLN 5302, Namibia, *H. Merxmüller & J. Giess 32566* (MO), OP184468, OP172516, OP172212. ibutions

## AUTHOR CONTRIBUTIONS

MAG and DLN designed the study and performed the sampling. MAG conducted DNA extractions and sequencing, performed phylogenetic analyses, submitted the sequences to NCBI GenBank, wrote the bulk of the manuscript, and constructed all phylogenetic trees. DLN helped with lab work and phylogenetic analyses, created figures 1 to 6 and assisted in writing and editing the manuscript. LM organized one of the collecting trips to South Africa, procured additional *Thesium* samples, and contributed in the writing and editing of the manuscript. All authors read and commented on the manuscript.

## Supporting Information

**Appendix S1**. Excel file showing sequencing details across the 396 *Thesium* accessions used in this study.

**Appendix S2**. Alignment of the nuclear ITS dataset.

**Appendix S3** Alignment of two concatenated plastid regions (*trnLF*, *trnDT*). Characters 1--1408 correspond to *trnLF* and 1409--3336 to *trnDT*.

**Appendix S4**. List of accepted names and synonyms for *Thesium* species as well as their general geographic distribution.

**Appendix S5**. List of authors and Creative Commons (CC) licenses for photographs used in Figures 3--6.

## Photograph Credits and Permissions

### Abbreviations

CC: Creative Commons
MAG: M.A. García
GLG: G. López González
DLN: D.L. Nickrent
Phytoimages: noncommercial use allowed (see Terms of Use http://phytoimages.siu.edu/).

3A. *Lacomucinaea* clade, *L. lineata*. Gerhard Glatzel, Phytoimages.

3B. *Kunkeliella* clade, *T. subsucculentum*. Mario Mairal (habit), iNaturalist (CC BY-NC-SA 4.0); Biodiversity Data Bank of the Canary Islands: R. Mesa-Coello (flowers); L. Rodríguez Navarro (fruit). Rules of use: https://www.biodiversidadcanarias.es/biota/normas?lang=en.

3C. *Hagnothesium* clade, *T. fragile*. MAG & GLG (habit and fruit); Nicola van Berkel (male flower), iNaturalist (CC BY 4.0).

3D. *Mauritanica* clade, *T. mauritanicum*. Abdelmonaim Homrani Bakali. Plant Biodiversity of South-Western Morocco (CC BY-NC 4.0).

3E. *Humilia* clade, *T. humile*. Felix Riegel (habit), iNaturalist (CC BY-NC 4.0); Tatyana Malchinski (fruits), Plantarium web site (https://www.plantarium.ru/). Message sent via email and permission given 21 Oct. 2022.

3F. *Macranthia* clade, *T. szowitsii*. MAG.

3G. Eurasian clade, *T. minkwitzianum.* Alexander Ebel (habit), Plantarium web site. Email message granting permission for use received 9 March 2022; DLN (insets), rehydrated herbarium specimens: *Kamelin & al. 18* (flower) and *Lazkov s.n.* (fruit), both LE.

3H. *Procumbens* clade, *T. procumbens*. MAG (habit and fruits); Gennadiy Okatov (flowers), iNaturalist (CC BY-NC 4.0).

3I. *Parnassi* clade, *T. parnassi*. Gianluca Nicolella. Acta plantarum, Galleria della Flora italiana (CC BY-NC-ND 4.0). https://www.actaplantarum.org/flora/flora_info.php?id=7747

3J. *Bavarum* clade, *T. bavarum*. Amadej Trnkoczy (shoots and fruits), Encyclopedia of Life (CC BY-NC-SA 3.0); Giuliano Da Zanche (flower), Flickr. Permission for use granted via Flickr and email message 3 Feb. 2023.

3K. *Alpina* clade, *T. alpinum*. Felix Riegel (shoot), iNaturalist (CC BY-NC 4.0); Hecklemore (flower), iNaturalist (CC BY-NC 4.0); Jiří Kameníček (fruits), BioLib.cz. Email message granting permission received 22 Oct. 2022.

3L. *Linophylla* clade, *T. humifusum*. Habit: Liliane Roubaudi (habit), Wikimedia Commons (CC BY-SA 2.0 FR); John Crossley (flowers), iNaturalist (CC BY-NC 4.0); Francisco Rodriguez - Faluke (fruit), iNaturalist (CC BY-NC 4.0).

3M. Second Asian Radiation, *T. catalaunicum*. Images provided by Joan Pedrol. 3N. *Rostratum* clade, *T. rostratum*. Andreas Fleischmann, Phytoimages.

3O. *Multicaule* clade, *T. ebracteatum.* Andrey Belekhov (shoot), Plantarium web site; message via Plantarium granting permission for use received 21 Oct. 2022. Jiří Kameníček (flower and fruit), BioLib.cz web site (https://www.biolib.cz/en/main/). Email message granting permission received 3 Feb. 2023.

3P. *Australe* clade, *T. chinense.* Vera Volkotrub (shoot), Plantarium web site; message via Plantarium granting permission for use received 23 Oct. 2022. Onidiras (flowers), iNaturalist (CC BY-NC 4.0); Fruit: from Suetsugu (2015), permission for use of Fig. 1b from Wiley (lic. no. 5480830039865) received 2 February 2023.

3Q. *Longifolia* clade, *T. refractum.* All from Plantarium web site: Elena Zhurba (habit), permission received 21 Oct. 2022. Photographer unknown (flower, from Primorsky Krai, Russia); Svetlana Nesterova (fruit), permission received 22 Oct. 2022.

3R. *Ramosum* clade, *T. ramosum*. MAG (shoot); DLN (flower and fruit).

3S. *Himalensia* clade, *T. himalense.* D.S. Rawat (shoot, flower), eFlora of India, via email granting permission for use received 22 Oct. 2022.

3T. *Alatavicum* clade, *T. alatavicum.* Galina Vladimirovna Chulanova, Plantarium web site. Received email granting permission for use received 7 February 2023.

4A. African clade, *T. nautimontanum*. DLN.

4B. *Spinosa* clade, *T. spinulosum*. MAG & GLG (habit); DLN (flower and fruit). 4C. *Namaquense* clade, *T. lacinulatum*. David Hoare (habit and flowers), iNaturalist (CC BY-NC 4.0); D. Wesuls (fruits), Useful Tropical Plants & Photo Guide to Plants of Southern African Plants (www.southernafricanplants.net). Site says: “Free use of the information and of the photographs is granted for non-commercial and educational purposes”.

4D. *Triflorum* clade, *T. scandens*. DLN.

4E. *Triflorum* clade, *T. triflorum.* Shaun Swanepoel (habit); Adriaan Grobler (flower and fruit); both iNaturalist (CC BY-NC 4.0).

4F. Core Cape clade, *T. euphorbioides*. DLN.

4G. *Foliosum* clade, *T. foliosum*. MAG & GLG (shoot); DLN (flower and fruit). 4H. *Ericaefolium* clade, *T. ericaefolium*. MAG & GLG (habit); DLN (flowers and fruits).

4I. *Scirpioides* clade, *T. flexuosum*. Habit: MAG & GLG (shoots); DLN (flower, fruit) 4J. *Strictum* clade, *T. albomontanum*. MAG & GLG (habit, fruits); DLN (flowers).

4K. *Capitatum* clade, *T. carinatum*. MAG & GLG (shoot with young fruits); DLN (flower, fruit).

4L. *Frisea* clade, *T. funale*. MAG & GLG (shoots); DLN (flowers and fruits).

4M. *Nigromontanum* clade, *T. nigromontanum*. MAG & GLG (shoots); DLN (flowers and fruits).

4N. *Virgatum* clade, *T. pseudovirgatum*. MAG & GLG (habit); DLN (flowers and fruits).

4O. *Capitellatum* clade, *T. prostratum*. MAG & GLG (shoots); DLN (flowers and fruits).

4P. *Acuminatum* clade, *T. capituliflorum*. MAG & GLG (shoots); DLN (flowers and fruits).

4Q. *Hispidulum* clade, *T. hispidulum*. MAG & GLG (shoots); DLN (flowers and fruits). 4R. *Commutatum* clade, *T. commutatum*. MAG & GLG (shoots); DLN (flowers and fruits).

4S. *Gnidaceum* clade, *T. gnidiaceum* (left); *T. impeditum* (right). Craig Peter (*T. gnidiaceum*, left), iNaturalist (CC BY-NC 4.0); SAplants (*T. impeditum*, right), Wikimedia Commons (CC BY-NC 4.0).

4T. *Gnidaceum* clade, *T. oresigenum* (left); *T. phylostachyum* (right). Nick Helme (*T. oresigenum*, left), iNaturalist (CC BY-NC 4.0); Nicola van Berkel (*T. phylostachyum* right), iNaturalist (CC BY 4.0).

5A. *Ussanguense* clade, *T. passerinoides*. DLN from herbarium specimen (*A.R. Christiaensen 249*5, BR).

5B. *Stuhlmannii* clade, *T. fimbriatum*. DLN from herbarium specimen (*P. Goldblatt 8056*, MO).

5C. *Austroamericium* clade, *T. aphyllum.* C. Luís A. Funez, Flora de Santa Catarina Sent, email message granting permission received 21Oct. 2022.

5D. *Tenuissimum* clade, *T. tenuissimum*. DLN from herbarium specimen (*W. Meijer 15407*, BR).

5E. *Reekmansii* clade, *T. madagascariense*. DLN from herbarium specimen (*C. Evrard 11284*, BR).

5F. *Reekmansii* clade, *T. wilczekianum*. DLN from herbarium specimen (*E. Milne- Redhead 3581*, BR).

5G. *Filipes* clade, *T. filipes*. *Ehoarn Bidault 2680* (MO), Tropicos Web Site (https://www.tropicos.org/name/28500430) (CC BY-NC-ND 3.0).

5H. *Amicorum* clade, *T. amicorum*. DLN from herbarium specimen (*St. Lisowski et al. 5736*, BR).

5I. *Viride* clade, *T. equisetoides. Ehoarn Bidault 2924* (MO), Tropicos (https://www.tropicos.org/name/28500426) (CC BY-NC-ND 3.0). Email message granting permission received 21 Oct. 2022.

5J. *Viride* clade, *T. fastigiatum*. *Cyrille Mas 1222* (MO), Tropicos (https://www.tropicos.org/name/28500428) (CC BY-NC-ND 3.0). Email message granting permission received 21 Oct. 2022.

5K. *Angulosum* clade, *T. angulosum*. Alison Young (shoots) iNaturalist (CC BY-NC 4.0); DLN (flowers and fruit L.S.).

5L. *Cupressoides* clade, *T. cupressoides*. DLN (habit); M. M. le Roux (flowers), Phytoimages; M.R. Popp (fruits), iNaturalist (CC0 1.0).

5M. LDD2 clade, *T. radicans*. *S. Collenette 8173* (E). Via GBIF, Royal Botanic Garden Edinburgh (CC0 1.0), https://www.gbif.org/occurrence/574687216.

5N. LDD2 clade, *T. psilotoides*. DLN from herbarium specimen (*R.S. Williams 1310*, NY).

5O. LDD2 clade, *T. decaryanum*. Guy Eric Onjalalaina, iNaturalist (CC BY-NC 4.0). 5P. *Pallidum* clade, *T. pallidum*. SAPlants Wikimedia Commons (CC BY-SA 4.0); M.M. le Roux (fruit), Phytoimages.

5Q. *Cornigerum* clade, *T. cornigerum*. Nick Helme, iNaturalist (CC BY-NC 4.0). 5R. *Kilimandscharicum* clade, *T. dolichomeras*. Nick Helme (habit and flowers), iNaturalist (CC BY-NC 4.0); B.T. Wursten (fruits), Flora of Mozambique (https://www.mozambiqueflora.com/speciesdata/image-display.php?species_id=121390&image_id=5). Email message granting permission received 22 Oct. 2022.

5S. *Resedoides* clade, *T. resedoides*. M.M. le Roux, Phytoimages.

5T. *Gracile* clade, *T. gracile*. Mervyn Lotter (habit), iNaturalist (CC BY-NC 4.0); M.M. le Roux (flowers and fruit), Phytoimages.

6A. *Alatum* clade, *T. alatum*. Sandra Falanga (shoots and flowers), iNaturalist (CC BY-NC 4.0); DLN from herbarium specimen (*P. Goetghebeur 4512*, BR) (fruits).

6B. *Alatum* clade, *T. utile*. Andrew Hankey (habit), iNaturalist (CC BY-SA 4.0); Nicola van Berkel (flower), iNaturalist (CC BY 4.0); DLN (fruit).

6C. *Davidsoniae* clade, *T. davidsoniae*. M.M. le Roux, Phytoimages.

6D. *Multiramulosum* clade, *T. multiramulosum*. Troos van der Merwe, iNaturalist (CC BY-SA 4.0).

6E. *Virens* clade, *T. subsimile*. MAG (habit and flowers L.S.) & DLN (fruits) from herbarium specimen (*K. Balkwill & M.-J. Balkwill 11245*, MO).

6F. *Nigrum* clade, *T. magalismontanum*. M.M. le Roux (shoots, flowers), Phytoimages; Tjeerd de Wit (inflorescence), iNaturalist (CC BY-SA 4.0); Message via Facebook granting permission received 22 Oct. 2022); DLN (fruit L.S).

6G. *Lobelioides* clade, *T. lobelioides*. MAG & DLN from herbarium specimen (*J.P.M. Brenen 14920*, UPS).

6H. *Goetzeanum* clade, *T. goetzeanum*. Richard Gill (habit), iNaturalist (CC BY-NC 4.0); M.M. le Roux (flowers), Phytoimages.

6I. *Gracilarioides* clade, *T. gracilarioides*. M. M. le Roux, Phytoimages.

6J. *Gracilarioides* clade, *T. polygaloides*. MAG & DLN from herbarium specimen (*Hobson 442*, MO).

6K. *Costatum* clade, *T. asterias*. magdastlucia (KwaZulu-Natal, South Africa), iNaturalist (CC BY-NC 4.0); M.M. le Roux (flower L.S.), Phytoimages.

6L. *Costatum* clade, *T. costatum*. M. M. le Roux, Phytoimages.

---

**Fig. S1.**
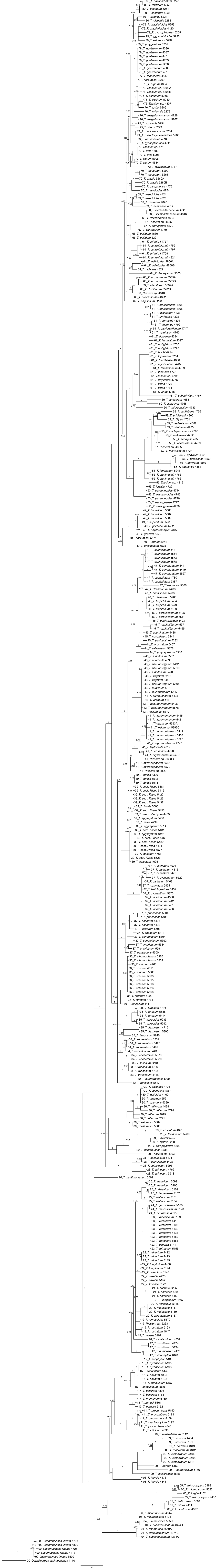
Phylogram resulting from the ITS dataset analyzed with Bayesian Inference. Numbers at nodes indicate posterior probabilities.

**Fig. S2.**
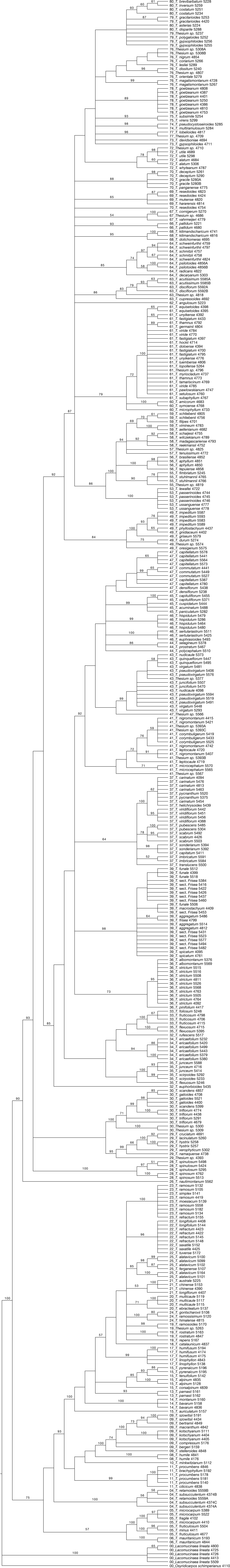
Cladogram resulting from the ITS dataset analyzed with Maximum Parsimony. Numbers at nodes indicate bootstrap values (1000 replicates).

**Fig. S3.**
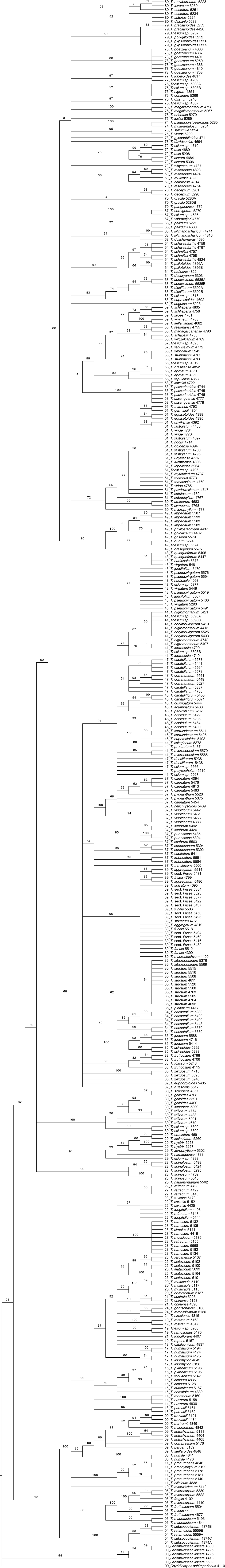
Cladogram resulting from the ITS dataset analyzed with Maximum Likelihood (GARLI). Numbers at nodes indicate bootstrap values (250 replications).

**Fig. S4.**
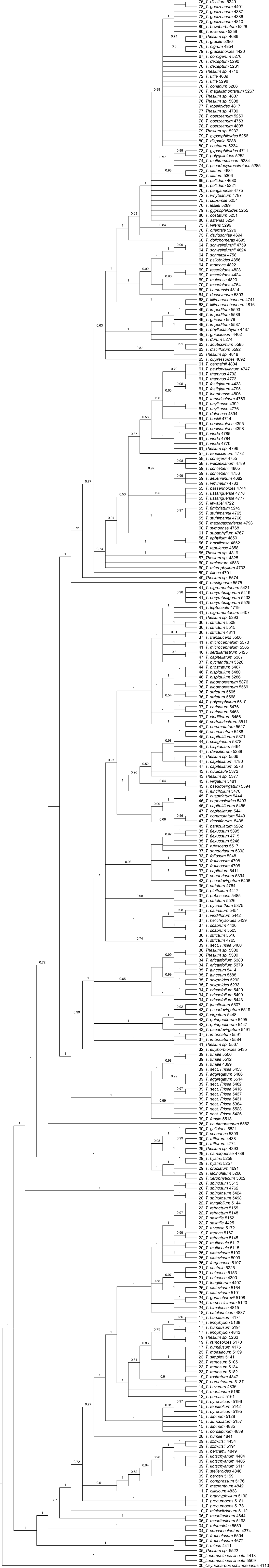
Cladogram resulting from the chloroplast gene partition (concatenated *trnLF* and *trnDT* datasets), analyzed with Bayesian Inference. Numbers at nodes indicate posterior probabilities.

**Fig. S5.**
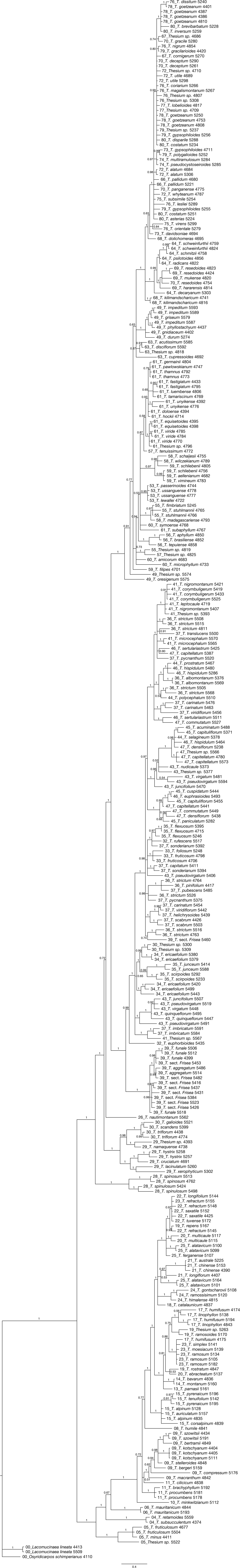
Phylogram resulting from the chloroplast gene partition (concatenated *trnLF* and *trnDT* datasets), analyzed with Bayesian Inference. Numbers at nodes indicate posterior probabilities.

**Fig. S6.**
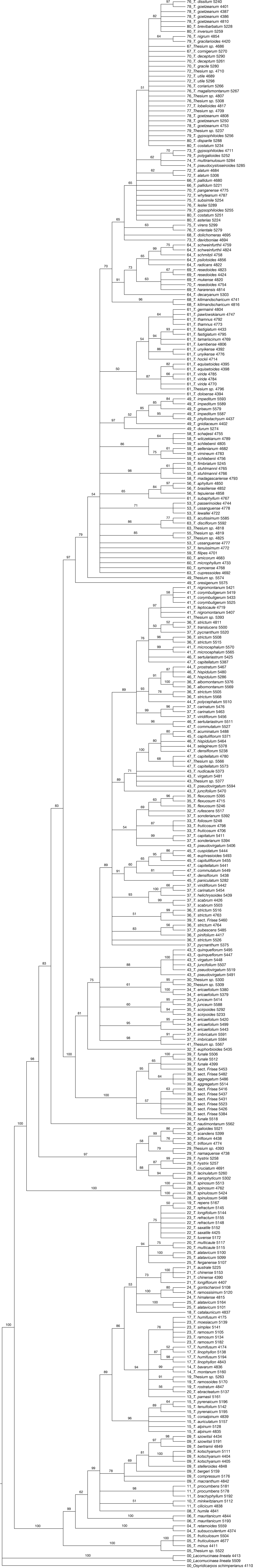
Cladogram resulting from the chloroplast gene partition (concatenated *trnLF* and *trnDT* datasets), analyzed with Maximum Parsimony. Numbers at nodes indicate bootstrap values (1000 replicates).

**Fig. S7.**
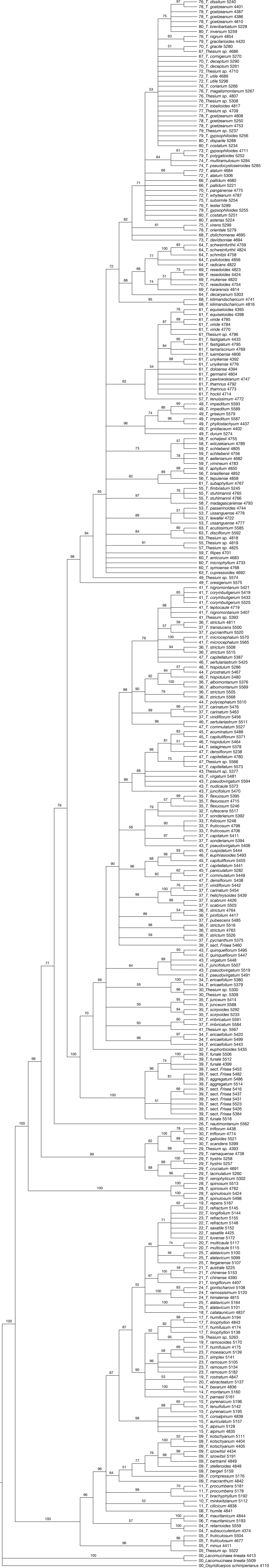
Cladogram resulting from the chloroplast gene partition (concatenated *trnLF* and *trnDT* datasets), analyzed with Maximum Likelihood (GARLI). Numbers at nodes indicate bootstrap values (250 replications).

**Fig. S8.**
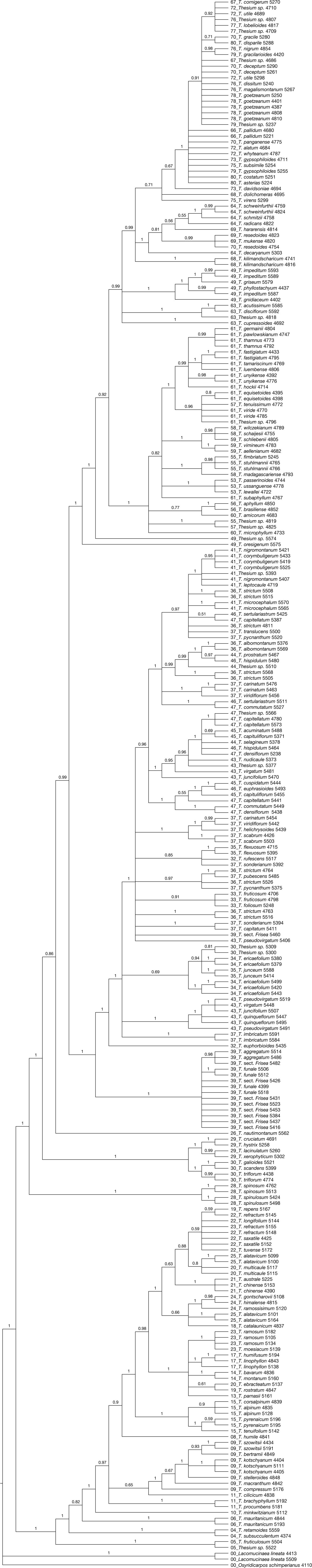
Cladogram resulting from the *trnDT* dataset, analyzed with Bayesian Inference. Numbers at nodes indicate posterior probabilities.

**Fig. S9.**
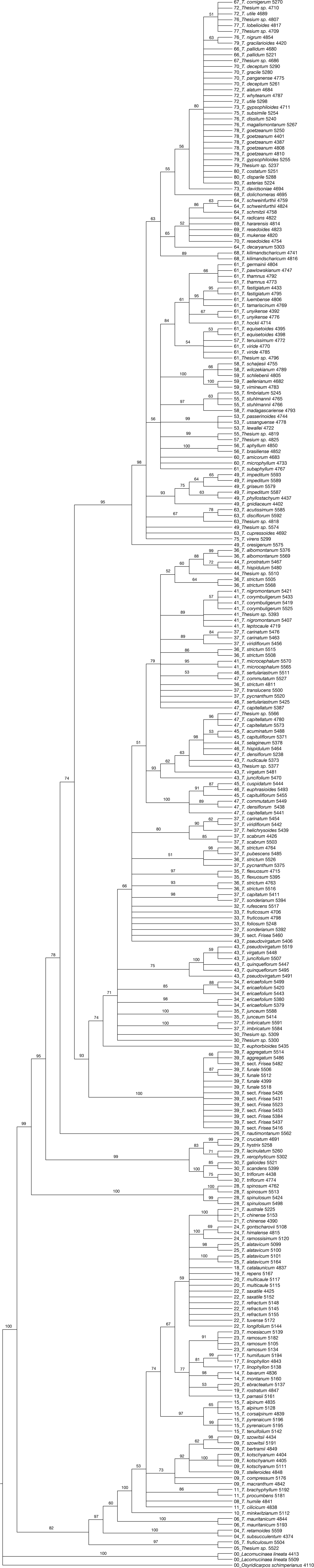
Cladogram resulting from the *trnDT* dataset, analyzed with Maximum Parsimony. Numbers at nodes indicate bootstrap values (1000 replicates).

**Fig. S10.**
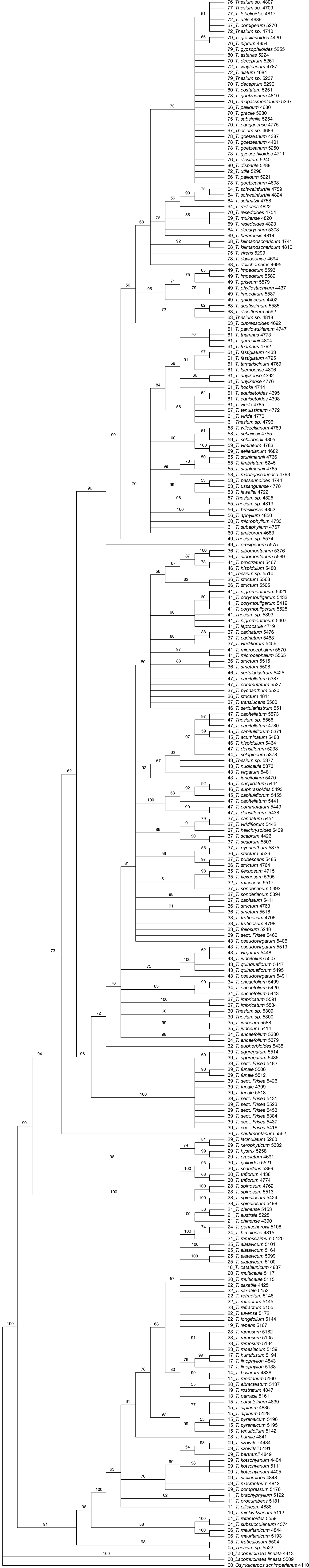
Cladogram resulting from the *trnDT* dataset, analyzed with Maximum Likelihood (GARLI). Numbers at nodes indicate bootstrap values (250 replications).

**Fig. S11.**
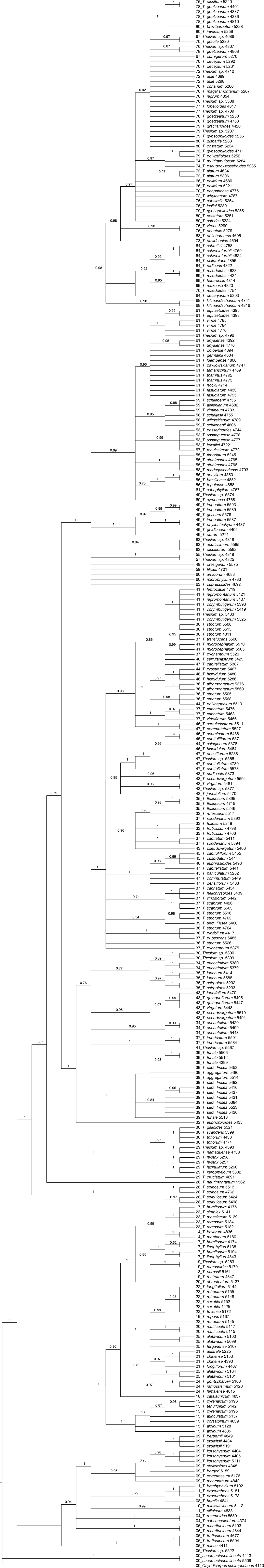
Cladogram resulting from the *trnLF* dataset, analyzed with Bayesian Inference. Numbers at nodes indicate posterior probabilities.

**Fig. S12.**
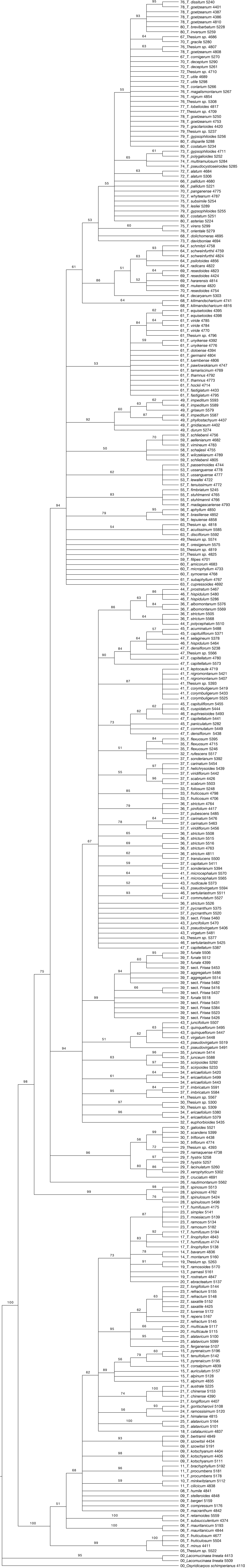
Cladogram resulting from the *trnLF* dataset, analyzed with Maximum Parsimony. Numbers at nodes indicate bootstrap values (1000 replicates).

**Fig. S13.**
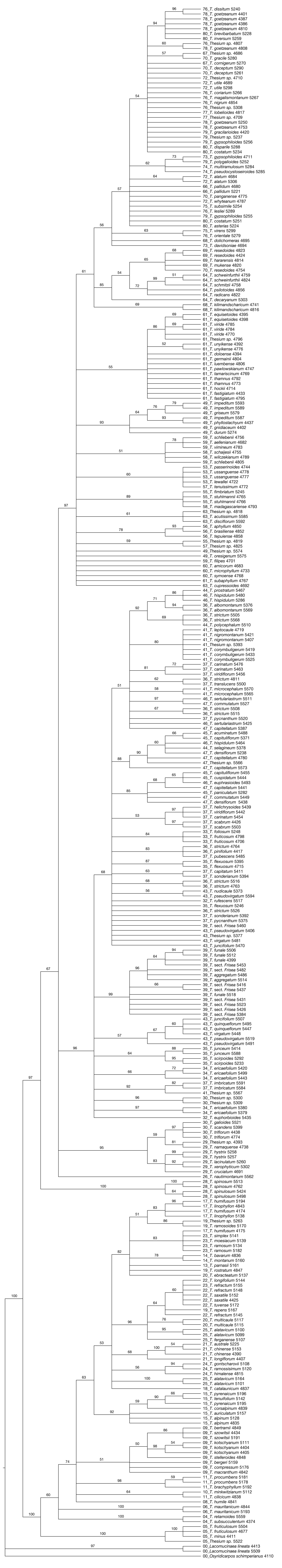
Cladogram resulting from the *trnLF* dataset, analyzed with Maximum Likelihood (GARLI). Numbers at nodes indicate bootstrap values (250 replications).

